# Rubisco proton production can drive the elevation of CO_2_ within condensates and carboxysomes

**DOI:** 10.1101/2020.07.08.125609

**Authors:** Benedict M. Long, Britta Förster, Sacha B. Pulsford, G. Dean Price, Murray R. Badger

## Abstract

Membraneless organelles containing the enzyme Ribulose-1,5-bisphosphate carboxylase/oxygenase (Rubisco) are a common feature of organisms utilizing CO_2_ concentrating mechanisms (CCMs) to enhance photosynthetic carbon acquisition. In cyanobacteria and proteobacteria, the Rubisco condensate is encapsulated in a proteinaceous shell, collectively termed a carboxysome, while some algae and hornworts have evolved Rubisco condensates known as pyrenoids. In both cases, CO_2_ fixation is enhanced compared with the free enzyme. Previous mathematical models have attributed the improved function of carboxysomes to the generation of elevated CO_2_ within the organelle via a co-localized carbonic anhydrase (CA), and inwardly diffusing HCO_3_^-^which has accumulated in the cytoplasm via dedicated transporters. Here we present a novel concept in which we consider the net of two protons produced in every Rubisco carboxylase reaction. We evaluate this in a reaction-diffusion, compartment model to investigate functional advantages these protons may provide Rubisco condensates and carboxysomes, prior to the evolution of HCO3^-^accumulation. Our model highlights that diffusional resistance to reaction species within a condensate allows Rubisco-derived protons to drive the conversion of HCO_3_^-^to CO_2_ via co-localized CA, enhancing both condensate [CO_2_] and Rubisco rate. Protonation of Rubisco substrate (RuBP) and product (PGA) plays an important role in modulating internal pH and CO_2_ generation. Application of the model to putative evolutionary ancestors, prior to contemporary cellular HCO_3_^-^accumulation, revealed photosynthetic enhancements along a logical sequence of advancements, via Rubisco condensation, to fully-formed carboxysomes. Our model suggests that evolution of Rubisco condensation could be favored under low CO_2_ and low light environments.

## INTRODUCTION

Carbon dioxide (CO_2_) fixation into the biosphere has been primarily dependent upon action of the enzyme ribulose-1,5-bisphosphate carboxylase/oxygenase (Rubisco) over geological timescales. Rubisco is distinguished by the competitive inhibition of its carboxylation activity by the alternative substrate molecular oxygen (O_2_), leading to loss of CO_2_ and metabolic energy via a photorespiratory pathway in most phototrophs (1). Almost certainly the most abundant enzyme on the planet (2), Rubisco’s competing catalytic activities have required evolution of the enzyme, and/or its associated machinery, to maintain capture of sufficient carbon into organic molecules to drive life on Earth. In concert with geological weathering, the evolution of oxygenic photosynthesis ≈ 2.4 billion years ago has transformed the atmosphere from one rich in CO_2_ and low in O_2_ to one in which the relative abundances of these gases has overturned (3). Under these conditions, the Rubisco oxygenation reaction has increased, to the detriment of CO_2_ capture. This catalytic paradox has led to different adaptive solutions to ensure effective rates of photosynthetic CO_2_ fixation including; the evolution of the kinetic properties of the enzyme (4), increases in Rubisco abundance in the leaves of many terrestrial C_3_ plants (5) and, the evolution of diverse and complex CO_2_ concentrating mechanisms (CCMs) in many cyanobacteria, algae, and more recently hornworts, CAM, and C_4_ plants (6, 7).

A defining characteristic of contemporary cyanobacteria is the encapsulation of their Rubisco enzymes within specialized, protein-encased micro-compartments called carboxysomes (8). These microbodies are central to the cyanobacterial CCM, in which cellular bicarbonate (HCO_3_ ^-^) is elevated by a combination of membrane-associated HCO_3_ ^-^ pumps and CO_2_ -to-HCO_3_ ^-^ converting complexes (9-11), to drive CO_2_ production within the carboxysome by an internal carbonic anhydrase (CA; 12, 13). This process results in enhanced CO_2_-fixation, with a concomitant decrease in oxygenation, and is a proposed evolutionary adaptation to a low CO_2_ atmosphere (14, 15).

An analogous CCM operates in many algal and hornwort species which contain chloroplastic Rubisco condensates called pyrenoids (16, 17). Pyrenoids are liquid-liquid phase separated Rubisco aggregates which lack the protein shell of a carboxysome (18). These CCMs accumulate HCO_3_^-^and convert it to CO_2_ within the pyrenoid to maximize CO_2_-fixation. Common to cyanobacterial and algal systems is the presence of unique Rubisco-binding proteins, enabling condensation of Rubisco from the bulk cytoplasm (18-25). Condensation of proteins to form aggregates within the cell is an increasingly recognized as a means by which cellular processes can be segregated and organized, across a broad range of biological systems (26-28). The commonality of pyrenoid and carboxysome function (29) despite their disparate evolutionary histories (6), suggest a convergence of function driven by Rubisco condensation. In addition, dependency of functional CCMs on their pyrenoids or carboxysomes (30, 31) has led to the speculation that the evolution of Rubisco organization into membraneless organelles likely preceded systems which enabled elevated cellular HCO_3_^-^(14), raising the possibility that Rubisco condensation and encapsulation may have been the first steps in modern aquatic CCM evolution.

We consider here that, in a primordial model system without active HCO_3_ ^-^ accumulation, co-condensation of Rubisco and CA enzymes is beneficial for the elevation of internal CO_2_ because Rubisco carboxylation produces a net of two protons for every reaction turnover (*SI Appendix*, Fig. S1; 32, 33). These protons can be used within the condensate to convert HCO_3_ ^-^ to CO_2_, with pH lowered and CO_2_ elevated as a result of restricted outward diffusion due to the high concentration of protein in the condensate and surrounding cell matrix acting as a barrier to diffusion. We propose that proton release within a primordial Rubisco condensate enabled the evolution of carboxysomes with enhanced carboxylation rates, prior to advancements which enabled cellular HCO_3_ ^-^ accumulation.

## RESULTS

### The modelling of free Rubisco and Rubisco compartments

To demonstrate the feasibility of our proposal we initially consider a model of free Rubisco, a Rubisco condensate, and a carboxysome based on a set of compartmentalized reactions described in Fig. 1 and associated tables of parameters (Table 1, Table 2, *SI Appendix, SI Methods*). We present data for the tobacco Rubisco enzyme as an exemplar, noting that evaluations of other Rubisco enzymes in the model (Table 2) provide comparative outcomes. We assume a system with fixed external inorganic carbon (C_i_; HCO_3_^-^and CO_2_) supply in the absence of a functional CCM, simulating a primordial evolutionary state prior to the development of HCO_3_ ^-^ accumulation in unicellular photosynthetic organisms (14).

**Table 1.**
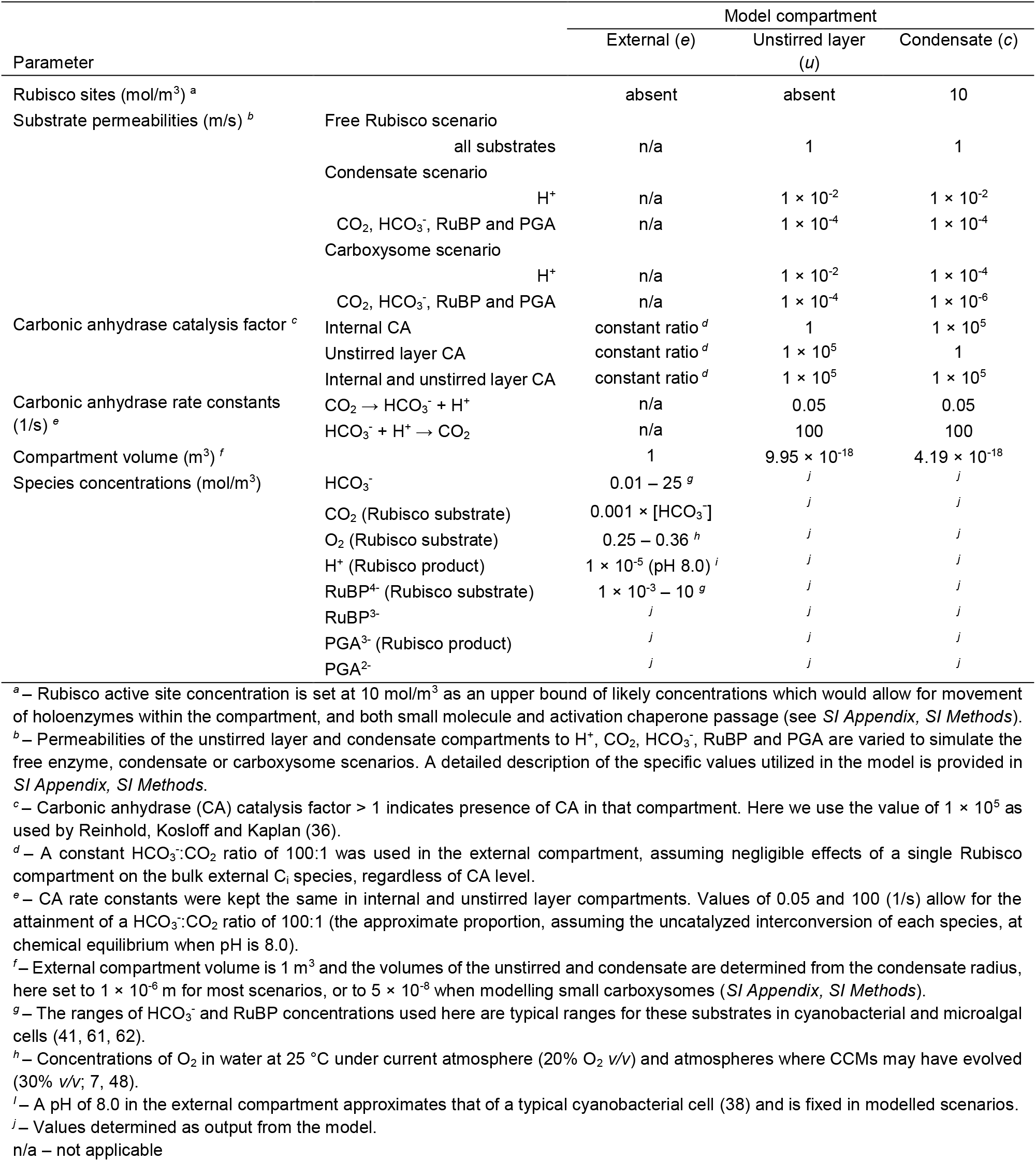
Typical initial values used in the COPASI biochemical compartment model in this study.

**Table 2.**
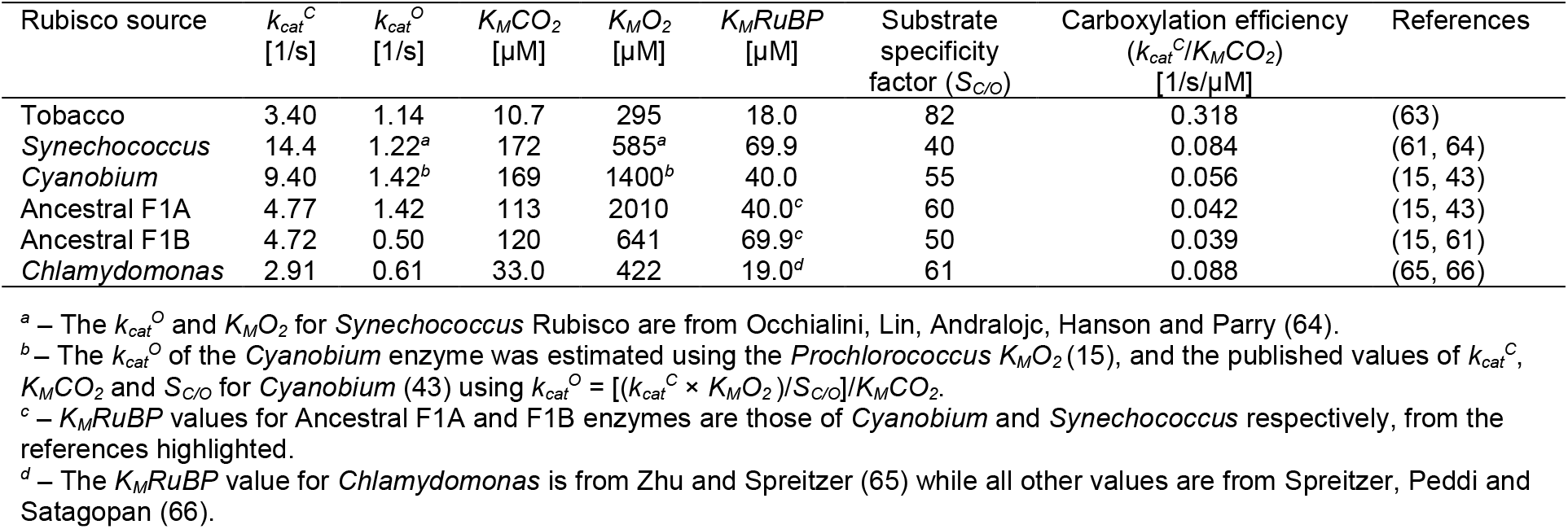
Rubisco catalytic parameters used in competition modelling.

**Fig. 1.**
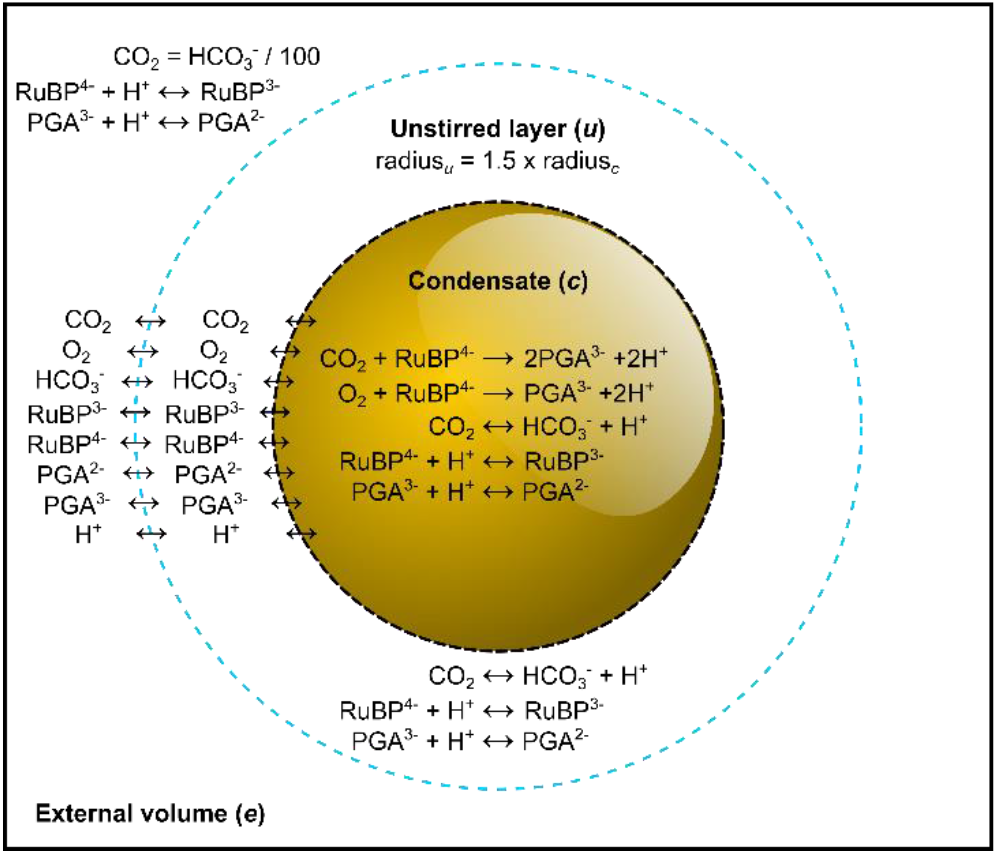
Rubisco compartment model. A visual description of the compartment model used in this study. The model consists of three reaction compartments. The external compartment (*e*) is analogous to a static cellular cytoplasm in which we set the concentration of inorganic carbon (C_i_) species (CO_2_ and HCO_3_ ^-^), along with RuBP and PGA which can undergo reversible reactions with protons (H^+^). Interconversion of C_i_ species in the unstirred (*u*) and condensate (*c*) compartments is catalyzed by carbonic anhydrase, whereas RuBP and PGA protonation/deprotonation is determined by the rate of conversion at physiological pH given *pKa* values of relevant functional groups (*SI Appendix*, Fig. S1). The central compartment of the model is a Rubisco condensate in which Rubisco carboxylation and oxygenation reactions occur, along with RuBP/PGA protonation and CA reactions. In modelling scenarios, we modify external CA by modulating its function in the unstirred layer. The diffusion of all reaction species between each compartment can be set in the model to simulate either a free Rubisco enzyme, a Rubisco condensate, or a carboxysome as described in Table 1. Model parameterization is described in detail in *SI Appendix, SI Methods*.

Our model consists of three nested compartments with a specialized Rubisco compartment (which can be described as a Rubisco condensate or carboxysome by modifying the compartment boundary permeabilities) at the center (Fig. 1). This compartment is surrounded by an unstirred boundary layer which we assume has diffusive resistance to substrate movement, and is bounded by an external compartment, at pH 8.0, which supplies reaction substrates. We contain Rubisco reactions within the central compartment but allow the protonation and deprotonation of reaction species (ribulose-1,5-bisphosphate [RuBP] and phosphoglycerate [PGA]) to occur in all compartments. We include the competing Rubisco substrates, O_2_ and CO_2_, and enable the latter to be interconverted with the more abundant HCO_3_ ^-^ species through pH control and the interaction of carbonic anhydrase (CA), whose position in the model we manipulate. Specific details of the model and its parameterization are provided in Table 1, Methods, and *SI Appendix, SI Methods*.

Previous models consider the function of carboxysomes, for example, within cells capable of active accumulation of HCO_3_ ^-^ in chemical disequilibrium with CO_2_, and apply diffusional resistances to Rubisco reactants and products within a modelled reaction compartment (9, 34-42). We also apply diffusional resistances to all substrates in our model, but consider cytoplasmic CO_2_ and HCO_3_ ^-^ supply to be in chemical equilibrium, as would occur in the absence of an active CCM, in order to address any beneficial role of Rubisco compartmentation alone. The novel aspects of this model are; a chemical equilibrium of CO_2_ and HCO_3_ ^-^ in the compartment surrounding Rubisco, the inclusion of proton production by the Rubisco carboxylation and oxygenation reactions (and their equilibration across the carboxysome shell by diffusion) and, proton movement via protonated ribulose-1,5-bishposphate (RuBP) and phosphoglycerate (PGA) species. Application of the model to existing experimental data (*SI Appendix*, Fig. S2) provides a good estimation of the differential function of both the free Rubisco enzyme and carboxysomes isolated from the cyanobacterium *Cyanobium* (43), thus providing confidence in the model.

### Carboxysome and condensate proton permeability

An important assumption in our model is that there is some resistance to substrate movement across Rubisco compartment boundaries, including protons. The diffusion of protons across the carboxysome envelope has been considered previously (44). but within the context that pH stabilization is entirely dependent upon free diffusion through the shell, and in the absence of Rubisco activity which could lead to internally produced protons. In that study, pH-dependent fluorescent protein inside the carboxysome responded within millisecond time-scales to changes in the external pH, resulting in the conclusion that protons entered or exited the carboxysome freely. However, this result is also consistent with some level of diffusional resistance to protons, since considerable restriction to proton permeability can yield internal pH equilibration within even faster time-frames (*SI Appendix*, Fig. S3). Indeed, these previous findings have been shown to be consistent with a steady-state *ΔpH* across the carboxysome shell, where the relative rates of internal proton production and leakage across the shell can maintain an acidic interior (38). We therefore assume permeabilities to protons which are consistent with existing data, yet enable some restriction on proton movement. Molecular simulations suggest that pores in the carboxysome shell favor negatively charged ions such as HCO_3_ ^-^, RuBP and PGA (45), and it is unlikely that the H_3_O^+^ ion will easily traverse the protein shell. For the diffusion of H_3_O^+^ in water, it is considered to have a higher diffusion rate than other ions in solution due to its participation in a proton wire system in collaboration with water (46).

For modelling purposes here, we have assumed that negatively charged ions have a permeability of 10^-6^ m/s across the carboxysome shell while H_3_O^+^ has a higher value of 10^-4^ m/s (see *SI Appendix, SI Methods*). The data in Fig. 2 show that proton permeability values greater than ≈ 10^-3^ m/s for the carboxysome shell leads to low carboxysome [CO_2_] and lower Rubisco carboxylation turnover (under sub-saturating substrate supply).

**Fig. 2.**
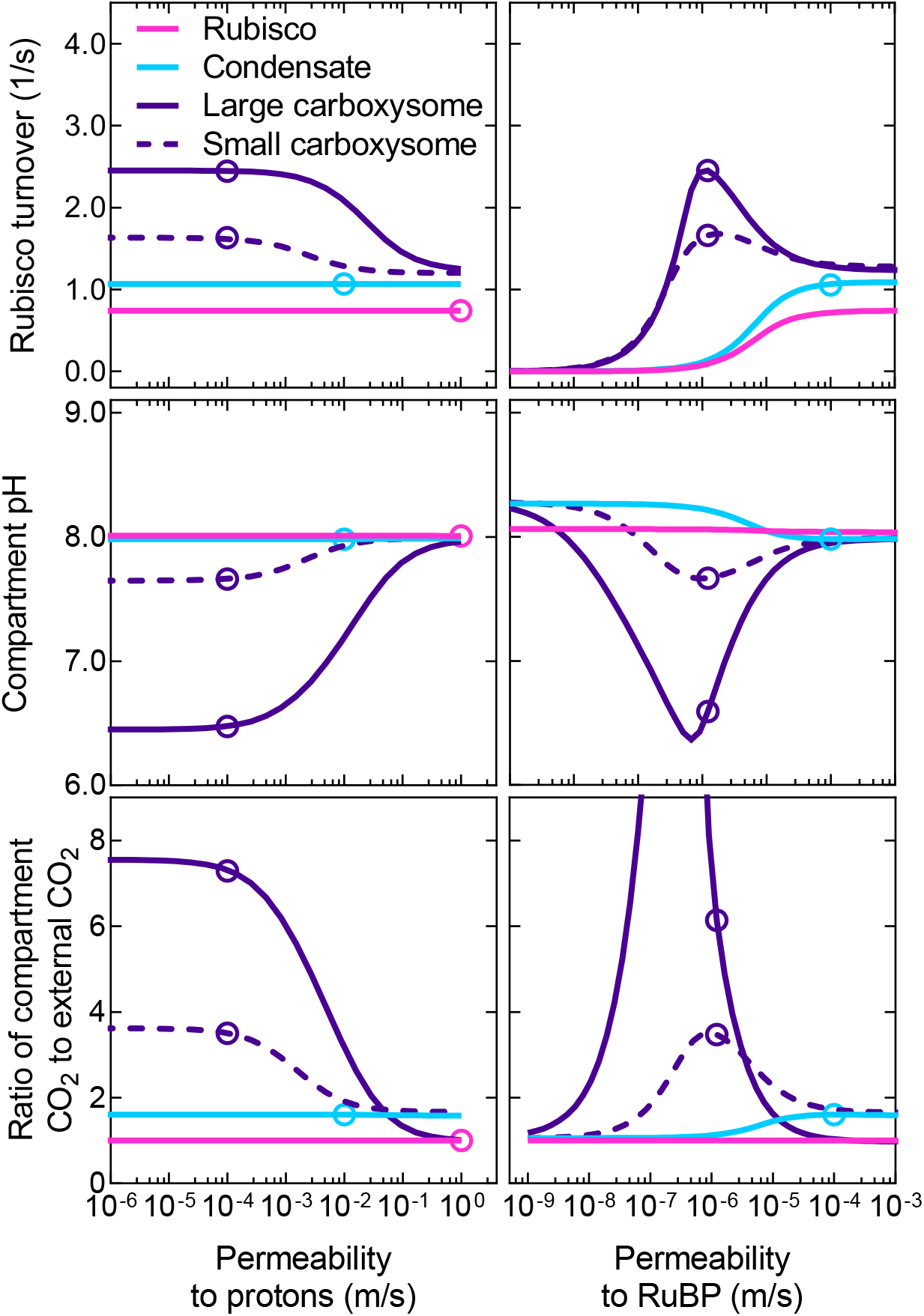
Carboxysome and condensate function are dependent on proton and RuBP permeabilities. Rubisco carboxylation turnover (top panels), compartment pH (middle panels), and the ratio of Rubisco compartment CO_2_ to external CO_2_ (bottom panels) are dependent upon the permeability of the compartment to protons (left panels) and RuBP (right panels). Shown are modelled responses for free Rubisco (*pink lines*), a Rubisco condensate (*blue lines*), a large (1 × 10^-6^ m radius) carboxysome (*purple lines*), and a small (5× 10^-8^ m radius) carboxysome (*purple dashed lines*) at sub-saturating substrate concentrations (1 mM HCO_3_ ^-^ [0.01 mM CO_2_ ] and either 35, 50, 87, or 1,300 µM RuBP for the free enzyme, condensate, small carboxysome and large carboxysome respectively; see Fig. 3 and *SI Appendix*, Fig. S4). *Open circles* represent the values obtained for typical permeabilities used in the model (Table 1). Data were generated using the COPASI (60) model run in parameter scan mode, achieving steady-state values over the range of proton and RuBP permeabilities indicated for the Rubisco compartment. For all cases CA activity was only present within the Rubisco compartment. Data presented are for the tobacco Rubisco with parameters listed in Table 2.

Unlike the carboxysome, Rubisco condensate proton permeability does not appear to affect Rubisco carboxylation in our model under sub-saturating substrate conditions (Fig. 2). However, the condensate [CO_2_] does appear to correlate with modelled changes in internal pH, suggesting a role for protons in determining condensate [CO_2_] and carboxylation rates (Fig. S4). In the case of a condensate, RuBP^3-^ is able to carry protons from outside to inside, and therefore provides protons required to convert HCO_3_ ^-^ to CO_2_. This can be observed within the model by varying RuBP permeability, with values above 10^-6^ m/s leading to increased compartment [CO_2_], and enhanced carboxylation (Fig. 2). This value is consistent with application of the model to experimental data for carboxysomes (Fig. S2). RuBP permeabilities below 10^-6^ m/s also leads to rate-limiting concentrations of RuBP in all compartment types under sub-saturating substrate supply (Fig. 2). Variation of condensate permeabilities to protons and RuBP shows that fluxes of reaction species across the condensate boundary are permeability-dependent (Fig. S5).

### Protons derived from Rubisco reactions influence both condensate and carboxysome function

The influence of Rubisco proton production on the response of carboxylation rate and Rubisco compartment [CO_2_] to external substrate supply (RuBP and HCO_3_ ^-^) is considered in Fig. 3. For the purpose of demonstrating the role of protons, we assess model responses under sub-saturating substrate conditions where we observe greatest proton responses within the model (*SI Appendix*, Fig. S4), and account for differential changes in the ‘apparent’ *K*_*M*_*RuBP* arising from an assumed decreased permeability to this substrate in condensates and carboxysomes (Fig. 3A; *SI Appendix*, Fig. S6).

**Fig. 3.**
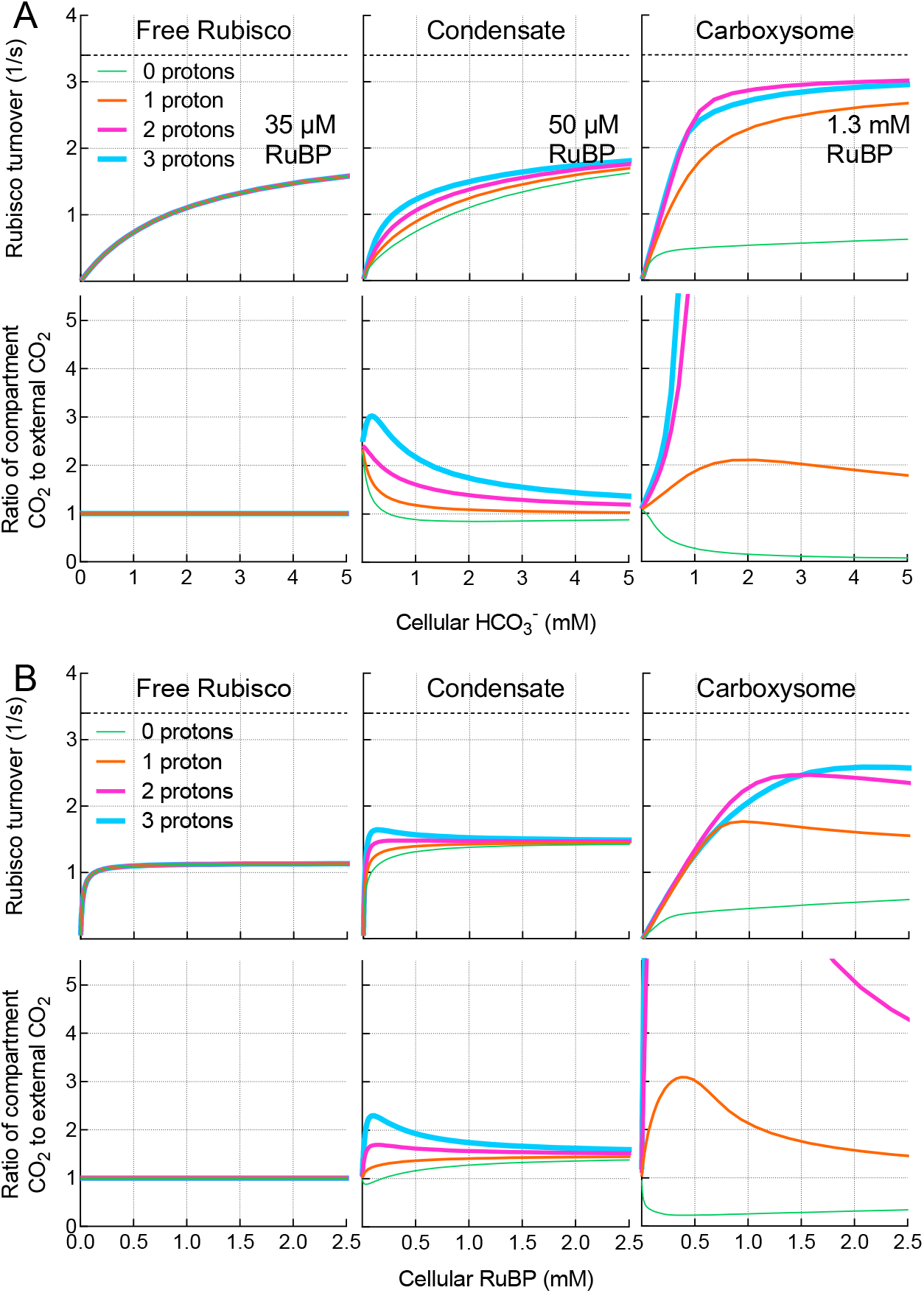
Modelled responses of free Rubisco, condensate and carboxysome to external substrate supply. Modelled responses of free Rubisco, condensate and carboxysome to external HCO_3_ ^-^ (A) and RuBP (B) supply. Carboxylation turnover rates (top panels) and the ratio of Rubisco compartment CO_2_ to external CO_2_ (lower panels) for the free enzyme, condensate and carboxysomes with zero, one, two or three protons being produced as products of the carboxylation reaction at sub-saturating RuBP (A) or HCO_3_ ^-^ (B). Variation in permeabilities to RuBP between the free enzyme, condensate and carboxysome within the model result in increases in the apparent *KMRuBP* as permeabilities decline (*SI Appendix*, Fig. S6). For HCO_3_ ^-^ responses, the RuBP concentration used in each scenario is indicated. For RuBP responses, the HCO_3_ ^-^ supply set at 1 mM [0.01 mM CO_2_ ]). External CO_2_ is set to 1/100 × external HCO_3_ ^-^. Net proton production by Rubisco carboxylation is theoretically two protons (Fig. 1). In all cases modelled rates include carbonic anhydrase (CA) activity only in the Rubisco compartment. Rubisco turnover in a carboxysome producing three protons per carboxylation drops below that of the two-proton system due a proton-driven shift in the RuBP^3-^:RuBP^4-^ ratio, decreasing the concentration of the Rubisco substrate species, RuBP^4-^. Maximum carboxylation turnover rate (*k*_cat_^*C*^; 3.4 [1/s]) of the tobacco Rubisco used in this example (Table 2), is indicated by the *dashed line* in the upper panels. The COPASI (60) model was run in parameter scan mode, achieving steady-state values over the range of HCO_3_ ^-^ concentrations indicated. Proton number was manufactured by modifying the Rubisco carboxylation reaction stoichiometry to produce zero, one, two or three protons in the model.

If we eliminate carboxylation-derived proton production within a Rubisco compartment in the model (zero protons; Fig. 3A) and assume diffusional limitations to proton movement, then proton-driven, CA-dependent CO_2_ production within either a condensate or carboxysome becomes limited by the influx of protons from the external environment. With increasing compartment proton production per Rubisco carboxylation reaction (one and two protons; Fig. 3A), the lower pH (and increased [H^+^] for CA-driven HCO_3_^-^dehydration) enhances CO_2_ concentration within a condensate, and even more so within a carboxysome, increasing Rubisco CO_2_ fixation rates. Levels of CO_2_ are further enhanced if more protons are able to be produced per Rubisco turnover (e.g. three protons in Fig. 3A). In contrast to a condensate or carboxysome, the free Rubisco enzyme is unaffected by proton production due to the absence of a compartment with associated diffusional limitations.

We also assessed the modelled responses of the free enzyme, its condensate, and carboxysomes to [RuBP] under sub-saturating HCO_3_^-^(1 mM; Fig. 3B). Again, proton production by Rubisco led to increases in condensate and carboxysome [CO_2_], despite relatively high external HCO_3_^-^supply, resulting in corresponding increases in Rubisco carboxylation rate.

In considering why these results are obtained, it is important to note that at pH 8.0, which we set for the bulk medium in our model, free [H^+^] is only 10 nM. In the Rubisco compartment volume utilized in our model (≈ 5 x 10^-18^ m^3^) this represents less than one proton, and means that the diffusional driving forces for proton exchange across the carboxysomal shell, for example, are 10^4^–10^6^ -fold smaller than those driving CO_2_, HCO_3_^-^, PGA and RuBP diffusion (which are in µM and mM ranges). Hence, inward H^+^ diffusion will be rate-limiting depending on the permeability of the compartment to protons. Therefore, other proton sources must provide substrate for the CA/HCO_3_ ^-^ dehydration reaction. The net outcome is that proton production by Rubisco carboxylation within a diffusion-limited compartment leads to decreased pH, elevated CO_2_, and improvement in carboxylation turnover, compared with the free enzyme (*SI Appendix*, Fig. S7).

The model also shows an increase in carboxylation turnover resulting from the protons arising through oxygenation within a carboxysome, although this appears negligible in a Rubisco condensate (*SI Appendix*, Fig. S8). Notably, like the model of Mangan, Flamholz, Hood, Milo and Savage (38), we find that carboxysome function does not require specific diffusional limitation to O_2_ influx in order to reduce oxygenation, due to competitive inhibition by the increase in CO_2_.

### Roles for RuBP, PGA and CO_2_ in Rubisco compartment pH

When we run the full model, including net proton production by Rubisco, carboxysome pH becomes acidic at limiting substrate concentrations as a result of limited proton efflux (Fig. 4). As external substrate increases and internal CO_2_ rises, the pH approaches that of the bulk external medium (which we set at pH 8.0 with the H^+^ concentration). Here, increasing CO_2_ efflux from the carboxysome is able to effectively dissipate protons from the Rubisco compartment, since there is a net loss of CO_2_ that would otherwise be used for proton production by the CA hydration reaction (CO_2_ + H_2_O ↔ HCO_3_ ^-^ + H^+^). This can be seen when we modify the rate of CO_2_ efflux from the carboxysome by altering compartment CO_2_ permeability. Slow CO_2_ efflux leads to increased free proton concentrations within the compartment, and fast efflux enables a return to approximately external pH as substrate supply increases (Fig. 4).

**Fig. 4.**
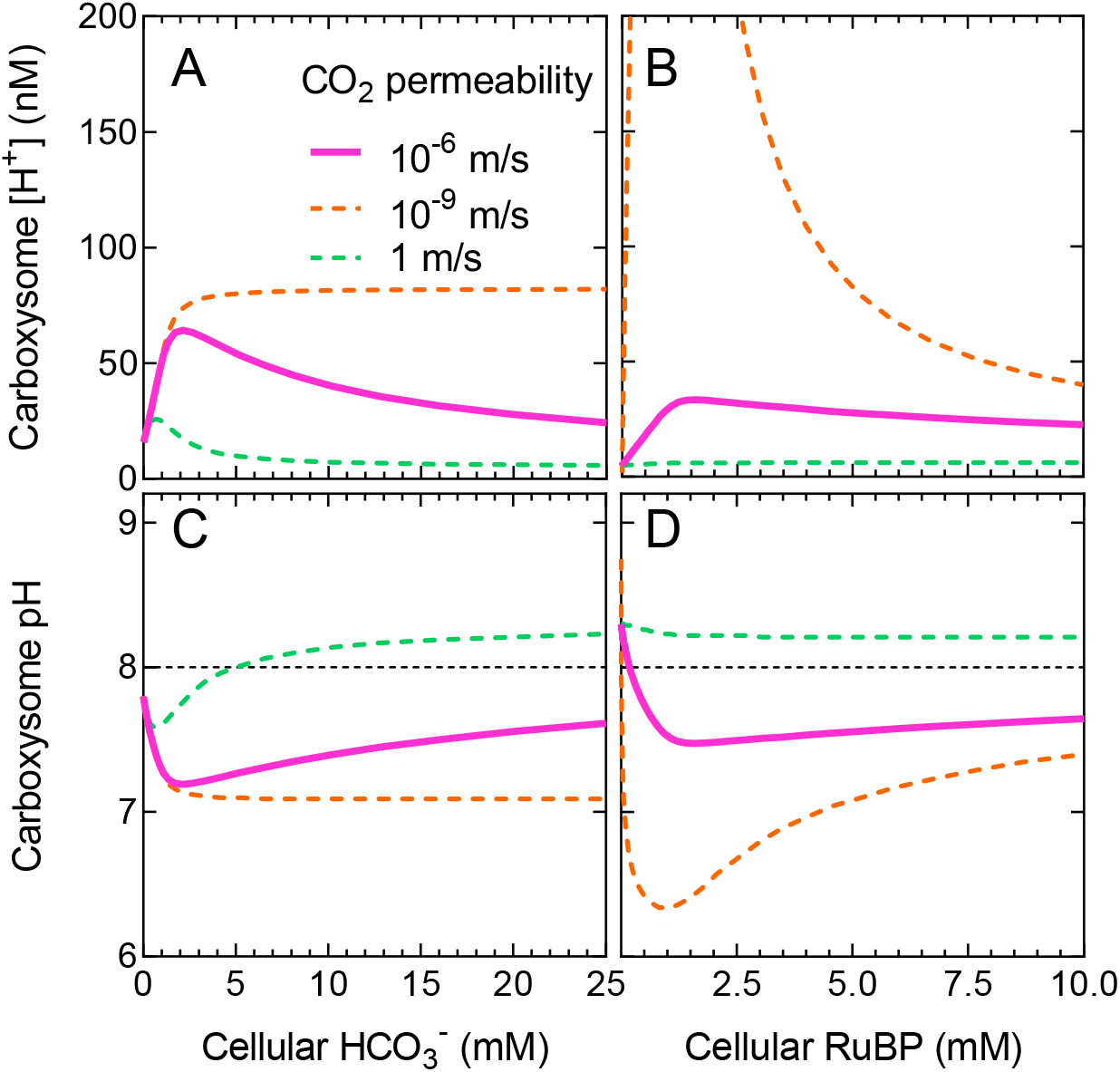
Carboxysome proton concentration is modulated by CO_2_ efflux. Carboxysome free proton concentration (A, B) and carboxysome pH (C, D) indicate that a functional carboxysome compartment undergoes net acidification at limiting HCO_3_ ^-^ and RuBP supply (*SI Appendix*, Fig. S4). Plotted are proton concentration and pH over a range of [HCO_3_ ^-^] and [RuBP] for modelled carboxysomes with an internal carbonic anhydrase (CA), allowing for two protons to be produced per carboxylation reaction, and under typical modelled CO_2_ permeability within the model (10^-6^ m/s; *solid pink lines*). If CO_2_ efflux were rapid and unimpeded (CO_2_ permeability 1 m/s; *dashed green lines*), pH rapidly returns to ≈ 8 (*black dashed line*, panels C and D) as external limiting substrate supply increases. Slow CO_2_ efflux (CO_2_ permeability 10^-9^ m/s; *dashed orange lines*) does not allow for dissipation of protons. Efflux of CO_2_ from the carboxysome contributes to pH maintenance is it represents the loss of substrate for the CA hydration reaction (CO_2_ + H_2_O ↔ HCO_3_ ^-^ + H^+^), which would otherwise lead to free proton release. Each dataset was modelled with an initial [RuBP] of 5 mM for HCO_3_ ^-^response curves, and 20 mM HCO_3_ ^-^ in the case of RuBP response curves and CA activity is confined only to the carboxysome compartment. All other permeabilities under these conditions are set to 10^-6^ m/s as for a carboxysome (Table 1). The COPASI (60) model was run in parameter scan mode, achieving steady-state values over a range of substrate concentrations. Data presented are for the tobacco Rubisco (Table 2).

When we consider reaction species’ fluxes across the Rubisco compartment boundary, we find that both RuBP^3-^ and PGA^2-^ can play a role in carrying protons out of both condensates and carboxysomes (Fig. 5, *SI Appendix*, Fig. S9), contributing to the stabilization of internal pH. In our model of carboxysomes, RuBP^3-^ and PGA^2-^ efflux plays a significant role in pH balance at very low HCO_3_ ^-^. However, as external HCO_3_ ^-^ rises, carboxysome CO_2_ efflux replaces RuBP and PGA as the major proton carrier (Fig. 5) as described above. For a condensate, CO_2_ efflux is the primary means of stabilizing internal pH within the model over a range of HCO_3_ ^-^ concentrations (*SI Appendix*, Fig. S9).

**Fig. 5.**
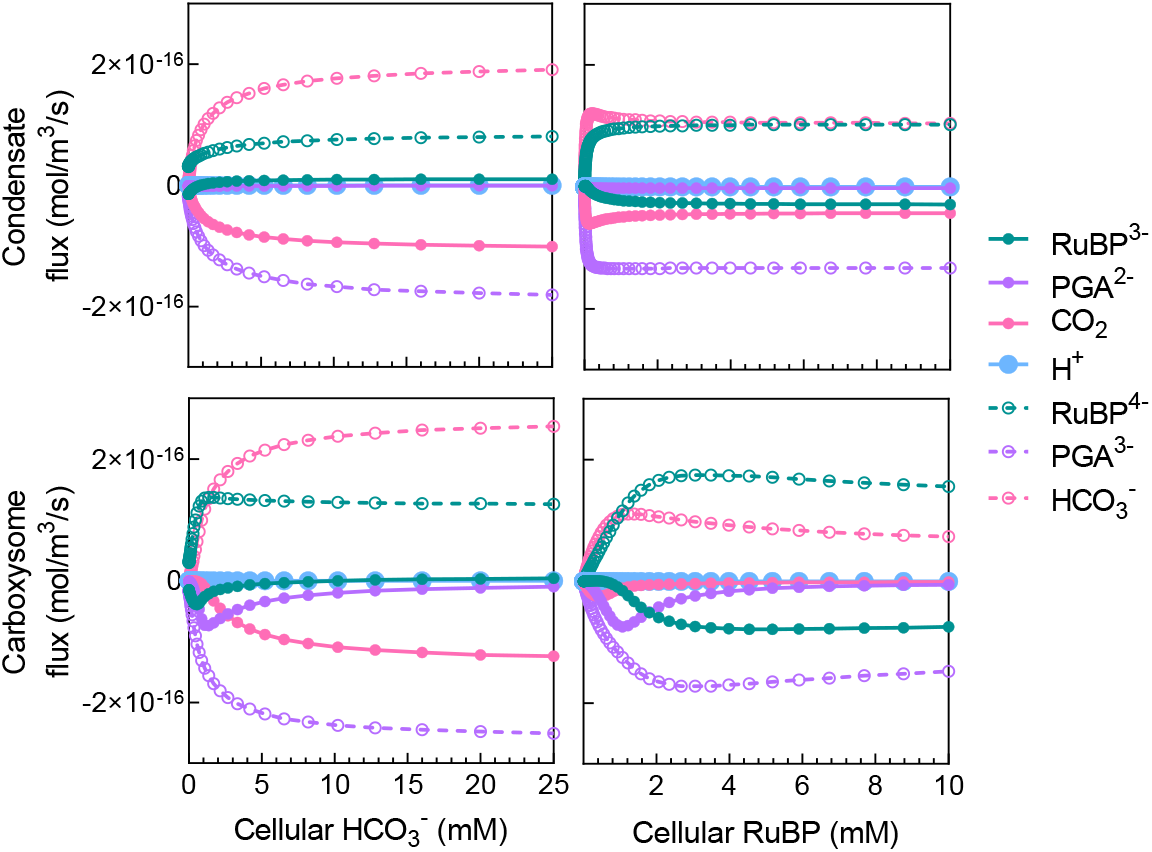
Proton carriers help maintain compartment pH. Diffusional flux of chemical species across the condensate/carboxysome boundary over a range of HCO_3_ ^-^ and RuBP concentrations in the model. Protons are carried by RuBP^3-^, PGA^2-^ and CO_2_ (as the substrate required for free proton release via the CA hydration reaction). Net free H^+^ fluxes are extremely small and contributions to internal pH primarily arise through net fluxes of proton-carrier substrates (*solid circles*). The deprotonated RuBP^4-^ and PGA^3-^ are the substrate and product of Rubisco carboxylation, respectively, within the model. Positive flux values indicate net influx into the compartment and negative values indicate net efflux. The COPASI (60) model was run in parameter scan mode, achieving steady-state at each substrate concentration. For HCO_3_ ^-^ response curves, RuBP was set to 50 µM for a condensate and 1.3 mM for a carboxysome based on changes in apparent *KMRuBP* values arising from diffusional resistance (Fig. 3, *SI Appendix*, Fig. S6). External [HCO_3_ ^-^] was set to 1 mM for the generation of RuBP response curves. Data presented are for the tobacco Rubisco (Table 2) and model parameters for a condensate or a carboxysome are indicated in Table 1. These data are summarized in *SI Appendix*, Fig. S9.

Similar responses can be seen over a range of RuBP concentrations, where CO_2_ efflux also plays a dominant role as a proton efflux carrier in the condensate, while RuBP^3-^ and PGA^2-^ efflux are the major contributors to shuttling protons out of the carboxysome (Fig. 5, *SI Appendix*, Fig. S9). These results highlight that the inclusion of RuBP and PGA as proton carriers is essential in describing the functioning of the carboxysome, as they contribute to maintaining internal pH and, therefore, pH-sensitive Rubisco activity (47).

Model output also emphasizes that compartment pH is highly dependent on the buffering capacity of RuBP and PGA. Not only does RuBP carry protons released in Rubisco reactions, both species also undergo protonation and deprotonation at physiological pH. Modifying their *pKa* values *in silico* significantly alters Rubisco compartment pH (*SI Appendix*, Fig. S10).

### The need for CA

In previous models of carboxysome function there is a need for internal CA to accelerate the conversion of HCO_3_ ^-^ to CO_2_ to support high rates of Rubisco CO_2_ fixation and CO_2_ leakage out of the carboxysome. While those models consider functional CCMs with active cellular HCO_3_ ^-^ accumulation, this need for CA is also true here, with internal interconversion needing acceleration to maximize Rubisco CO_2_ fixation at saturating external HCO_3_ ^-^ (*SI Appendix*, Fig. S11). Additionally, we find that CA inclusion within a condensate at high rates provides additional benefit to Rubisco carboxylation rates (*SI Appendix*, Fig. S11, Fig. S12).

It is also apparent that in the carboxysome CA function is dependent upon RuBP and HCO_3_^-^supply, emphasizing that provision of protons from the Rubisco reaction is essential for the production of CO_2_ from HCO_3_ ^-^ via the CA enzyme (*SI Appendix*, Fig. S13). This dependency appears to be much less in a condensate due to our assumption of much higher permeabilities of RuBP^3-^ and CO_2_ to the condensate interior (*SI Appendix*, Fig. S11) and the low *K*_*M*_*CO*_*2*_ of the tobacco Rubisco modelled in this scenario.

### Carboxysome evolution via Rubisco condensation

Our model shows that Rubisco co-condensed with CA gives improved function over its free enzyme (Fig. 3). Given that Rubisco condensation underpins carboxysome biogenesis (20, 25), we considered that the model may provide insights into carboxysome evolution via intermediate states utilizing Rubisco condensates, prior to HCO_3_ ^-^ accumulation in the cell through inorganic carbon (C_i_) transport.

To investigate this proposal, we analyzed the performance of potential evolutionary intermediates in a series of hypothesized pathways from free Rubisco to carboxysomes, in the absence of functional C_i_ accumulation (assuming this was a later evolutionary progression; 14). We first assumed a photorespiratory loss of ½ mole of CO_2_ for every mole of O_2_ fixed within the model (while carboxylation yields two molecules of PGA from one CO_2_, oxygenation yields only one PGA and one CO_2_ is lost via photorespiration), and by calculating average net carboxylation rates by each hypothesized evolutionary intermediate under different atmospheric conditions (see Methods). This allowed relative fitness comparisons of proposed evolutionary states under what might reasonably be considered low or high atmospheric CO_2_, as may have been experienced under atmospheres where CCMs arose (7, 48). We assume here that a greater average net carboxylation rate for a particular evolutionary intermediate (over a given range of HCO_3_ ^-^) would result in greater relative fitness and, therefore, evolutionary advantage. We apply this concept over HCO_3_ ^-^ ranges within the model, rather than single-point comparisons, to provide a clearer view of Rubisco responses resulting from variations in the observed response curves and, to assess responses under different atmospheric CO_2_ compositions (*SI Appendix*, Fig. S14).

We made these fitness calculations at both 20% and 30% (*v/v*) O_2_, to simulate alternative atmospheres under which CCMs may have arisen (7, 48). We calculated the average net carboxylation rates for the defined HCO_3_ ^-^ ranges (see Methods), and generated phenotype matrices to allow comparison of possible evolutionary states (Fig. 6; *SI Appendix, SI* datasets S1 and S2). In evolutionary state comparisons, those states which lead to larger average net carboxylation rates were considered to have greater fitness.

**Fig. 6.**
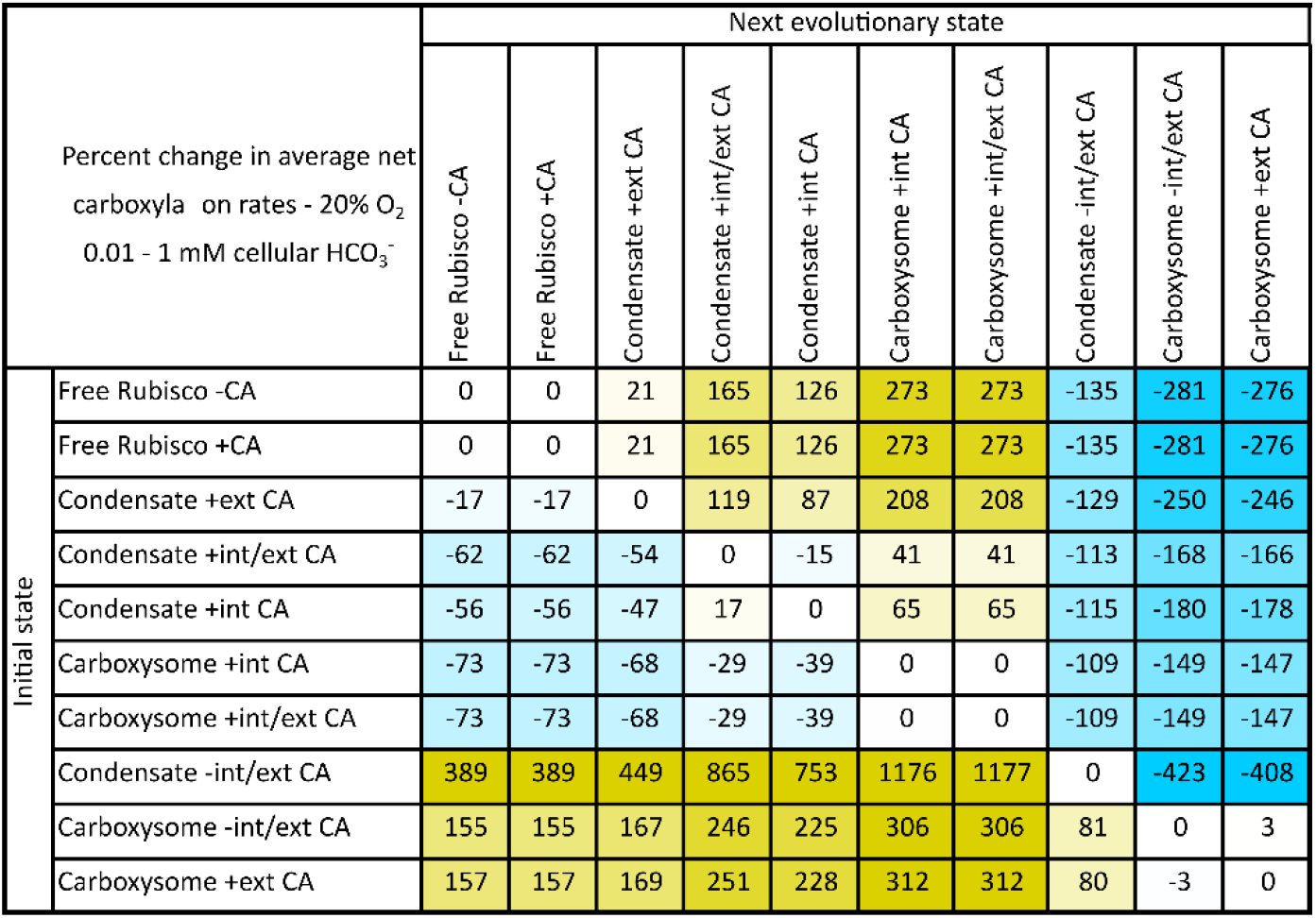
Fitness matrix for proposed evolutionary steps from free Rubisco to contemporary carboxysomes, before the evolution of active C_i_ transport. An example fitness matrix for the tobacco Rubisco enzyme showing percentage difference in average net Rubisco carboxylation rates (see Methods and *SI Appendix*, Fig. S14) over a 0.01 – 1 mM HCO_3_ ^-^ in a 20% (*v/v*) O_2_ atmosphere in systems lacking C_i_ transport and HCO_3_ ^-^ accumulation. Within this table an initial evolutionary state (*left, rows*) can be compared with any potential next evolutionary state (*top, columns*). Values are percent changes in average net carboxylation rates between each state. Positive (*yellow*) values indicate an improvement in average net Rubisco carboxylation turnover when evolving from an initial state to the next evolutionary state. Negative (*blue*) values indicate a net detriment. As an example, evolution from free Rubisco with associated carbonic anhydrase (Free Rubisco + CA) to a Rubisco condensate with an internal CA (Condensate + int CA) shows a 126% improvement in average net carboxylation turnover. We assume that such an evolutionary adaptation would result in a competitive advantage over the initial state. Contrarily, a Rubisco condensate with both internal and external CA (Condensate +int/ext CA) shows a decrease in average net carboxylation turnover of 15% if the external CA was lost in an evolutionary adaptation (Condensate + int CA). Data presented here are for the tobacco Rubisco (Table 2), using the compartment model to simulate all potential evolutionary states (Table 1). External CA is modelled in the unstirred layer. The same pattern of potential evolutionary improvements is apparent regardless of the Rubisco source or carboxysome size used in the model, assuming sufficient RuBP supply (*SI Appendix, SI datasets* S1 and S2).

The relative fitness of Form I Rubisco enzymes (49) from a variety of sources were compared within the model, over all proposed evolutionary intermediates, to observe any differences resulting from varying Rubisco catalytic parameters (Table 2). A complete analysis of each Rubisco source and its performance at each proposed evolutionary step, under varied HCO_3_ ^-^ and O_2_ conditions is supplied in the *SI Appendix, SI* datasets S1 and S2.

Regardless of the type of Rubisco used in this analysis, the same pattern of potential evolutionary augmentations was favored (Fig. 6, Fig. 7, and *SI Appendix, SI* datasets S1 and S2). In all cases, condensation of Rubisco in the absence of a CA enzyme, as an initial evolutionary step (‘Rubisco -CA’ to ‘Condensate – int/ext CA’; Fig. 6), resulted in a decrease in fitness, emphasizing that the starting point for the evolution of Rubisco condensates and carboxysomes likely began with a cellular CA present (‘Rubisco + CA’). Again, we emphasize here that our modelling assumes no active C_i_ accumulation as observed in modern aquatic CCMs where the occurrence of a cellular CA, outside a carboxysome for example, dissipates an accumulated HCO_3_^-^pool as membrane-permeable CO_2_ and leads to a high-CO_2_-requiring phenotype (42).

**Fig. 7.**
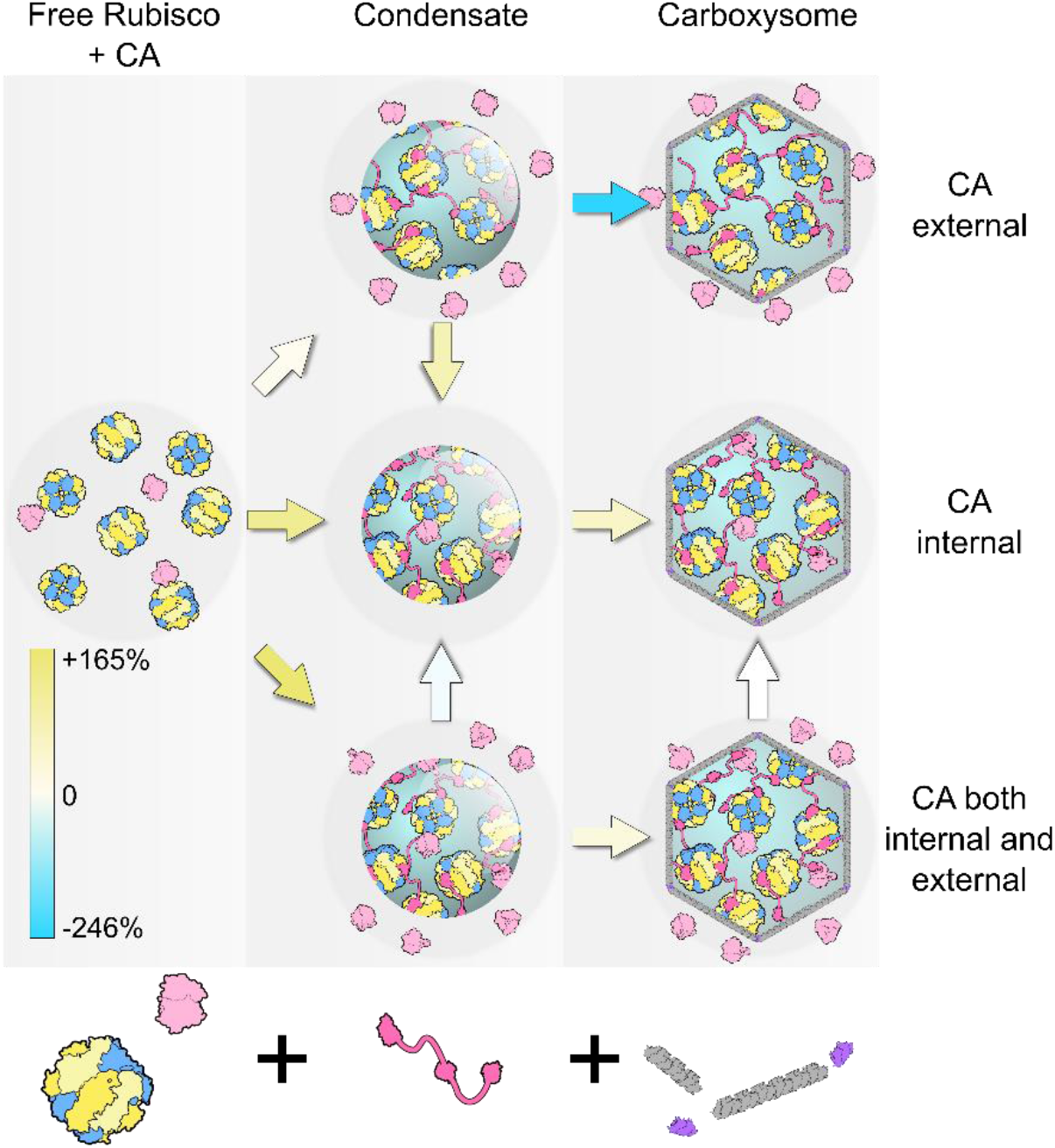
Proposed evolution pathways to carboxysomes from free Rubisco via condensation, before the advent of C_i_ transport. Model simulations propose condensation of Rubisco (*blue/yellow*) in the presence of a cellular carbonic anhydrase (CA) enzyme (*light pink*), here presented as three possible evolutionary alternatives with the CA external (in the unstirred layer), internal, or both external and internal of the condensate. More detailed analysis shows that evolution of a Rubisco condensate in the absence of a CA is not feasible (Fig. 6). Condensation is proposed to occur through the evolution of a condensing protein factor (here CcmM from β-carboxysomes, *bright pink*), and carboxysome formation via the acquisition of bacterial microcompartment shell proteins (*grey, purple*). Contemporary carboxysomes are represented by those containing only internal CA. Percent increase or decrease in average net carboxylation rates between each proposed evolutionary intermediate is indicated by the colored arrows, and the color-scale indicates the values presented in Fig. 6. Between proposed evolutionary stages, *yellow-shaded* arrows indicate an improvement in average net carboxylation rate and *blue-shaded* arrows a net decrease, suggesting a loss in competitive fitness. No net change is indicated by a *white arrow*. The same pattern of potential evolutionary improvements is apparent regardless of the Rubisco source or carboxysome size used in the model, assuming sufficient RuBP supply (*SI Appendix, SI datasets* S1 and S2). We assume the adaptation of increased cellular HCO_3_ ^-^ followed as an evolutionary enhancement (14), hence the relative fitness of systems with CA external to the Rubisco compartment (here modelled in the unstirred layer), where in contemporary systems this is problematic (42).

In Fig. 6 we provide an example evolutionary matrix for hypothesized progressions from a free Rubisco enzyme to a contemporary carboxysome. As a primary evolution, we speculated that condensation of Rubisco could have occurred either with or without co-condensation of CA (‘Condensate + int CA’ or ‘Condensate + ext CA’), or an alternate evolution where some CA was co-condensed and some remained external to the condensate in the unstirred layer (‘Condensate + int/ext CA’). All three possibilities gave rise to condensates with improved fitness over the free enzyme (Fig. 6, Fig. 7, and *SI Appendix, SI* datasets S1 and S2). While the greatest improvement was calculated for a condensate with both internal and external CA (‘Condensate + int/ext CA’), this state showed a negative transition in evolving to a condensate with only internalized CA (‘Condensate + int CA’).

Following initial condensate formation, we propose that the next probable advancement would be the acquisition of bacterial micro-compartment proteins (8) to form a shell with enhanced diffusional resistance. Within the model, only condensates with internalized CA enzymes (‘Condensate + int CA’ and ‘Condensate + int/ext CA’) displayed an improved CO_2_ fixation phenotype during a single step acquisition of a carboxysome shell (Fig. 6, Fig. 7 and *SI Appendix, SI* datasets S1 and S2). There was no difference in fitness phenotype between carboxysomes with internal CA (‘Carboxysome + int CA’) and those with CA both internal and external in the unstirred layer (‘Carboxysome +int/ext CA’) in the model. Further improvements could be made by the addition of HCO_3_ ^-^ transporters, the loss of unstirred layer CA, the acquisition of specific internal CAs, and evolution of Rubisco kinetic properties.

### Low light may drive condensate and carboxysome evolution

Observing increased relative responses of compartment CO_2_ to low RuBP concentrations in the model, where condensate pH is maximally decreased (*SI Appendix*, Fig. S4), we assessed the relative fitness of condensates and carboxysomes at sub-saturating RuBP (50 µM) over HCO_3_^-^ranges in the model. Low cellular RuBP can generally be attributed to light-limited RuBP regeneration via the Calvin Cycle in photoautotrophs (50). A concentration of 50 µM RuBP is approximately three-times the *K*_*M*_*RuBP* of the tobacco Rubisco used here (Table 2) and supports ≈ 63% of the CO_2_-saturated rate for a condensed Rubisco (*SI Appendix*, Fig. S6). At 50 µM RuBP we observe enhanced net carboxylation turnover in the condensate compared with the free enzyme, especially at low HCO_3_ ^-^ (*SI Appendix*, Fig. S15). An additional benefit can be observed for very small carboxysomes since changes in the apparent *K*_*M*_*RuBP* are size-dependent (*SI Appendix*, Fig. S6 and Fig. S15). However, 50 µM RuBP is insufficient to support appreciable carboxylation in a modelled large carboxysome due to decreased substrate permeability (*SI Appendix*, Fig. S6 and Fig. S15).

## DISCUSSION

### The functional advantages of a condensate/carboxysome

The modelling of both Rubisco condensates and carboxysomes in this study demonstrates a number of factors which we predict play a key role in the function and evolution of these Rubisco “organelles”. Firstly, the formation of a Rubisco condensate creates a localized environment in which HCO_3_ ^-^ can be converted to CO_2_ in the presence of CA. CO_2_ can be elevated relative to the external environment by the creation of a viscous unstirred protein solution boundary layer. The presence of condensates or carboxysomes in prokaryotic cells or chloroplasts, where protein concentrations are high, would favor this (51, 52).

The condensation of Rubisco results in the co-localization of Rubisco-reaction protons. This enhances the potential to elevate CO_2_ by driving both the conversion of HCO_3_ ^-^ to CO_2_ in the Rubisco compartment, and decreasing compartment pH under certain conditions (*SI Appendix*, Fig. S4). This role for protons is seen most clearly in carboxysomes where both CA and Rubisco activity are highly dependent on proton production by the Rubisco reaction (Fig. 3, and *SI Appendix*, Fig. S13). Rubisco condensates show a smaller enhancement by Rubisco proton production due to greater permeability to protonated RuBP and PGA, but under conditions of low RuBP and low HCO_3_ ^-^, condensate pH can be lowered and CO_2_ elevated (Fig. 3 and *SI Appendix*, Fig. S4). In addition, the advantages of condensate formation are enhanced when the ratio of external HCO_3_ ^-^ to CO_2_ is increased (*SI Appendix*, Fig. S12) as would occur at higher cytoplasmic pH, and in the presence of active HCO_3_ ^-^ accumulation and subsequent transfer of CA from the cytoplasm to the Rubisco compartment.

The modelling emphasizes that the exchange of protons between internal and external environments is probably independent of free proton or H_3_O^+^ diffusion. Instead, the exchange is dominated by the movement of protonated RuBP and PGA (with *pKa*’s around 6.6), and efflux of CO_2_ which consumes a proton internally (Fig. 4 and Fig. 5; *SI Appendix*, Fig. S5).

### The effects of Rubisco compartment pH and the relevance of sugar phosphate proton carriers

By accounting for the *pK*_*a*_ of physiologically relevant phosphate groups on RuBP and PGA (*SI Appendix*, Fig. S1), the model reveals that these species allow for sufficient ingress of protons into a Rubisco compartment to drive higher rates of carboxylation. This is because the concentration gradients for RuBP and PGA are significantly high (in the mM range), compared with protons at pH 8.0 (in the nM range), enabling them to act as proton carriers in physiologically relevant concentrations. The sophisticated model of Mangan, Flamholz, Hood, Milo and Savage (38) also considers the net production of a proton within the carboxysome [i.e. consumption of one proton by the CA dehydration reaction and the generation of two protons by Rubisco carboxylation]. They calculate that a relatively acidic carboxysome is possible under steady-state conditions, depending on proton permeability. However, analysis of viral capsid shells which have some similarity to the carboxysome icosahedral shell, suggest proton transfer may need to be mediated by specific channels (53, 54). It is important to establish the real permeability of the carboxysome shell to the hydronium ion, and whether high rates of exchange are mediated by channels, proton wires or shuttles linked to protonated sugar phosphates suggested here.

### Drivers of Rubisco compartment evolution

Our evolution analysis suggests that the specificity which Rubisco has for CO_2_ over O_2_ (*S*_*C/O*_) is a likely key driver in determining fitness for the free Rubisco enzyme under low CO_2_ atmospheres (*SI Appendix, SI Results*; Fig. S16). Indeed, the tobacco enzyme, having the greatest *S*_*C/O*_ (Table 2) displayed the best performance under all low CO_2_ scenarios (Fig. 8). Rubisco carboxylation efficiency (i.e. its carboxylation rate constant; *k*_*cat*_^*C*^/K_M_CO_2_) is also an important driver in determining relative fitness under low O_2_ atmospheres, for both the free enzyme and condensates (Fig. S16 – Fig. S19). Furthermore, this analysis suggests that under increased atmospheric O_2_ (as would have occured some 300 million years ago, when levels of O_2_ rose and CO_2_ fell; 14) there is selective pressure to reduce *k*_*cat*_ ^*C*^ of the free enzyme (Fig. S16). This suggest that ancestral cyanobacterial Rubisco may have had kinetics similar to the tobacco Form IB enzyme, implying relatively high photorespiration rates in pre-CCM cyanobacteria. Notably, contemporary cyanobacteria appear to contain a full suite of photorespiratory genes (55), despite limited Rubisco oxygenase activity in modern carboxysomes (56).

**Fig. 8.**
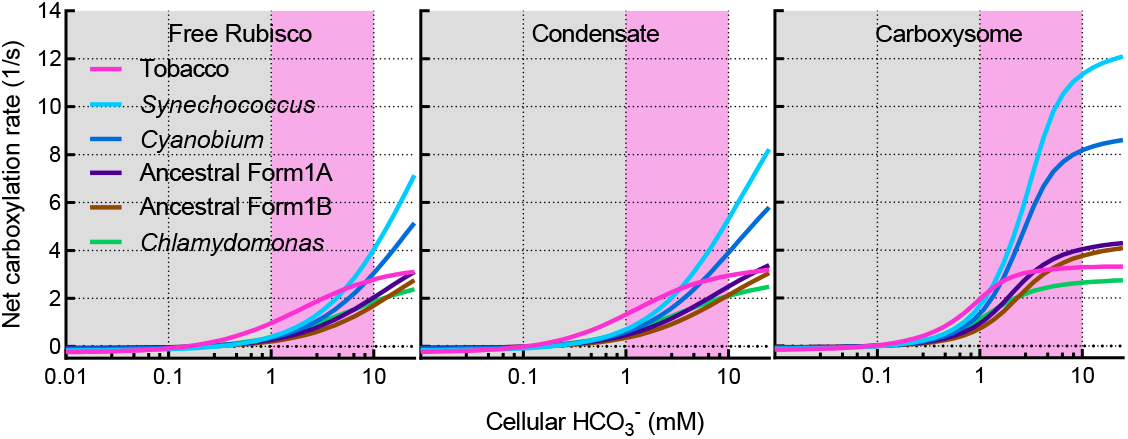
Net Rubisco carboxylation turnover rates in competing enzymes. Net carboxylation turnover rates of competing Rubisco enzymes (Table 2) in the model as free enzymes, condensates or carboxysomes. Each state was modelled using the parameters outlined in Table 1 and as described in the Methods. Rates are depicted under low (0.01 – 1 mM HCO_3_ ^-^; *grey shaded area*) and high (1 – 10 mM HCO_3_ ^-^; *pink shaded area*) CO_2_ environments. Data presented here are calculated under a 20% (*v/v*) O_2_ atmosphere and assume no active accumulation of HCO_3_ ^-^.

Together, our evolution analysis suggests that a low CO_2_ atmosphere may be a key driver in the initial formation of Rubisco condensates. At elevated CO_2_, regardless of O_2_ concentration, condensate fitness is unconstrained by Rubisco catalytic parameters (*SI Appendix*, Fig. S17 – Fig. S19). Large carboxysome formation likely provided compartment conditions which enabled the evolution of Rubisco enzymes with higher *k*_*cat*_^*C*^, but appear to be unconstrained by [O_2_] in the atmosphere (*SI Appendix*, Fig. S20). The correlation between *k*_*cat*_^*C*^ and fitness is apparent only in this scenario, since improving the maximum carboxylation turnover rate is the only means to improve net carboxylation rates when extremely high CO_2_ concentrations around the enzyme can be achieved. Smaller carboxysomes, however, appear not to be driven by any Rubisco catalytic parameter at elevated CO_2_ (*SI Appendix*, Fig. S21). Changes in atmospheric O_2_ may have led to the selection of enzymes with better specificity, catalytic efficiency, and *K*_*M*_*RuBP* during intermediate stages of Rubisco condensate evolution (*SI Appendix*, Fig. S17 – Fig. S19).

The relative response of Rubisco condensates and small carboxysomes to low RuBP supply suggests that low light may also provide conditions conducive to condensate and carboxysome evolution. Low light generally leads to low RuBP (50). In the model such conditions lead to greater advantage of Rubisco condensates over large carboxysomes (*SI Appendix*, Fig. S15). This highlights that elaborations on simple Rubisco condensation can be afforded through facilitated substrate supply or potential diffusion barriers other than carboxysome formation. The association between pyrenoids and thylakoids, for example, suggests that luminal protons can possibly contribute towards the conversion of HCO_3_ ^-^ to CO_2_ (56), and allows the possibility that the pyrenoid function may be enhanced by the combined action of both Rubisco and thylakoid generated protons. This result highlights alternative evolution of Rubisco compartments such as pyrenoids which would provide enhancements exclusive of shell formation.

### Limitations of the model

The model is designed as an idealized Rubisco compartment within a static environment. We use a single ‘condensate’ of 1 µm in radius for the demonstration of condensate function, approximating a large pyrenoid (57). Notably, both carboxysomes and pyrenoids range significantly in size (8, 58), and the effect of compartment size in the model is addressed in the *SI Appendix, SI Methods* and Fig. S22. It does not include an ‘extracellular’ compartment from which the cytoplasmic compartment can receive C_i_ via either diffusion or specific C_i_ pumps. We do not model the system as a functional CCM, holding a static equilibrium between C_i_ species in the system, rather than a disequilibrium which occurs in cyanobacterial cells (59). In the absence of active HCO_3_^-^pumps or a cytoplasmic CA, preferential diffusion of CO_2_ through the cell membrane, coupled with its draw-down via carboxylation by Rubisco, would lead to relatively low cytoplasmic [C_i_] with a HCO_3_^-^:CO_2_ ratio possibly favoring CO_2_ (*SI Appendix*, Fig. S12). However, notably there is no fitness benefit to Rubisco condensation in the absence of a CA (Fig. 6) and scenarios where the HCO_3_^-^:CO_2_ ratio lean heavily toward CO_2_ are less likely.

We do not apply a pH sensitivity to Rubisco catalysis in the model and omit unknown contributors to cellular buffering. This is both for simplicity and to highlight that without pH modulation in the model we would see dramatic changes in pH. Previous models (38) apply the pH-dependency of Rubisco as established by Badger (47), as would be applicable in this instance. We would expect to see a decrease in Rubisco activity in our model resulting from acidification or alkalization of the Rubisco compartment. However, we do not currently know how a condensate or carboxysome might modulate pH changes in reality, and how these might affect the enzymes within it. We also do not model CO_2_ and Mg^2+^ dependencies upon Rubisco activation (47), which would be relevant considerations, especially at low CO_2_ supply, from a physiological standpoint. These considerations may underpin mechanistic controls at low substrate supply within both compartment types.

Importantly, we note that the permeabilities for diffusion across the compartment interfaces have no physical measurements to support them. The values for carboxysomes are in line with previous models but it should be realized that these values have been derived from model analysis and not independent physical measurements. The values assumed for condensates, although seeming reasonable and supporting a functional model, have not been physically established independently.

### Conclusions

Accounting for proton production by Rubisco reactions, their utilization in the CA reaction, their transport via RuBP and PGA, and applying diffusional resistance to their movement, our model highlights a role for protons in Rubisco condensate and carboxysome function and evolution. Application of our model to proposed evolutionary intermediate states prior to contemporary carboxysomes provides a hypothetical series of advancements, and suggest that low CO_2_ and low light environments may be key environmental drivers in the evolutionary formation of Rubisco condensates, while increases in atmospheric O_2_ may have played a role in Rubisco catalytic parameter selection. Our modelled outcomes are achieved through assumption of diffusional resistances to reaction species, which align well with previous models but remain to be determined experimentally. Taken together, our analysis provides insights into the function of phase-separated condensates of proton-driven enzyme reactions.

## METHODS

### Mathematical modelling

Modelling of Rubisco compartment scenarios and data output were carried out using the biochemical network simulation program COPASI (copasi.org), described by Hoops, *et al*. (60). COPASI (v4.25, build 207) was used to simulate reaction time-courses achieving steady-state conditions in a three-compartment model where reaction species are linked in a biochemical network (Fig. 1). For standard modelling conditions catalytic parameters of the tobacco Rubisco were used, while those of *C. reinhardtii, C. marinum* PCC7001 and *S. elongatus* PCC7942, and predicted ancestral Form IA and Form IB enzymes (15) were also used in evolutionary fitness analysis (Table 2). Michaelis-Menten rate equations were applied to Rubisco catalysis as dependent upon substrate and inhibitor concentrations within the Rubisco compartment (*SI Appendix, SI Methods*). O_2_ was applied as an inhibitor of the carboxylation reaction and CO_2_ an inhibitor of oxygenation. Greater detail of model parameterization is provided in the *SI Appendix, SI Methods*. The ordinary differential equations describing the model components can be found in the *SI Appendix*. Reactions, reaction species and model parameters can be found in *SI Appendix*, Table S1, Table S2, and Table S3 respectively.

Variation in compartment types (i.e. the free enzyme, a condensate, or a carboxysome) was simulated in the model by varying unstirred boundary and condensate permeabilities to all reaction species (Table 1, *SI Appendix, SI Methods*). The permeabilities to RuBP^3-^ and RuBP^4-^ were assumed to be the same, as were those for PGA^2-^ and PGA^3-^, and a single permeability value applied to either RuBP or PGA species. Proton production by carboxylation and oxygenation reactions was varied by adjusting the proton stoichiometry for either reaction (*SI Appendix, SI Equations*, Table S1). Protonation and deprotonation of RuBP and PGA in each compartment was enabled by assigning rate constants equivalent to their *pKa* values (*SI Appendix*, Table S3).

The size of the external compartment was set to 1 m^3^ within the model, and both the Rubisco compartment and unstirred boundary layer volumes determined by setting the Rubisco compartment radius. For standard modelling procedures we used a spherical Rubisco compartment radius of 1 × 10^-6^ m^3^ and the unstirred boundary layer volume was determined as a simple multiplier of the Rubisco compartment radius. The Rubisco compartment radius used in our modelling generates a large condensate or carboxysome, akin to contemporary pyrenoids, however variation in the compartment size has little effect on the conclusions of the modelled outcomes since even small carboxysomes display higher Rubisco turnover rates that large condensates in the model (*SI Appendix*, Fig. S22). Extremely small carboxysomes have lower amplitude responses to proton and RuBP permeabilities (Fig. 2) and size may have led to favorable Rubisco kinetics during evolution to modern carboxysomes (*SI Appendix*, Fig. S21, *SI dataset* S2).

CO_2_ concentration in the external compartment was set to 0.01 × external [HCO_3_ ^-^] (*SI Appendix, SI Equations*), assuming negligible effects of a single Rubisco compartment on the bulk external C_i_ species. Interconversion between CO_2_ and HCO_3_^-^was allowed to proceed in the unstirred and Rubisco compartments with rate constants of 0.05 for the forward reaction (CO_2_ → HCO_3_^-^+ H^+^), and 100 for the back reaction (HCO_3_ ^-^+ H^+^ → CO_2_). CA contribution was enabled by applying a multiplying factor to each rate, such that a factor of 1 simulates the absence of CA. Typical CA multiplying factors for each type of modelled compartment are listed in Table 1. CA function external to a Rubisco compartment was modelled by modifying CA in the unstirred layer (Table 1).

O_2_ concentration in the model was typically set at contemporary atmospheric levels 20% (*v/v*) by assigning a concentration of 0.25 mM, the water-soluble concentration at 25 °C. For simulations at 30% (*v/v*) O_2_ (an estimated volumetric concentration in the atmosphere when it is proposed CCMs arose; 14), a concentration of 0.36 mM was used.

Steady-state reactions were initialized by setting the reactant concentrations in the external compartment and running the model to achieve steady-state. External pH was set at 8.0 using a compartment [H^+^] of 1 × 10^-5^ mM. For saturating Rubisco substrate concentrations, initial [HCO_3_ ^-^] was set to 20 mM and [RuBP^4-^] was set to 5 mM. Interconversion of RuBP^4-^ and RuBP^3-^ at *pKa* 6.7 and pH 8.0 results in ≈ 95% of all RuBP as the RuBP^4-^ species.

Rubisco site concentrations were typically set to 10 mM. This value is similar to that calculated for α-and β-carboxysomes (61) although higher than that estimated for pyrenoids (18). It is nonetheless a reasonable upper limit for the purposes of examining system responses within the model.

### Evolution analysis

Hypothesized free enzyme, condensate or carboxysome evolutionary intermediates (Fig. 6) were generated and analyzed within the model using the parameters in Table 1, applying the catalytic parameters of each Rubisco enzyme in Table 2. For each scenario the model was run over a range of [HCO_3_^-^] from 0.01 to 25 mM, at saturating RuBP (5 mM). Both carboxylation and oxygenation rates were output for each scenario and converted to turnover numbers by accounting for active site concentrations within the model. Net carboxylation turnover rates (1/s) were calculated by assuming a photosynthetic cost of ½ mole of CO_2_ loss for each mole O_2_ fixed (*SI Appendix*, Fig. S14). Modelling was carried out at both 20% and 30% O_2_ (*v/v*; Table 1). Performance comparisons of each hypothesized evolutionary state were made for both low CO_2_ (0.01 – 1 mM HCO_3_ ^-^) and high CO_2_ (1 – 10 mM HCO_3_ ^-^) ranges by calculating the average net carboxylation rates for each scenario at each CO_2_ range and O_2_ concentration (*SI Appendix*, Fig. S14). Fitness comparisons were determined from the absolute differences in average net carboxylation rates between each modelled scenario (Fig. 6; *SI Appendix, SI datasets* S1 and S2). An additional dataset was calculated for the tobacco Rubisco enzyme at 20% (*v/v*) O_2_ and 50 µM RuBP to determine compartment-type performance under conditions simulating low light.

Correlations between average net carboxylation rates (over specified modelled atmospheres) and Rubisco catalytic parameters (Table 2) for each proposed evolutionary state (Fig. 6; *SI Appendix*, Fig. S16 – Fig. S21) were calculated as Pearson correlation statistics, with *p* values calculated as two-tailed distribution of calculated *t*-statistics (*SI Appendix, SI Results, SI datasets* S1 and S2). In our results we highlight correlations where *p* < 0.05. For plotting purposes (*SI Appendix, SI Results*, Fig. S16 – Fig. S21) Rubisco catalytic parameters (Table 2) were normalized to the largest value in each parameter set and plotted against average net carboxylation rates.

## AUTHOR CONTRIBUTIONS

M.R.B designed and developed the model. B.M.L. and B.F. carried out modelling experiments. B.M.L. and M.R.B generated the Figures. B.M.L. wrote the manuscript draft. B.M.L., B.F., S.B.P., G.D.P. and M.R.B. discussed, wrote and edited manuscript revisions.

## ACKNOWLEDGEMENTS

This work is supported by a sub-award from the University of Illinois as part of the research project Realizing Increased Photosynthetic Efficiency (RIPE) that is funded by the Bill & Melinda Gates Foundation, Foundation for Food and Agriculture Research, and the U.K. Government’s Department for International Development under grant number OPP1172157, and an award from The Australian Research Council, Centre of Excellence grant for Translational Photosynthesis (CE140100015) to M.R.B., G.D.P. We thank Angus Long for enabling rapid processing of COPASI data output, and Avi Flamholz, Luke Oltrogge and Matthew Mortimer for comments on the manuscript.

## Notes

http://dx.doi.org/10.17632/c52km273vv.3

### Competing Interest Statement

The authors have declared no competing interest.

### Summary of Updates

Changes to Figure 1 and Table 1 to highlight modelling conditions for the external compartment. A typographical error in Table 2 has been corrected, changing the value 3.30 to 33.0

