## Supplementary Information for "Rubisco proton production can drive the elevation of CO_2_ within condensates and carboxysomes"

**This PDF file includes:**

SI Results
SI Methods
Tables S1 to S3
Figures S1 to S22
SI Equations
Legends for Datasets S1 and S2
SI references

**Other supplementary materials for this manuscript include the following:**

Datasets S1 and S2

**Data availability:**

All supplementary datasets and the COPASI model files (Free Rubisco model.cps, Condensate model.cps, Small carboxysome model.cps, and Large carboxysome model.cps) are available at the following link: http://dx.doi.org/10.17632/c52km273vv.3. The COPASI file can be accessed using the freely available COPASI software available at copasi.org

### SI Results

#### Which Rubisco catalytic parameters correlate with evolutionary fitness?

Applying the same evolutionary fitness analysis to the extant and extinct Rubiscos in our analysis (Table 2), we were able to estimate the Rubisco catalytic properties that would make the most competitive advances through condensation and encapsulation prior to evolution of mechanisms enabling $HCO_3^-$ accumulation. We determined correlation statistics for the average net carboxylation turnover rates (1/s) under different atmospheres, and each catalytic parameter (Table 2), for the most likely evolutionary steps determined from the initial analysis (Fig. 7). These analyses were modelled over a range of $HCO_3^-$ in the model at an [RuBP] of 5 mM, except for 'low light' simulations where we set RuBP to 50 µM. We draw the reader's attention to *SI Appendix, SI datasets* S1 and S2 and corresponding correlation plots in Fig. S16 to Fig. S21, where we present the data for the analysis presented in this section.

Regardless of the proposed step in the evolution toward contemporary carboxysomes (Fig. 7), the tobacco Rubisco, characterized by a high specificity ($S_{C/O}$) for $CO_2$ over $O_2$, and high carboxylation efficiency ($k_{cat}^C/K_MCO_2$, 1/s/µM; Table 2), had the greatest average net carboxylation turnover rate at the low $CO_2$ range (0.01 – 1 mM cytoplasmic $HCO_3^-$; Fig. 8). This was observed under both high (30% *v/v*) and low (20% *v/v*) $O_2$ atmospheres.

Competitive fitness analyses at the high $CO_2$ range (1 – 10 mM cytoplasmic $HCO_3^-$; Fig. 8) indicated that the contemporary carboxysomal Rubisco from *Synechococcus* (characterized by low $S_{C/O}$, high $k_{cat}^C$, high $K_MCO_2$, and high $K_MRuBP$, Table 2) performed best in all proposed evolutionary stages, regardless of atmospheric $O_2$, with the exception of free enzyme comparisons where it was out-competed by the tobacco enzyme.

For the free enzyme ('Rubisco + CA') at both low $CO_2$ and low $O_2$, $S_{C/O}$ and $k_{cat}^C/K_MCO_2$ displayed strong positive correlations with the average net carboxylation rate ($r = 0.86 – 0.99$, $p < 0.05$). However, under low $CO_2$ and elevated $O_2$, while $S_{C/O}$ still correlated positively with average net carboxylation rate ($r = 0.83$, $p = 0.042$), carboxylation turnover ($k_{cat}^C$) showed a strong negative correlation ($r = -0.91$, $p = 0.011$), indicating a selective pressure for high specificity and low catalytic turnover for the free enzyme under elevated $O_2$. At elevated $CO_2$, there was no Rubisco parameter which correlated with average net Rubisco carboxylation rate, regardless of $O_2$ concentration.

For Rubisco condensates (with either internal, external, or both internal and external CA) correlations between average net carboxylation rate and Rubisco catalytic parameters were very similar to the free enzyme. Again, under low $CO_2$ and low $O_2$, $S_{C/O}$ and $k_{cat}^C/K_MCO_2$ correlated positively with average net carboxylation turnover ($r = 0.85 – 0.99$, $p < 0.05$), the only exception being an apparent negative correlation between net carboxylation and $K_MRuBP$ under these conditions for a condensate with an external CA ($r$ = -0.83, $p$ = 0.041). This implies that a Rubisco condensate with an external CA would be under selective pressure to improve Rubisco RuBP utilization under low $CO_2$ and $O_2$.

At elevated $O_2$ and low $CO_2$, a condensate with an external CA (modelled by modification of the unstirred layer CA function) appeared to show similar dependencies on reduced $k_{cat}^C$ and improved $S_{C/O}$ as the free enzyme. However, for condensates with internal CAs ('Condensate + int CA' and 'Condensate + int/ext CA'), net carboxylation correlated with increased $S_{C/O}$ and $k_{cat}^C/K_MCO_2$ ($r$ = 0.83 – 0.99, $p < 0.05$).

Like the free enzyme, there was no correlation between net carboxylation rates and any Rubisco catalytic parameter for any condensate type at high $CO_2$, regardless of the $O_2$ concentration. This implies no selective pressure to evolve the Rubisco catalytic parameters analyzed under a high $CO_2$ environment for either free Rubisco or Rubisco condensates.

Comparative analysis of different Rubisco enzymes in large carboxysomes showed there was a strong correlation between $k_{cat}^C/K_MCO_2$ and net carboxylation rates at low $CO_2$ ($r$ = 0.89, $p < 0.05$), regardless of $O_2$ concentration. At high $CO_2$ concentrations (the likely cellular environment for contemporary carboxysomes in functional CCMs), $k_{cat}^C$ was the only Rubisco catalytic parameter to correlate with net carboxylation rate in the model ($r$ = 0.98, $p < 0.05$). For small carboxysomes, where a higher surface area/volume ratio means greater diffusional flux of substrates and products (Fig. S22), both $k_{cat}^C/K_MCO_2$ and $S_{C/O}$ correlated with net carboxylation rates at low $CO_2$ ($r$ = 0.88 – 0.98, $p < 0.05$), suggesting that carboxysome size may correlate with catalytic efficiency and specificity, possibly due to lower $CO_2$ availability in smaller carboxysomes (Fig. S22). Unlike their larger counterparts, small carboxysomes appear unconstrained by any catalytic parameter when $CO_2$ is high.

### SI Methods

### Model description

*Rubisco function in the model*

For the purpose of modelling the function of Rubisco enzymes in a condensate, we examine a compartment filled with Rubisco active sites as an isolated entity within a larger compartment supplied with the theoretical substrates and products of the enzyme (Fig. 1). Rubisco utilizes ribulose-1,5-bisphosphate (RuBP) as its 5-carbon substrate which can either be carboxylated with $CO_2$ to produce two molecules of the 3-carbon product 3-phosphoglycerate (PGA), or oxygenated with $O_2$ to produce one molecule of PGA and one molecule of 2-phosphoglycolate (2PG). Notably, Rubisco carboxylation is competitively inhibited by $O_2$ while the oxygenation reaction is competitively inhibited by $CO_2$ (1).

In this study we include the net production of two protons with each reaction, as outlined below and in greater detail in Fig. S1.

$$CO_2 + RuBP \xrightarrow{carboxylation} 2 \times PGA + 2 \times H^+$$

$$O_2 + RuBP \xrightarrow{oxygenation} 1 \times PGA + 1 \times 2PG + 2 \times H^+$$

Rubisco carboxylase and oxygenase reaction velocity (*V*) are both driven in the model by irreversible Michaelis-Menten kinetics, coupled with terms for inhibition for both $O_2$ and $CO_2$ as follows:

$$V = \frac{\left(\frac{[RuBP]}{[RuBP] + Km_{RuBP}}\right) \times V_{max} \times [S]}{Km + [S] + \frac{Km \times [I]}{Ki}}$$

where [*S*] is either the $CO_2$ or $O_2$ concentration in the Rubisco compartment and *Km* is the Michaelis-Menten constant for the relevant gas (Table 2). *I* and *Ki* refer to the concentration and Michaelis-Menten constant of the opposing gaseous substrate in the Rubisco compartment, respectively, allowing for inhibition of each reaction by the competing substrate. Maximum rates of either carboxylation or oxygenation ($V_{max}$) are dependent on the enzyme parameters used in modelling experiments (Table 2). The concentration of substrate RuBP within the Rubisco reactions is specifically the deprotonated species, $RuBP^{4-}$ (which exists as the predominant form [≈95%] at pH 8.0) and the *Km* for RuBP is dependent upon the source of each Rubisco enzyme used in modelling experiments (Table 2). Since 2PG is both lost from the Calvin-Benson Cycle (2), and typically in negligible quantities under most conditions due to the specificity of Rubisco for $CO_2$ over $O_2$ (3), we do not include 2PG as a component in the model.

*Reaction species and protons in the model*

In contrast with existing models which address carboxysome function within complex cellular scenarios within CCMs with active $HCO_3^-$ accumulation (4-7), here we assume no $HCO_3^-$ accumulation and modify reaction output by enabling two moles of $H^+$ to be produced as by-products of both the carboxylase and oxygenase reactions along with PGA (above). Proton balance within the system is maintained by setting the external pH to a constant value, while protons are allowed to interact with other chemical species by reversible reactions in each model compartment (Fig. 1). Specifically, we allow $H^+$ take part in the protonation and deprotonation of RuBP and PGA species within the model, driven by their *pKa* values (6.7 and 6.5 respectively), yielding variable quantities of $RuBP^{4-}$:$RuBP^{3-}$ and

$PGA^{3-}$:$PGA^{2-}$ depending on the pH of each compartment, and driven by their respective rate constants as follows:

$$RuBP^{3-}\ or\ PGA^{2-} \leftrightarrow RuBP^{4-}\ or\ PGA^{3-} + H^{+}$$

The second order association constants used for $RuBP^{4-}$ and $PGA^{3-}$ were $5.01187 \times 10^9$ and $3.15 \times 10^9$ $m^3/(mol.s)$ respectively (corresponding to the *pKa* of each substrate), and the first order dissociation constants were $1 \times 10^6$ 1/s for both $RuBP^{3-}$ and $PGA^{2-}$.

We also include the interconversion of $HCO_3^-$ and $CO_2$ as a reaction which utilizes a proton (Fig. 1), and we enable control of the rate of this interconversion by carbonic anhydrase (CA) using rate constants for the forward ($k_1$; $CO_2$ to $HCO_3^-$) and backward ($k_2$; $HCO_3^-$ to $CO_2$) reactions.

forward reaction $$CO_2 + H_2O \xrightarrow{CAfactor \times k_1} HCO_3^- + H^+$$

backward reaction $$HCO_3^- + H^+ \xrightarrow{CAfactor \times k_2} CO_2 + H_2O$$

The first order rate constants $k_1$ (0.05/s) and $k_2$ (100/s) allow attainment of $HCO_3^-$:$CO_2$ ratio of 100:1, which approximates the proportion, assuming the uncatalyzed interconversion of each species at chemical equilibrium when pH is 8.0. Notably this ratio is typically much higher in cells displaying functional CCMs where a disequilibrium favoring $HCO_3^-$ is maintained (8). Here we deliberately model a primordial condition with no CCM in these experiments (see below).

An additional multiplying '*CA factor*' is used when modelling the presence of CA activity in specific model compartments, where a *CA factor* of 1 represents the absence of CA activity, assuming the attainment of chemical equilibrium at the uncatalyzed rate. Where CA is present in the model we use a value $1 \times 10^5$ for this parameter (Table 1), as used in the model of Reinhold, Kosloff and Kaplan (7).

Compartment pH is calculated from total $H^+$ concentration at any given scenario in the model, driven primarily by setting the 'external' compartment to pH 8.0 with an $H^+$ concentration of 10 nM. This value approximates the pH of a typical cyanobacterial cellular environment which has been shown to vary between pH 7.3 in the dark and pH 8.3 in the light (5). This range represents a ten-fold variation in total proton concentration in the external compartment (from 5 – 50 nM). The lower pH would seem to favor carboxylation as it would lead to a lower Rubisco compartment pH and greater $CO_2$ production. However, it is unlikely to lead to significant changes in Rubisco carboxylation rates inside the compartment, as it equates to a small decrease in compartment proton permeability where internal proton concentrations are maintained and maximum carboxylation rates are observed (Fig. 2). The higher pH value of 8.3 represents a halving of the proton concentration from 10 to 5 nM compared with our standard modelling scenarios, however proton flux will still be predominantly maintained by other reaction species (Fig. 5), and is therefore unlikely to make a substantial difference to model outputs.

We nonetheless make comparisons between scenarios with consistent pH for the purposes of demonstrating the role of protons in the system.

*RuBP concentrations used in modelling*

RuBP is typically found within cyanobacterial and microalgal cells within the range of 0 – 50 mM, depending on external $C_i$ supply and light-limited photosynthetic rate (9, 10). Regardless of the catalytic requirements, most Rubisco enzymes experience RuBP concentrations well above their $K_M RuBP$ (which is typically in the µM range; Table 2), both in the modelling presented here and under most growth conditions, except for low light (11). For most modelling scenarios we adopt 5 mM RuBP as a saturating supply concentration.

The application of diminished permeability to RuBP access in both condensate and carboxysome modelling, we find that this results in an increase in the apparent $K_M RuBP$ (Fig. S6), an observation made for comparison of free Rubisco and carboxysomes isolated from *Cyanobium* (12). Since the apparent $K_M RuBP$ differs between each compartment type and size we model, this has the potential to confound compartment comparisons at a specific [RuBP]. To account for this, we arbitrarily chose the RuBP response of a modelled condensate at saturating $HCO_3^-$ (20 mM) and determined the relative carboxylation rate at 50 µM RuBP. This provided ≈63% of the maximum carboxylation rate (Fig. S6). We then determined the concentration of RuBP which provided an equivalent relative rate for the free enzyme (35 µM), and both small and large carboxysomes (87 µM and 1.3 mM, respectively).

*$CO_2$ and $HCO_3^-$ concentrations in the 'external' compartment and carbonic anhydrase (CA) activity*

In a cellular context, the role of the relatively membrane-impermeable and dominant inorganic carbon ($C_i$) species $HCO_3^-$ is of importance as a companion species to the Rubisco substrate (and more membrane-permeable) $CO_2$. We therefore include both $HCO_3^-$ and $CO_2$ as the $C_i$ species within the model, and we set $CO_2$:$HCO_3^-$ in this compartment to approximate chemical equilibrium at pH 8.0. (13) While other $C_i$ species ($H_2CO_3$ and $CO_3^{2-}$) are important to consider in cellular modelling contexts (5), for simplicity here we consider $HCO_3^-$ as the major $C_i$ source at ≈pH 8.0, and $CO_2$ as the Rubisco substrate.

In considering the relative concentrations of $HCO_3^-$ and $CO_2$ within the model, we simulate an environment which is not representative of a modern $CO_2$ concentrating mechanism (14) where a disequilibrium toward $HCO_3^-$ is favored (13). Instead we assume the external compartment as a bulk volume at pH 8.0 and with a constant $HCO_3^-$:$CO_2$ ratio, unlikely to be affected by the fluxes of a small

Rubisco compartment or fluctuations resulting from CA activity, and $O_2$ in equilibrium with the prevailing atmospheric gas composition. Current hypotheses predict that active $C_i$ uptake and intracellular $HCO_3^-$ accumulation likely evolved subsequent to the evolution of carboxysomes in cyanobacteria, for example (15). Evolutions leading to an elevation of intracellular $HCO_3^-$ would lead to the need to eliminate cytosolic CA (16) to maintain this pool of $HCO_3^-$. By simulating a primordial system without $C_i$ uptake, or overaccumulation of $HCO_3^-$, we are able to address both; 1) Rubisco compartment behavior alone as a function of $C_i$ supply, and 2) Rubisco compartment responses in predicted evolutionary contexts where we can manipulate the location and speed of CA enzymes, and compartment permeabilities to substrates.

We maintain the external compartment $HCO_3^-$:$CO_2$ ratio at 100:1 in most modelling scenarios, assuming negligible changes to this bulk compartment in response to relatively small fluxes from a single Rubisco condensate. Thus, examination of the effects of external CA activity on Rubisco compartment function are addressed by modifying CA activity in the unstirred layer (Table 1).

*Model reaction compartments and their permeabilities*

The model utilizes three separate reaction compartments termed 'external', 'unstirred' and 'condensate' (Fig. 1). These are analogous, respectively, to a fixed cell cytoplasm, an unstirred boundary layer around the condensate, and the condensate (or Rubisco compartment) itself . We envision the condensate as either a liquid-liquid phase separated protein droplet such a pyrenoid (17), or a carboxysome which is formed via similar structural rules (18) but encased in a protein layer with increased diffusional resistance to substrate passage. Rubisco carboxylation and oxygenation are confined to the 'condensate' compartment while the reversible reactions involving the protonation of RuBP, PGA and $HCO_3^-$ (above) are allowed to occur in all three compartments (Fig. 1).

As highlighted above, we assume that the external compartment, akin to a fixed cell cytoplasm, is not dependent on the function of $C_i$ transporters to elevate [$HCO_3^-$]. Instead we view the external compartment simply as a means to supply substrate to the Rubisco compartment and that it simulates a primordial cellular state prior to CCM evolution. The external compartment volume is set at 1 $m^3$ to enable sufficient substrate supply. Within the context of the analysis carried out here, this volume has little relevance but can be modified to consider alternative substrate supply scenarios.

In the absence of accurately known permeabilities of each substrate through unstirred cellular boundaries and either phase-separated protein matrices or complex protein shell layers, we apply simple assumptions in order to demonstrate the concepts presented. While molecular dynamic modelling suggests that carboxysome shell channel proteins may be selective permeable to substrates (19, 20), actual permeabilities still remain elusive. We therefore assume equal permeabilities for all substrates except for protons (see below) at each diffusional boundary in the model, recognizing that differences may exist in reality, but remain unknown.

The permeability coefficient we have chosen for substrates (other than protons) crossing the unstirred boundary layer is consistent with those calculated for diffusion of small solutes associated with spherical bodies of various sizes (21). For example, a sphere of $\approx 10^{-6}$ m radius (the size we assume for a Rubisco condensate here) is calculated to have an unstirred layer permeability around $10^{-4}$ m/s (21). We also assume that a value of $10^{-4}$ m/s approximates the upper bound of small molecule permeabilities where the viscosity of the crowded cytosol or stroma of the organism in which the aggregates are formed can lead to a slowing of diffusion of up to four times that in water (22). For the same reason, we also consider the protein aggregated Rubisco condensate as having a similar resistance to small molecule flux and assign permeabilities of $10^{-4}$ m/s for all substrates except for protons (below).

For carboxysomes, we assume the shell to have permeability some 100-fold lower than the unstirred layer and Rubisco condensate. Mangan, Flamholz, Hood, Milo and Savage (5) provide a detailed analysis of likely carboxysome shell permeabilities to relevant reaction species and conclude that significant hindrance to passage across the shell is likely. A relatively wide range of permeabilities are consistent with function in their model and ours (e.g. Fig. 2), without application of selectivity to particular species. We use a permeability of $10^{-6}$ m/s for all substrates, except for protons, at the carboxysome shell, which brings each species within the functional range described previously (5), acknowledging an absence of real data to support the specific permeability of each species. Nonetheless, the prime outcome here is that, within carboxysome modelled scenarios, the shell resistance dominates the calculation of diffusion processes into and out of the carboxysome.

We treat the permeability of the unstirred layer and condensate/carboxysome to protons differently in the model. Proton movement is rapid in aqueous environments due to their ability to jump between hydrogen bonds via a water-based ‘proton wires’ mechanism (23). Based on this we have assumed a 100-fold greater permeability for protons than other solutes. We therefore apply a proton permeability of $10^{-2}$ m/s at the unstirred layer boundary and the condensate, and $10^{-4}$ m/s at the carboxysome. This implies some barrier to proton diffusion, however, and is perhaps counter to evidence that proton movement across the shell rapid and unimpeded (24). We highlight within the manuscript that there are a number of issues with this conclusion, and can model the same outcomes of the experiment by Menon, Heinhorst, Shively and Cannon (24) to show that they are not inconsistent with relatively low proton permeability (Fig. S3). In viral capsid systems, which are structurally analogous to carboxysomes, the compartment shell layer forms a significant barrier to proton diffusion (25, 26), and it is assumed that specific proton pores may be needed to admit protons at significant rates to the inside of the virus. It is both feasible and consistent with existing evidence that complex and specific restriction to proton passage across the shell exists.

We apply a simple multiplication function for the thickness of the unstirred layer based on the radius of the Rubisco compartment (Table 1). This generates an unstirred layer of thickness $5 \times 10^{-7}$ m for condensates of $1 \times 10^{-6}$ m radius, and is consistent with vesicles of a similar size range (27). However, the thickness of the unstirred layer is of little consequence in the model. Rather, it is the permeability coefficient which determines the rate limiting nature of the unstirred layer compartment. The compartment acts only as a volume which can come to steady state equilibrium with the external and condensate volumes either side.

*Rubisco compartment size*

When considering appropriate sizes for the modelling of Rubisco compartments, we initially considered a pyrenoid as a canonical Rubisco condensate. These condensates are found in a wide variety of microalgae and hornwort species (28-30), and range widely in size and structure. In some hornworts, for example, pyrenoids can be highly dissected to form many disc-shaped ‘subunits’ as small as ≈25 × 200 nm (31), while the more regular spheroid pyrenoid of *Chlamydomonas reinhardtii* has been observed at 2 µm in diameter (32). For most scenarios, Rubisco condensates were assumed as spheres with radius $1 \times 10^{-6}$ m, approximating that of a pyrenoid from *C. reinhardtii*.

We recognize that carboxysome sizes also vary considerably across genera, and are often quite large in β-cyanobacteria (up to $\approx 6 \times 10^{-7}$ m in diameter; 33, 34) and as small as $\approx 9 \times 10^{-8}$ m wide in some α-cyanobacteria (35). Indeed, β-carboxysome size appears highly variable within the same species and is likely dependent upon growth conditions and stoichiometry of protein components (33, 34). For the purpose of considering a possible evolutionary adaptation of a condensate into a carboxysome, and for simple comparisons between compartment types in the model, we consider both condensates and carboxysomes as spheres with the same radius ($1 \times 10^{-6}$ m).

The size of a condensate or carboxysome will affect both its surface area and internal volume, therefore the diffusional flux, both across its boundary and within it. Compartment size will therefore have a supply-rate effect on Rubisco substrates, and therefore affect Rubisco function in the model. Smaller carboxysomes in the model have less dramatic responses than larger structures with respect to elevation of internal [$CO_2$], due to an increase in internal pH as carboxysome size decreases (Fig. S22). These responses are also driven by substrate concentrations in the external medium, however, with greater decreases in compartment pH (and therefore increases in $CO_2$) apparent at limited substrate concentrations. (Fig. S4, Fig. S22).

Large carboxysomes are capable of greater internal [$CO_2$], leading to carboxylation turnover rates which are approximately 4% higher than the smallest of carboxysomes, and a concomitant decrease in internal [RuBP] (Fig. S22). A similar result is apparent for condensates, although the ability to generate very high internal [$CO_2$] is diminished and the variation in carboxylation turnover is only 2% across the entire size range considered. This also means that the analysis of evolutionary progressions from condensates to carboxysomes may be marginally affected by the size of a carboxysome compared with its progenitor condensate, although notably the carboxylation turnover of even the smallest carboxysome is faster than that of the largest condensate (Fig. S22).

*Rubisco active site concentration*

Rubisco active site concentrations in pyrenoids have been measured at ≈4.8 mM (17), while estimates for carboxysomes are as high as ≈15 mM for tightly packed spherical Rubisco $L_8S_8$ holoenzymes of 11 nm diameter in a crystalline array in both $C_3$ chloroplast stromae (36) and carboxysomes (9), and values around 10 mM calculated from stoichiometries and estimated packing in α-carboxysomes (9, 37). In the absence of more definite data which presents a consolidated value for both pyrenoids and carboxysomes, we therefore set a value of 10 mM active sites as an upper bound of likely concentrations in our simulated Rubisco condensates and carboxysomes, which would allow for movement of holoenzymes within the compartment, and both small molecule and activation chaperone passage. We maintain this value between compartments types in the model for simple comparisons.

Given the relative increase in Rubisco carboxylation turnover in carboxysome scenarios in our model, we would expect carboxysomes specifically to show an upper limit in Rubisco active site concentrations since excessive enzyme amounts will lead to decrease in net carboxylation due to consumption of either RuBP or $CO_2$, depending on which is limiting. This too may be dependent on carboxysome size as this will affect diffusional supply rates (above).

### SI Tables

#### Table S1. COPASI model reactions and their descriptions

| Name | Reaction | Description |
|---|---|---|
| RubiscoC1 | Cc + R4c → [2 × P3c] + [2 × Hc]; Oc | Rubisco carboxylation; inhibition by $O_2$ |
| RubiscoO | Oc + R4c → P3c + [2 × Hc]; Cc | Rubisco oxygenation; inhibition by $CO_2$ |
| CAe forward | Ce → Be + He | Carbonic anhydrase forward reaction in the external compartment |
| CAe back | Be + He → Ce | Carbonic anhydrase backward reaction in the external compartment |
| CAu forward | Cu → Bu + Hu | Carbonic anhydrase forward reaction in the unstirred compartment |
| CAu Back | Bu + Hu → Cu | Carbonic anhydrase backward reaction in the unstirred compartment |
| CAc forward | Cc → Bc + Hc | Carbonic anhydrase forward reaction in the Rubisco compartment |
| CAc back | Bc + Hc → Cc | Carbonic anhydrase backward reaction in the Rubisco compartment |
| DiffCeu | Ce → Cu | Diffusion of $CO_2$ from the external compartment to the unstirred compartment |
| DiffCuc | Cu → Cc | Diffusion of $CO_2$ from the unstirred compartment to the Rubisco compartment |
| DiffBeu | Be → Bu | Diffusion of $HCO_3^-$ from the external compartment to the unstirred compartment |
| DiffBuc | Bu → Bc | Diffusion of $HCO_3^-$ from the unstirred compartment to the Rubisco compartment |
| DiffOeu | Oe → Ou | Diffusion of $O_2$ from the external compartment to the unstirred compartment |
| DiffOuc | Ou → Oc | Diffusion of $O_2$ from the unstirred compartment to the Rubisco compartment |
| DiffHeu | He → Hu | Diffusion of $H^+$ from the external compartment to the unstirred compartment |
| DiffHuc | Hu → Hc | Diffusion of $H^+$ from the unstirred compartment to the Rubisco compartment |
| DiffR4eu | R4e → R4u | Diffusion of $RuBP^{4-}$ from the external compartment to the unstirred compartment |
| DiffR4uc | R4u → R4c | Diffusion of $RuBP^{4-}$ from the unstirred compartment to the Rubisco compartment |
| DiffR3eu | R3e → R3u | Diffusion of $RuBP^{3-}$ from the external compartment to the unstirred compartment |
| DiffR3uc | R3u → R3c | Diffusion of $RuBP^{3-}$ from the unstirred compartment to the Rubisco compartment |
| DiffP3eu | P3e → P3u | Diffusion of $PGA^{3-}$ from the external compartment to the unstirred compartment |
| DiffP3uc | P3u → P3c | Diffusion of $PGA^{3-}$ from the unstirred compartment to the Rubisco compartment |
| DiffP2eu | P2e → P2u | Diffusion of $PGA^{2-}$ from the external compartment to the unstirred compartment |
| DiffP2uc | P2u → P2c | Diffusion of $PGA^{2-}$ from the unstirred compartment to the Rubisco compartment |
| protonRe | R4e + He → R3e | Protonation of RuBP in the external compartment |
| unprotonRe | R3e → R4e + He | Deprotonation of RuBP in the external compartment |
| protonRu | R4u + Hu → R3u | Protonation of RuBP in the unstirred compartment |
| unprotonRu | R3u → R4u + Hu | Deprotonation of RuBP in the unstirred compartment |
| protonRc | R4c + Hc → R3c | Protonation of RuBP in the Rubisco compartment |
| unprotonRc | R3c → R4c + Hc | Deprotonation of RuBP in the Rubisco compartment |
| protonPe | P3e + He → P2e | Protonation of PGA in the external compartment |
| unprotonPe | P2e → P3e + He | Deprotonation of PGA in the external compartment |
| protonPu | P3u + Hu → P2u | Protonation of PGA in the unstirred compartment |
| unprotonPu | P2u → P3u + Hu | Deprotonation of PGA in the unstirred compartment |
| protonPc | P3c + Hc → P2c | Protonation of PGA in the Rubisco compartment |
| unprotonPc | P2c → P3c + Hc | Deprotonation of PGA in the Rubisco compartment |

**Table S2. COPASI model reaction species**

| Name | Units | Description |
|---|---|---|
| Be | $mol/m^3$ | $HCO_3^-$ in the external compartment |
| Ce | $mol/m^3$ | $CO_2$ in the external compartment |
| Oe | $mol/m^3$ | $O_2$ in the external compartment |
| He | $mol/m^3$ | $H^+$ in the external compartment |
| R4e | $mol/m^3$ | $RuBP^{4-}$ in the external compartment |
| R3e | $mol/m^3$ | $RuBP^{3-}$ in the external compartment |
| P3e | $mol/m^3$ | $PGA^{3-}$ in the external compartment |
| P2e | $mol/m^3$ | $PGA^{2-}$ in the external compartment |
| Bu | $mol/m^3$ | $HCO^{3-}$ in the unstirred compartment |
| Cu | $mol/m^3$ | $CO_2$ in the unstirred compartment |
| Ou | $mol/m^3$ | $O_2$ in the unstirred compartment |
| Hu | $mol/m^3$ | $H^+$ in the unstirred compartment |
| R4u | $mol/m^3$ | $RuBP^{4-}$ in the unstirred compartment |
| R3u | $mol/m^3$ | $RuBP^{3-}$ in the unstirred compartment |
| P3u | $mol/m^3$ | $PGA^{3-}$ in the unstirred compartment |
| P2u | $mol/m^3$ | $PGA^{2-}$ in the unstirred compartment |
| Bc | $mol/m^3$ | $HCO^{3-}$ in the Rubisco compartment |
| Cc | $mol/m^3$ | $CO_2$ in the Rubisco compartment |
| Oc | $mol/m^3$ | $O_2$ in the Rubisco compartment |
| Hc | $mol/m^3$ | $H^+$ in the Rubisco compartment |
| R4c | $mol/m^3$ | $RuBP^{4-}$ in the Rubisco compartment |
| R3c | $mol/m^3$ | $RuBP^{3-}$ in the Rubisco compartment |
| P3c | $mol/m^3$ | $PGA^{3-}$ in the Rubisco compartment |
| P2c | $mol/m^3$ | $PGA^{2-}$ in the Rubisco compartment |

**Table S3. COPASI model parameters; their units, and descriptions**

| Parameter | Units | Initial value(s) | Description |
|---|---|---|---|
| VolC | m³ | 0 | Condensate volume |
| VolU | m³ | 0 | Unstirred layer volume |
| SurfC | m² | 0 | Condensate surface area |
| SurfU | m² | 0 | Unstirred layer surface area |
| RadiusC | m | $1 \times 10^{-6}$ | Condensate radius |
| RadiusU | m | 0 | Unstirred layer radius |
| Kc | mol/m³ | Table 2 | Rubisco $K_M CO_2$ |
| Vc | 1/s | Table 2 | Rubisco carboxylation turnover rate |
| Ko | mol/m³ | Table 2 | Rubisco $K_M O_2$ |
| Vo | 1/s | Table 2 | Rubisco oxygenation turnover rate |
| Kr | mol/m³ | Table 2 | Rubisco $K_M RuBP$ |
| Vcarb | mol/m³/s | 0 | Maximum carboxylation velocity |
| Vox | mol/m³/s | 0 | Maximum oxygenation velocity |
| sites | mol/m³ | 10 | Rubisco compartment active site concentration |
| kr1prot | m³/(mol.s) | $5.01187 \times 10^{9}$ | RuBP protonation dissociation constant |
| kr2unprot | 1/s | $1 \times 10^{6}$ | RuBP deprotonation rate constant |
| kp1prot | m³/(mol.s) | $3.15 \times 10^{9}$ | PGA protonation rate constant |
| kp2unprot | 1/s | $1 \times 10^{6}$ | PGA deprotonation rate constant |
| PermcC | m/s | Table 1 | Rubisco compartment permeability to $CO_2$ |
| PermcB | m/s | Table 1 | Rubisco compartment permeability to $HCO_3^-$ |
| PermcO | m/s | Table 1 | Rubisco compartment permeability to $O_2$ |
| PermcH | m/s | Table 1 | Rubisco compartment permeability to $H^+$ |
| PermcR | m/s | Table 1 | Rubisco compartment permeability to RuBP species |
| PermcP | m/s | Table 1 | Rubisco compartment permeability to PGA species |
| PermuC | m/s | Table 1 | Unstirred layer permeability to $CO_2$ |
| PermuB | m/s | Table 1 | Unstirred layer permeability to $HCO_3^-$ |
| PermuO | m/s | Table 1 | Unstirred layer permeability to $O_2$ |
| PermuH | m/s | Table 1 | Unstirred layer permeability to $H^+$ |
| PermuR | m/s | Table 1 | Unstirred layer permeability to RuBP species |
| PermuP | m/s | Table 1 | Unstirred layer permeability to PGA species |
| CAe | unitless | Table 1 | Carbonic anhydrase catalysis factor in the external compartment |
| CAu | unitless | Table 1 | Carbonic anhydrase catalysis factor in the unstirred compartment |
| CAc | unitless | Table 1 | Carbonic anhydrase catalysis factor in the Rubisco compartment |
| CAk1e | 1/s | 0.05 | Carbonic anhydrase forward rate constant in the external compartment |
| CAk2e | 1/s | 100 | Carbonic anhydrase backward rate constant in the external compartment |
| CAk1u | 1/s | 0.05 | Carbonic anhydrase forward rate constant in the unstirred compartment |
| CAk2u | 1/s | 100 | Carbonic anhydrase backward rate constant in the unstirred compartment |
| CAk1c | 1/s | 0.05 | Carbonic anhydrase forward rate constant in the Rubisco compartment |
| CAk2c | 1/s | 100 | Carbonic anhydrase backward rate constant in the Rubisco compartment |
| Number | unitless | 1 | Number of Rubisco compartments |

### SI Figures

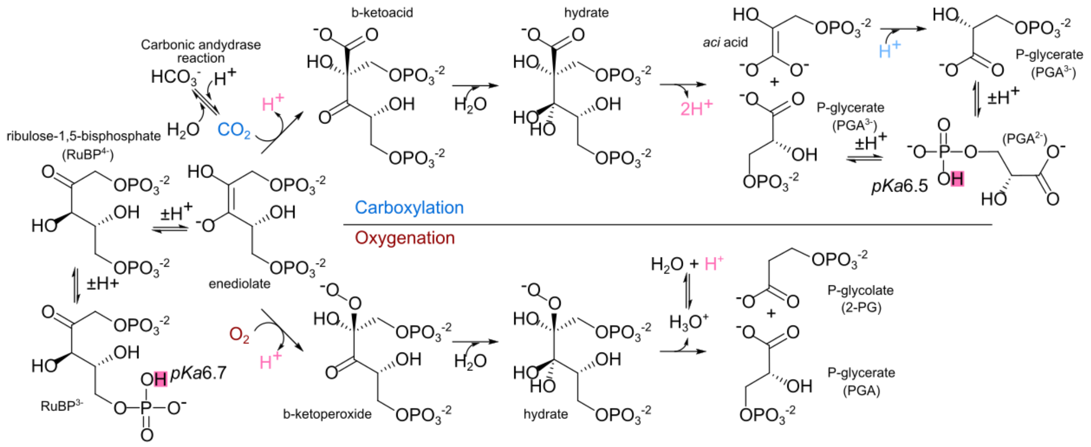


**Fig. S1. Rubisco reaction mechanism and the role of free protons**

In the action of Rubisco upon RuBP, a single proton is released (highlighted in pink) during conversion of the RuBP enediolate to the β-ketoacid (in the case of carboxylation) or the β-ketoperoxide (in the case of oxygenation). During the carboxylation process, an additional two protons (pink) are released from the hydrated gem-diol, while one is utilized (light blue) in the re-protonation of the aci-acid to yield a second molecule of PGA. Initial enediolate formation results from proton abstraction and therefore does not represent proton release or utilization. Two net protons are also released via the oxygenation reaction. Carbonic anhydrase (CA) catalyzes the interconversion of CO2 and HCO3-, utilizing free protons in the dehydration of HCO3-. Both RuBP and PGA can undergo protonation and deprotonation at physiological pH with phosphate group hydroxyl protons (pink squares) having pKa values of 6.7 and 6.5, respectively. RuBP and PGA pKa values were determined from software services provided by ChemAxon (chemaxon.com, Chemicalize). Figure adapted from Tcherkez, et al. (38).

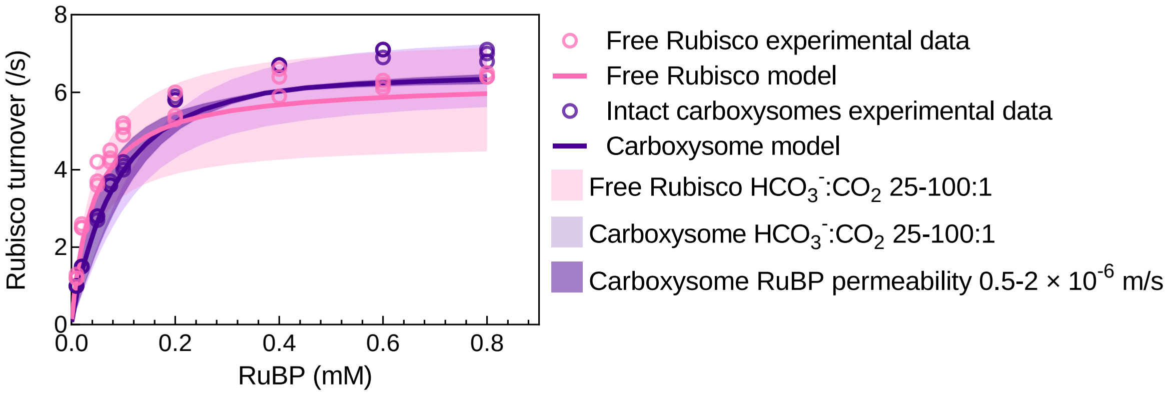


**Fig. S2. Application of the model to experimental data**

Experimental data were obtained for Rubisco turnover of the free *Cyanobium* Rubisco enzyme ('Free Rubisco experimental data' – *pink circles*) and the same enzyme in intact carboxysomes ('Intact carboxysomes experimental data' – *purple circles*) over a range of RuBP concentrations – data from a previous report (see Supplementary materials in reference 12). Typical free enzyme and carboxysome comparisons give rise to an increase in the apparent $K_{MRuBP}$ and $k_{cat}^{C}$ for intact carboxysomes compared with the naked enzyme in our hands. These shifts in Rubisco catalytic parameters observed in the experimental data are also apparent in the model, suggesting that catalytic parameters determined for the free enzyme are maintained within a carboxysome. Model output was generated using the kinetic properties of the free *Cyanobium* Rubisco enzyme (Table 2) with an initial [$HCO_3^-$] of 20 mM and pH 8.0 (as used in the experimental regime in 12), and the conditions for either the free enzyme or a carboxysome as outlined in Table 1. Carbonic anhydrase (CA) enzyme was absent from free Rubisco modelling, as was the case in experiments. For carboxysomes, CA was confined to the Rubisco compartment for modelling. Carboxysome size in the model was set to a radius of 5 × $10^{-8}$ m, to represent the size of carboxysomes from *Cyanobium* (12). *Solid pink* and *purple lines* are the model output for the free enzyme and carboxysomes respectively, using an $HCO_3^-$:$CO_2$ ratio of 50:1 in the model. *Light pink and light purple shaded regions* indicating the model output between $HCO_3^-$:$CO_2$ ratio ranges of 25 and 100:1 to account for experimental variability in the actual ratio in laboratory experiments. The model uses a ratio of 100:1 for most scenarios in this report. The *dark purple shaded area* indicates possible rates within the model when varying RuBP permeability within the carboxysome between 0.5 and 2.0 × $10^{-6}$ m/s. The model uses an RuBP permeability of 1 × $10^{-6}$ m/s for the carboxysome in this report.

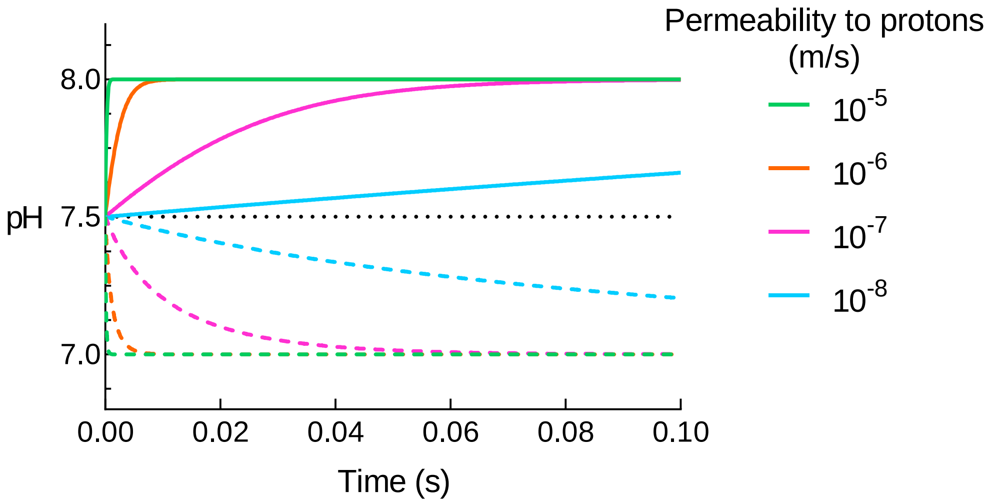


**Fig. S3 Time course of internal carboxysome pH response to changes in external pH**

Modelled equilibration of carboxysome internal pH upon exposure to relatively acidic or basic external environments. This experiment attempts to replicate that of Menon, Heinhorst, Shively and Cannon (24) in which purified carboxysomes of the proteobacterium *Halothiobacillus neapolitanus* were first equilibrated overnight in buffer at pH 7.5, then exposed to external buffer conditions of pH 7.0 or 8.0 in the absence of substrates (RuBP or $HCO_3^-$). In their experiments both broken and intact carboxysomes achieved stable pH values (as measured by the response of a carboxysomal pH-sensitive GFP-tagged Rubisco) within 20 milliseconds, implying free proton movement across the carboxysome shell. Here we simulate this experiment, modelling a 100 nm diameter carboxysome supplied with no substrates (eliminating internal proton production by Rubisco) and an initial internal pH of 7.5 (31.6 nM $H^+$). We investigated the time course response of carboxysomes with variable permeability to protons as the internal compartment equilibrated to external pH values of 7.0 (100 nM $H^+$; dashed lines) and 8.0 (10 nM $H^+$; solid lines). The COPASI (39) model was run in time course mode over 0.1 s with 0.0001 s intervals. Permeability of the Rubisco compartment to protons was varied from $10^{-8}$ to $10^{-5}$ m/s (colored lines). The results highlight that 'broken' carboxysomes (permeabilities approaching 1 m/s) should equilibrate with the external environment within nanosecond timescales, suggesting the methods used by Menon et al. may be limited by the timescale of pH-sensitive GFP responsiveness. If correct, this implies that carboxysomes have proton permeabilities which give rise to pH equilibration between 0 and 20 millisecond time-frames, consistent with permeabilities of $10^{-7}$ m/s and greater. In the modelling experiments presented here we use a proton permeability of $10^{-4}$ m/s for the carboxysome and assume a higher value of $10^{-2}$ m/s for Rubisco condensates (see SI Appendix, SI Methods).

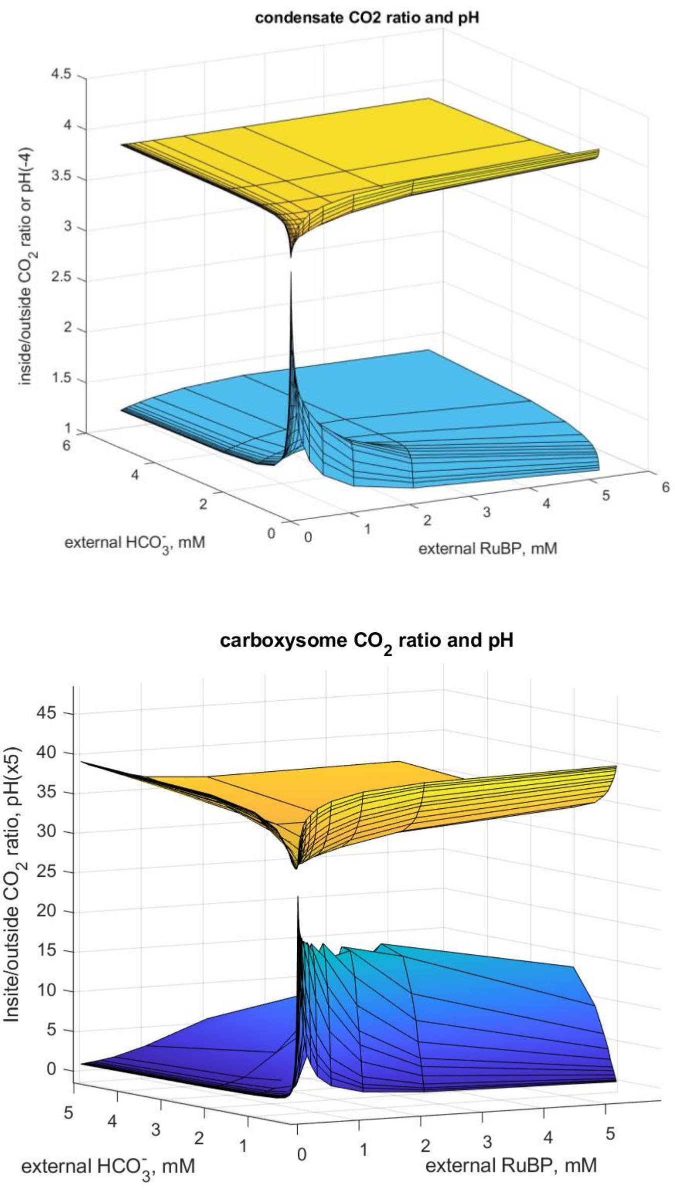


**Fig. S4. Condensates and carboxysomes display low pH and elevated $CO_2$ at low substrate concentrations**

Condensate (top) and carboxysome (bottom) relative $CO_2$ concentrations (*blue shaded surface*) and internal compartment pH (*yellow shaded surface*) as a function of both external $HCO_3^-$ and RuBP concentrations in the model. Compartment $CO_2$ is presented as the ratio of $CO_2$ inside the compartment to that outside (calculated as external $HCO_3^-$ concentration/100, see Methods). To place both $CO_2$ and pH on the same *z*-axis, in the top 3D-surface plot pH is the scale value +4 and for the lower plot pH is the scale value divided by 5.

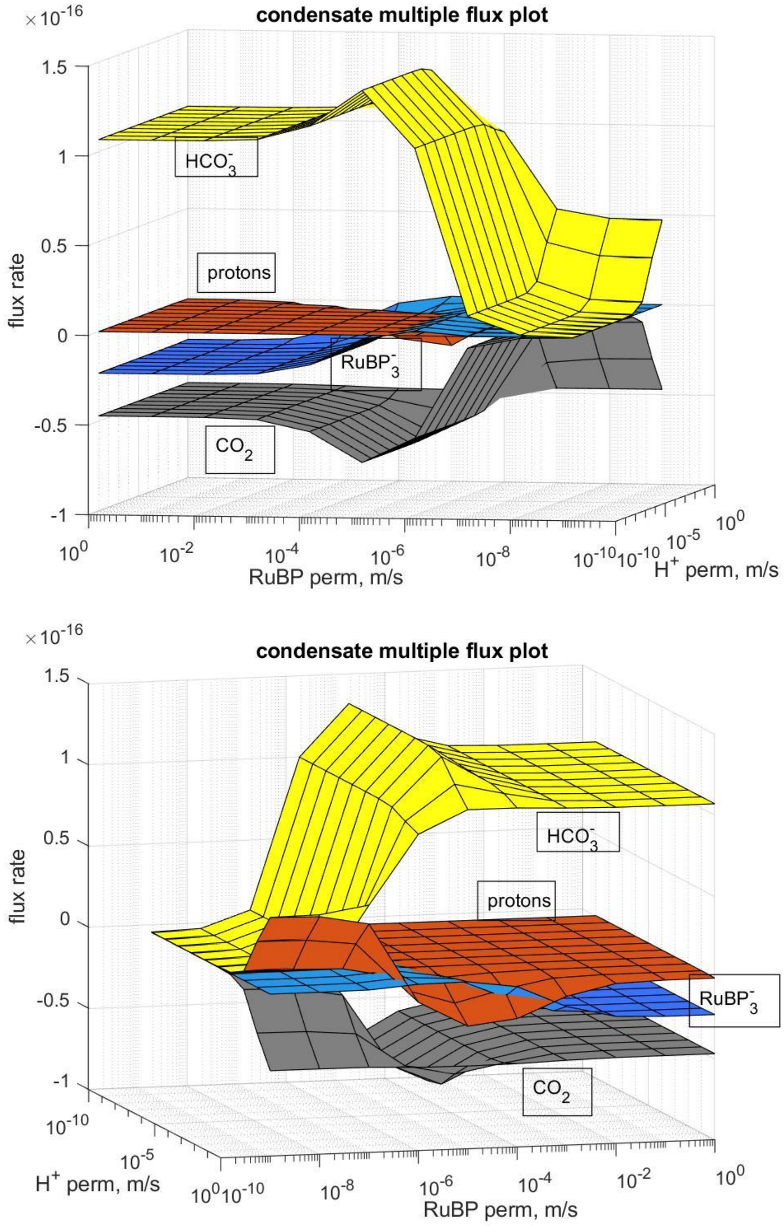


**Fig. S5. Condensate metabolite fluxes in response to variable proton and RuBP permeability**

3D surface plots of $HCO_3^-$ (*yellow*), protons (*red*), $CO_2$ (*grey*) and $RuBP^{3-}$ (*blue*) flux rates (mol/m$^3$/s) across the condensate/unstirred layer boundary using the COPASI model. The second plot is a rotation of the first plot around the *z*-axis to enable clear presentation of the data. Positive values represent an influx into the condensate while negative values represent an efflux.

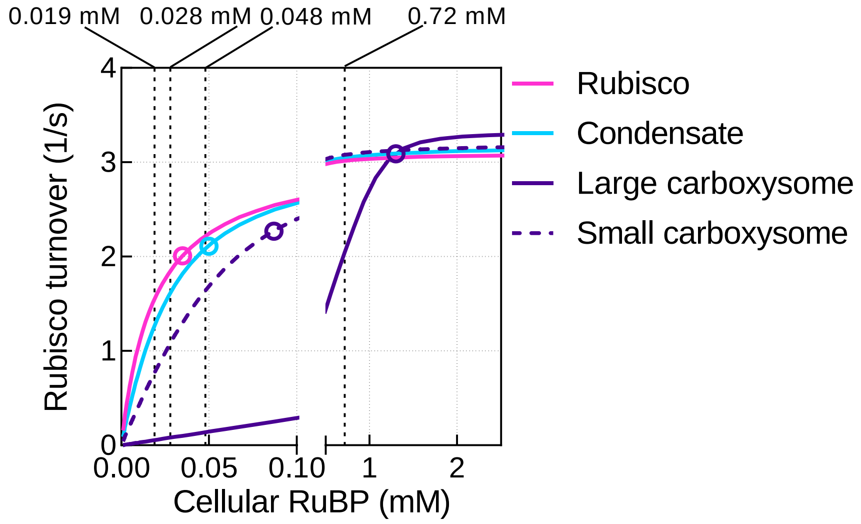


**Fig. S6. Change in the apparent *$K_M$RuBP* of Rubisco in a compartment**

RuBP response curves for free Rubisco (*pink line*), a Rubisco condensate (*blue line*) and both a large (1000 nm radius) carboxysome (*purple line*) and small (50 nm radius) carboxysome (*purple dashed line*) in the model, showing a shift in the apparent *$K_M$RuBP* values arising from decreased substrate permeability. Here the tobacco Rubisco is modelled as the free enzyme, a condensate, or a carboxysome as outlined in Table 1, using a *$K_M$RuBP* value of 0.018 mM (Table 2), at saturating $HCO_3^-$ (20 mM) at 20% (*v/v*) $O_2$. Michaelis-Menten enzyme kinetic parameters were subsequently estimated from the output carboxylation turnover rates using GraphPad Prism 9.0. Apparent *$K_M$RuBP* values for the free enzyme (0.019 mM), a condensate (0.028 mM), and a large (0.72 mM) and small carboxysome (0.048 mM) are shown above the plot for each curve. Using the model parameters outline in Table 1, condensation of Rubisco results in a modest rise in apparent *$K_M$RuBP* while encapsulation in a carboxysome increased the free enzyme value by four-fold in this instance. In each case carbonic anhydrase (CA) is modelled as present only in the Rubisco compartment in each case. Circles indicate Rubisco carboxylation rates at the RuBP concentrations used in ‘sub-saturating’ substrate modelling experiments.

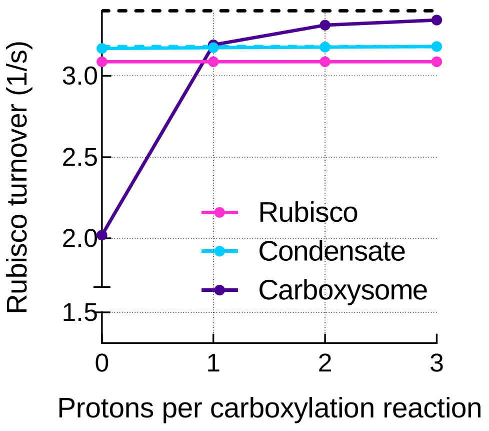


**Fig. S7. Maximum Rubisco carboxylation turnover rate as a function of net proton production**

Maximum Rubisco turnover rates calculated for free Rubisco, a Rubisco condensate and carboxysomes as a function of net proton production via the carboxylation reaction using the described model. Rates are calculated for saturating substrate conditions (5 mM RuBP, 20 mM $HCO_3^-$) with 0, 1, 2, or 3 protons produced as products of the carboxylation reaction. In each case carbonic anhydrase (CA) activity is confined only to the Rubisco compartment. Conditions describing the free enzyme, condensate or carboxysome are indicated in Table 1. Correct stoichiometry produces two protons as products of the carboxylation reaction. Note a net increase in maximum turnover rates in the condensate and carboxysome over the free enzyme when one or more protons are produced. In the absence of proton production, carboxysomal Rubisco function is limited by reduced $CO_2$ production via CA, dependent on proton diffusion into the compartment from the external volume. Maximum turnover rate of the tobacco Rubisco used in this modelling (3.4 1/s) is indicated by the *black dashed line*. Maximum turnover rate achieved for the condensate scenario at when 3 protons are produced (3.18 1/s) is indicated by the *blue dashed line*. Proton number was manufactured in the model by modifying the Rubisco carboxylation reaction stoichiometry to produce 0, 1, 2, or 3 protons. Data presented are for the tobacco Rubisco with parameters listed in Table 2 using typical modelling parameters as set out in Table 1.

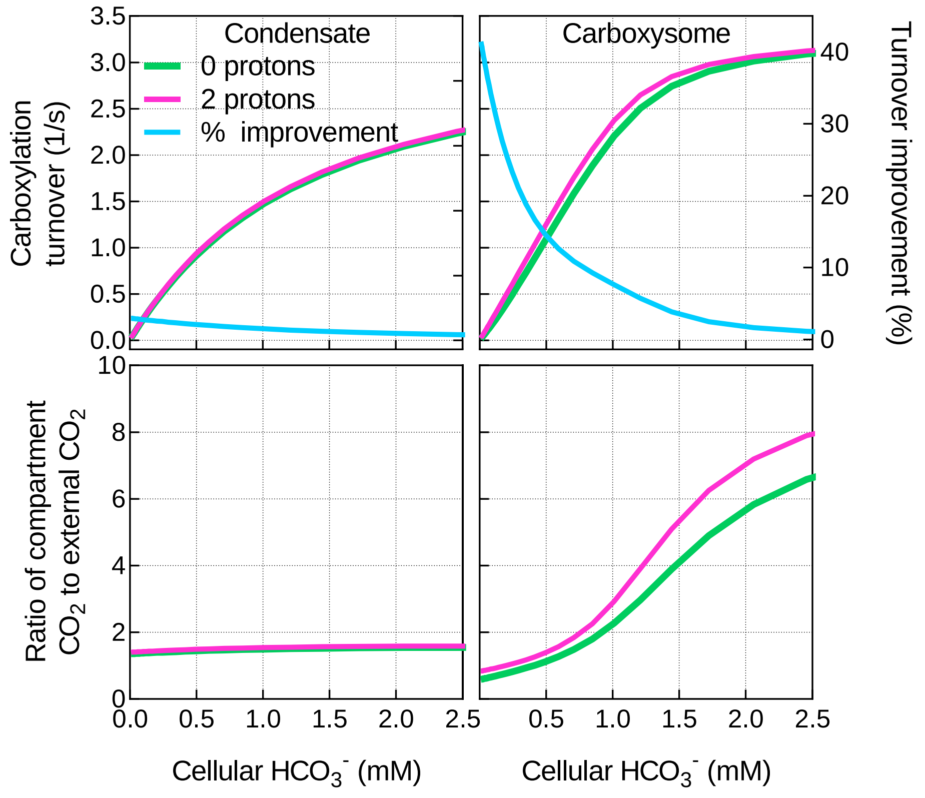


**Fig. S8. Protons derived from the Rubisco oxygenation reaction support carboxylation in a carboxysome**

Protons resulting from the oxygenation of RuBP within the model contribute to an improvement in carboxylation turnover (top panels) in a carboxysome (right panels) but less so in a condensate (left panels), especially at low $HCO_3^-$ supply. This improvement is diminished at higher $HCO_3^-$ concentrations as oxygenase is suppressed by increasing $CO_2$ supply. Protons arising from the oxygenation reaction result in additional carboxysome $CO_2$ production (bottom panels) via the internalized carbonic anhydrase (CA). Here we have run the model using the tobacco Rubisco (Table 2) with either zero or two protons arising from the oxygenation reaction in either a condensate or carboxysome using the parameters in Table 1. The COPASI (39) model was run in parameter scan mode, achieving steady-state values over the range of $HCO_3^-$ concentrations indicated.

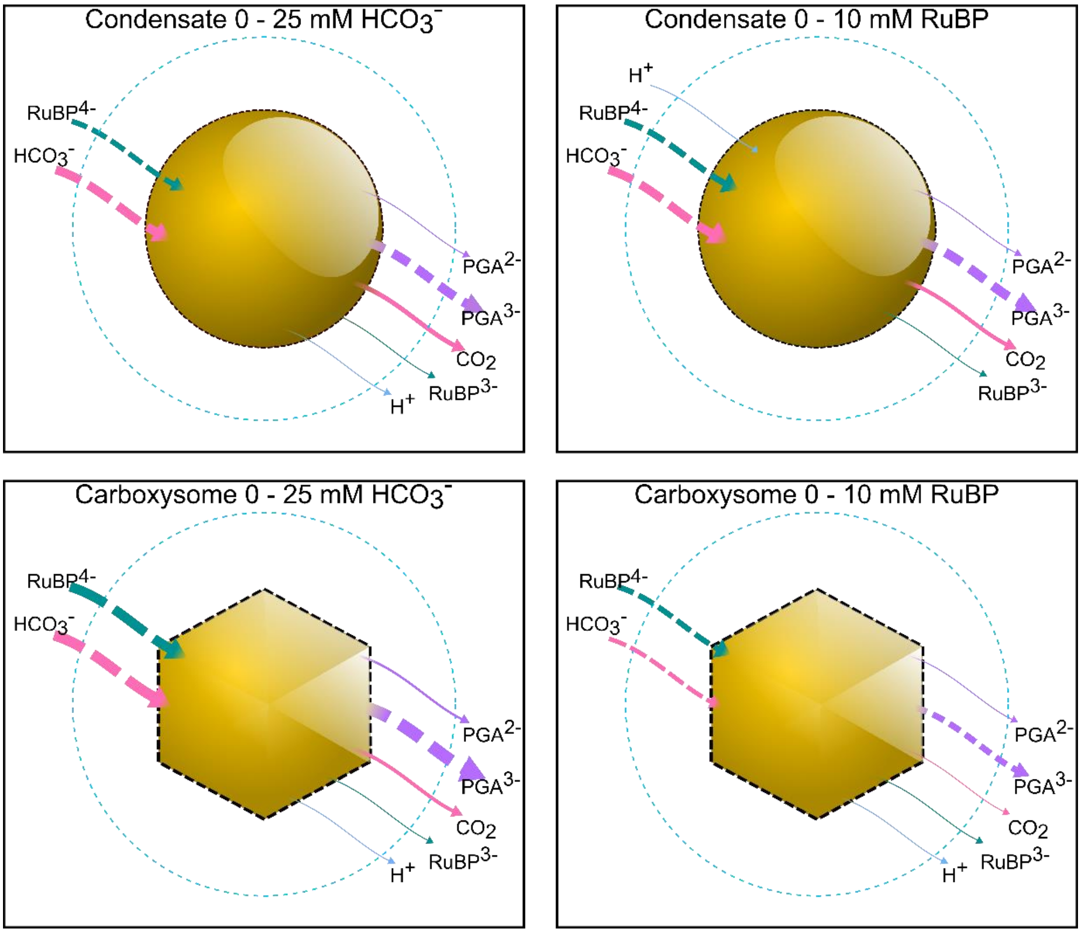


**Fig. S9. Summary of major net fluxes in condensates and carboxysomes.**

Averaged diffusional fluxes of chemical species across the condensate or carboxysome boundary over a range of $HCO_3^-$ and RuBP concentrations in the model. The thickness of each arrow is indicative of the relative net flux of that species and the direction indicates net flux into or out of the Rubisco compartment. Solid lines indicate species which are responsible for proton movement into or out of the compartment. Protons are carried by $RuBP^{3-}$, $PGA^{2-}$ and $CO_2$ (as the substrate required for free proton release via the CA hydration reaction). Net free $H^+$ fluxes (*solid light blue lines*) are extremely small and contributions to internal pH primarily arise through net fluxes of proton-carrier substrates (*solid green* [$RuBP^{3-}$], *solid purple* [$PGA^{2-}$], and *solid pink* [$CO_2$] *lines*). The deprotonated $RuBP^{4-}$ and $PGA^{3-}$ (*dashed green and purple lines*, respectively) are the substrate and product of Rubisco carboxylation, respectively, within the model. Data are average fluxes over the full data series for each condition as described in Fig. 5.

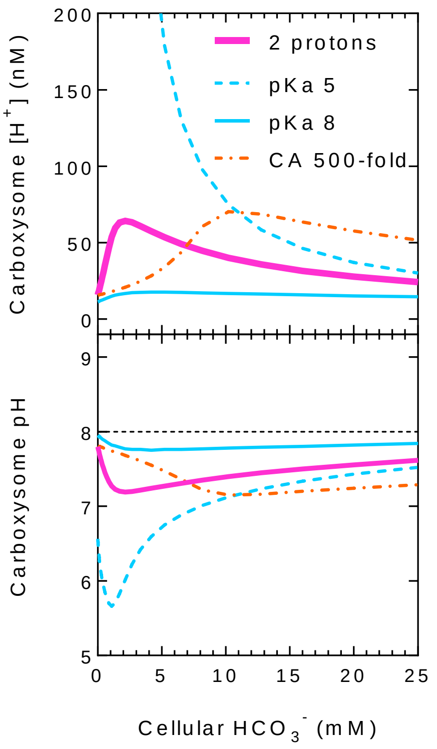


**Fig. S10. RuBP and PGA buffering, and CA activity, contribute to carboxysome pH control**

Proton concentration (top panel) and pH (lower panel) of carboxysomes in response to external $HCO_3^-$ supply in the model. Typical modelling scenarios (Table 1) allow for the production of 2 protons (*pink line*) from the Rubisco carboxylation reaction. This results in a net generation of free protons within the carboxysome, leading to enhanced dehydration of $HCO_3^-$ by carbonic anhydrase, production of $CO_2$ and higher rates of carboxylation by Rubisco. While $CO_2$ efflux from the carboxysome helps maintain moderate internal pH (Fig. 4), proton buffering by RuBP and PGA, and relative CA rate also contribute. Buffer capacity of RuBP and PGA is facilitated through protonation and deprotonation of functional groups with *pKa* values of 6.7 and 6.5 respectively (Fig. S1, Table S3). Manipulating these values within the model to *pKa*'s of 5 (*dashed blue line*; kr1prot and kp1prot set to $1 \times 10^8$) and 8 (*solid blue line*; kr1prot and kp1prot set to $1 \times 10^{11}$) for both molecules reveals that their buffering capacity is a major contributor to pH stability. While our typical model scenarios have carboxysomal CA activity at $10^5$-fold the background rate of conversion, modifying this to a lower rate (500-fold; *dot-dashed orange line*) contributes to lower internal free proton concentrations at low external $HCO_3^-$, yet still enables high carboxylation rates at saturating substrate concentrations (Fig. S11). This is due to the relative rates of proton production by the carboxylation and consumption by CA at low external $HCO_3^-$. The COPASI (39) model was run in parameter scan mode, achieving steady-state values over the indicated range of $HCO_3^-$ concentrations at 5 mM RuBP. Data presented are for the tobacco Rubisco (Table 2).

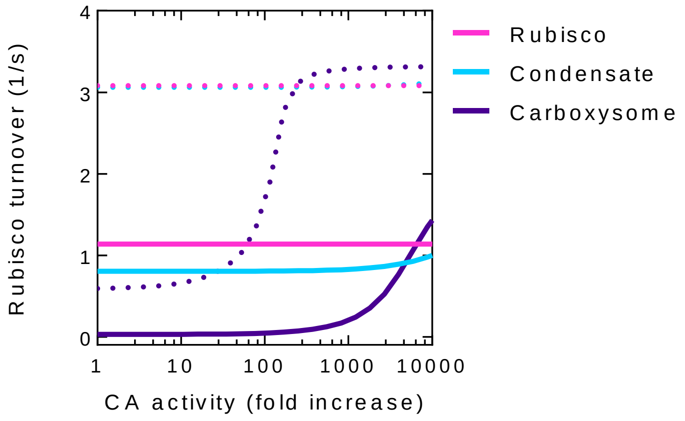


**Fig. S11. Compartment carboxylation rate is dependent upon CA activity**

Modulation of CA activity within the model affects carboxylation rate differentially in each compartment type. *Solid lines* represent free Rubisco and compartment carboxylation rates at sub-saturating substrate concentrations (1 mM $HCO_3^-$ and either 35, 50 or 1,300 µM RuBP for the free enzyme, condensate and carboxysome respectively). *Dotted lines* indicate rates achieved under saturating substrate conditions (20 mM $HCO_3^-$ and 5 mM RuBP). The COPASI (39) model was run in parameter scan mode, achieving steady-state values over the indicated range of CA activities in the Rubisco compartment at the substrate concentrations as indicated above. CA activity was confined to the Rubisco compartment only for this modelling. Data presented are for the tobacco Rubisco (Table 2).

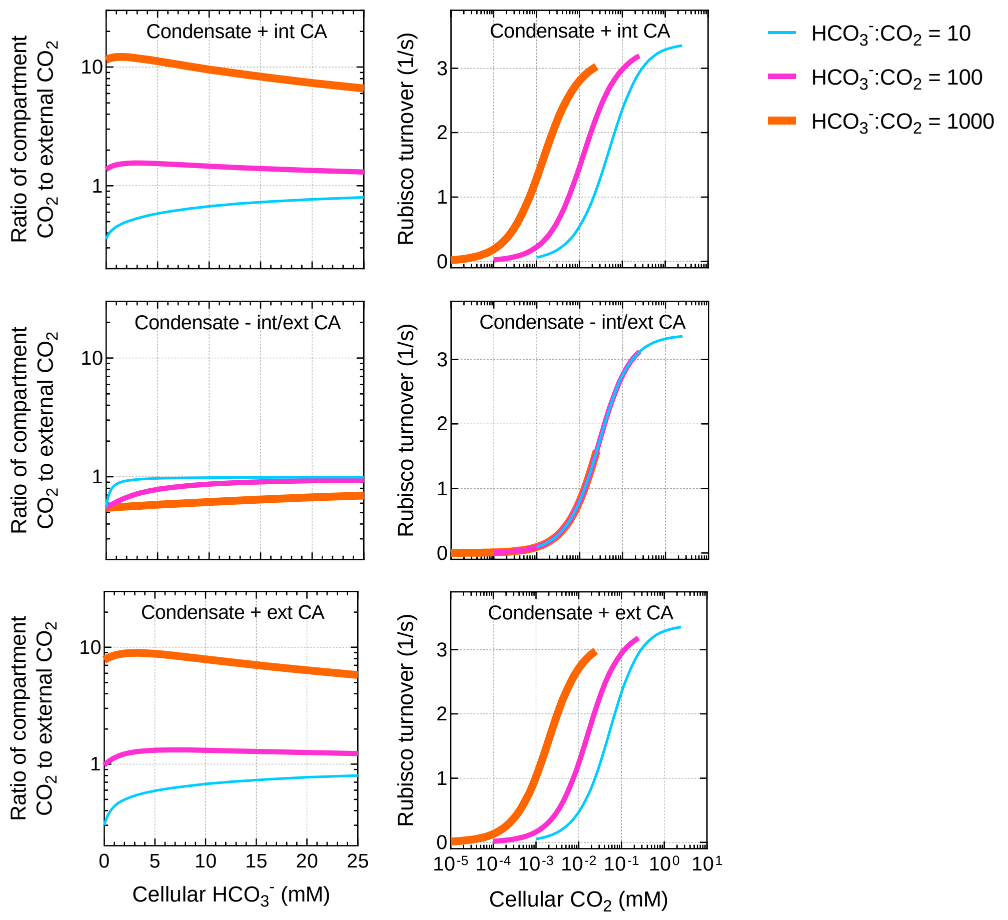


**Fig. S12 High external $HCO_3^-CO_2$ ratios enhance Rubisco compartment function**

Variation in the ratio of $HCO_3^-$:$CO_2$ in the external compartment (here analogous to a fixed cellular cytoplasm) results in changes in relative Rubisco compartment $CO_2$ (left panels, plotted over a range of cellular [$HCO_3^-$]) and carboxylation turnover rates (right panels, plotted on a log scale over a range of cellular [$CO_2$]). In the presence of either an internal carbonic anhydrase (CA; top panels – Condensate + int CA) or an external CA (bottom panels – Condensate + ext CA – modelled in the unstirred layer [Table 1]) associated with a Rubisco condensate, elevation of the relative cytoplasmic ratio of $HCO_3^-$:$CO_2$ leads to both enhanced relative $CO_2$ within the condensate and higher Rubisco turnover rates relative to the cellular [$CO_2$]. This is because Rubisco condensation, in the presence of an internal or external CA, can lead to enhanced conversion of $HCO_3^-$ to $CO_2$ in the Rubisco compartment. A decrease in the ratio of $HCO_3^-$:$CO_2$ in the cytoplasm (e.g. $HCO_3^-$:$CO_2$ = 10) diminishes the supply of $HCO_3^-$ to the Rubisco compartment and less $CO_2$ is therefore generated at the site of fixation. A cell with no CA, either in the cytoplasm or in the condensate (middle panels – Condensate - int/ext CA) cannot generate additional $CO_2$ within the condensate, such that Rubisco turnover rates are entirely dependent on $CO_2$ supply from the cytoplasm. This is the prime reason that condensation is unlikely to have evolved in the absence of a cellular CA (Fig. 6). Modelled scenarios in this report utilize a $HCO_3^-$:$CO_2$ ratio of 100 (*pink lines*). Values approaching a ratio of 10 (*blue lines*) may occur in the absence of cytoplasmic CA in combination with the preferential diffusion of $CO_2$ across the cell membrane. Values approaching a ratio of 1000 (*orange lines*) may occur in the advent of active $HCO_3^-$ transport and subsequent loss of CA from the cytoplasm and/or movement to the condensate. Modelled here are Rubisco compartment $CO_2$/external compartment $CO_2$ ratios and Rubisco carboxylation turnover rates for a condensate using the parameters in Table 1. The 'external' compartment $HCO_3^-$:$CO_2$ ratio (colored lines) was varied for each scenario. The COPASI (39) model was run in parameter scan mode, achieving steady-state values over the indicated range of $HCO_3^-$ concentrations at 5 mM RuBP. $CO_2$ in the external compartment (cellular $CO_2$) was determined from the $HCO_3^-$:$CO_2$ ratio in each scenario at each [$HCO_3^-$]. Data presented are for the tobacco Rubisco (Table 2).

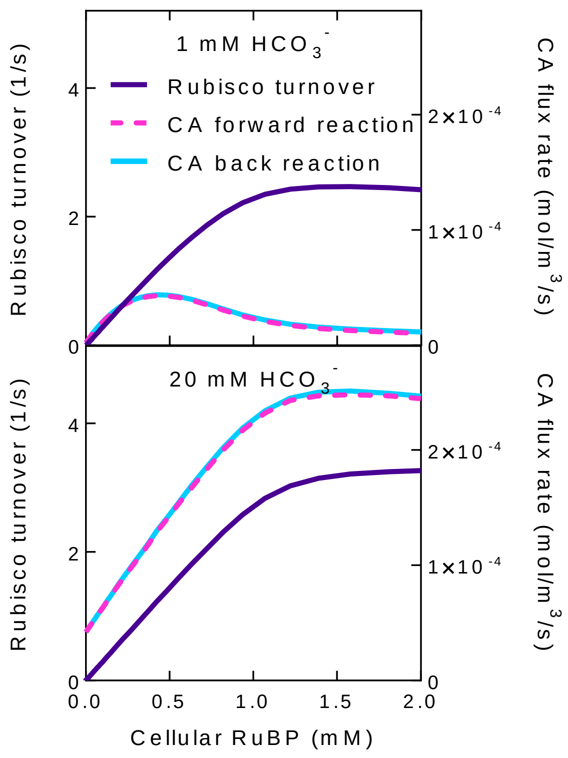


**Fig. S13. Carboxysome CA activity is dependent on Rubisco activity**

Carbonic anhydrase flux rates are closely coupled to Rubisco activity through the provision of internal protons by the Rubisco reaction. This is demonstrated by the response of both CA and Rubisco to RuBP. Here the COPASI (39) model was run in parameter scan mode, achieving steady-state values over the indicated range of [RuBP] in the carboxysome at either sub-saturating (1 mM) or saturating (20 mM) $HCO_3^-$. CA activity was confined to the carboxysome for this modelling. Data presented are for the tobacco Rubisco (Table 2).

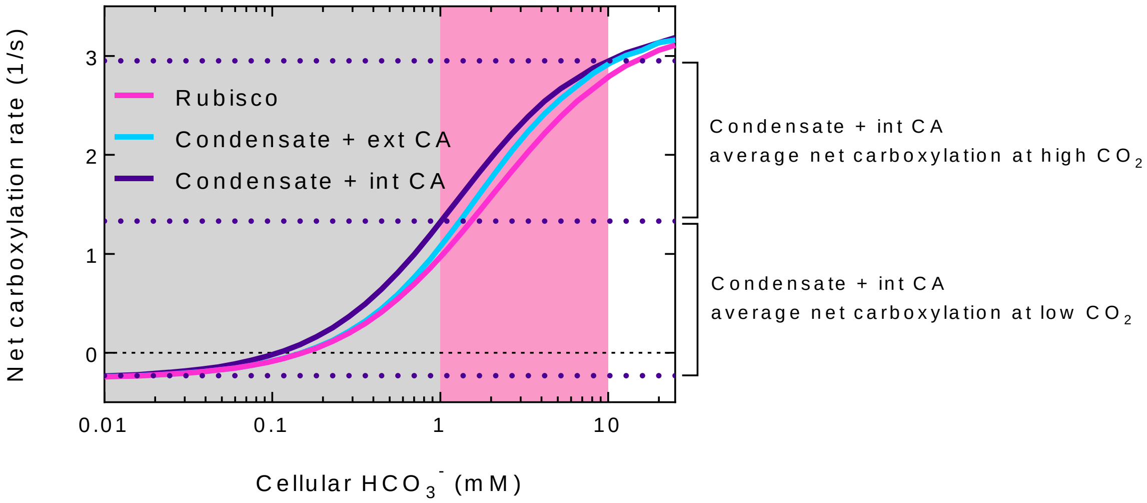


**Fig. S14. Method for estimating relative fitness of evolutionary intermediates from free Rubisco to contemporary carboxysomes**

Net carboxylation rates for potential evolutionary intermediates were calculated as the Rubisco carboxylation turnover minus ½ oxygenation turnover (1/s) over a range of $HCO_3^-$ concentrations in the model, assuming the same photorespiratory cost in all systems. In order to determine relative fitness of one evolutionary intermediate over another at high or low $CO_2$, net carboxylation rates were averaged for each intermediate over the range 0.01 – 1 mM $HCO_3^-$ (for low $CO_2$) and 1 – 10 mM $HCO_3^-$ (for high $CO_2$). The average net carboxylation rates for each intermediate at different $CO_2$ ranges are then compared to assess relative fitness under those conditions. In this example, relative rates of the free tobacco enzyme, a tobacco Rubisco condensate with an external carbonic anhydrase (Condensate + ext CA), or the tobacco Rubisco condensate with an internal CA (Condensate + int CA), are compared. The *horizontal dotted lines* set the limits for the net carboxylation rate ranges which are averaged for the Condensate + int CA condition under the low and high $CO_2$ conditions; the best performing evolutionary intermediate in this scenario. The Condensate + ext CA and free Rubisco conditions, however, suggest very little difference between each other over the low $CO_2$ range, yet the Condensate + ext CA has a greater average net carboxylation rate at high $CO_2$. The examples presented here were calculated under a 20% (*v/v*) $O_2$ atmosphere and the appropriate parameters for each scenario are indicated in Table 1.

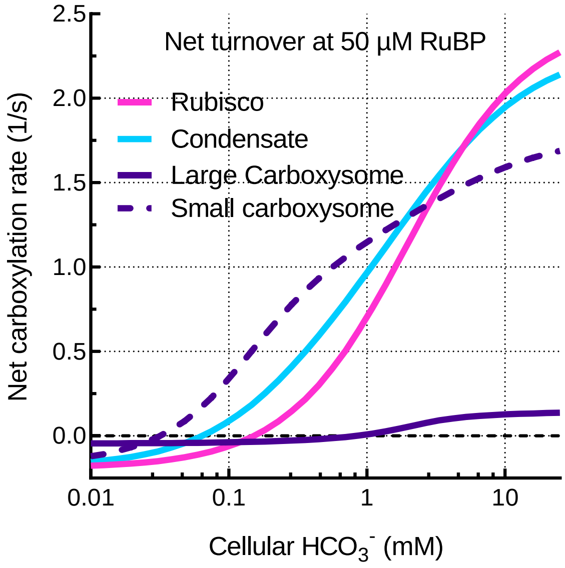


**Fig. S15. Carboxylation turnover at 50 µM RuBP**

Net carboxylation turnover rates of free Rubisco (*pink line*), a Rubisco condensate (*light blue line*), large (1000 nm radius) carboxysomes (*purple line*), and small (50 nm radius) carboxysomes (*purple dashed line*) at low RuBP (50 µM) in the model. Net carboxylation turnover rates were calculated as carboxylation minus ½ oxygenation turnover, assuming the same photorespiratory cost to carboxylation each case. Presented here are net carboxylation turnover rates for the tobacco enzyme (Table 2), modelled using the parameters for each scenario as in Table 1. Note that the condensate scenario here performs better than the free enzyme at low $HCO_3^-$ supply within the model, while the large carboxysome (which outperforms both the free enzyme and condensate under saturating conditions; Fig. S7) performs poorly. Very small carboxysomes have an additional performance benefit. This result supports the notion that Rubisco condensates and small carboxysomes have a performance advantage under light-limited and low $C_i$ conditions, whereas a large carboxysome system is unlikely to be successful. Smaller carboxysomes may have a net advantage over larger carboxysomes, driven by relatively lower RuBP diffusional limitations, but improved $CO_2$ fixation rates compared with condensates. Model conditions assume carbonic anhydrase (CA) is present in only the Rubisco compartment.

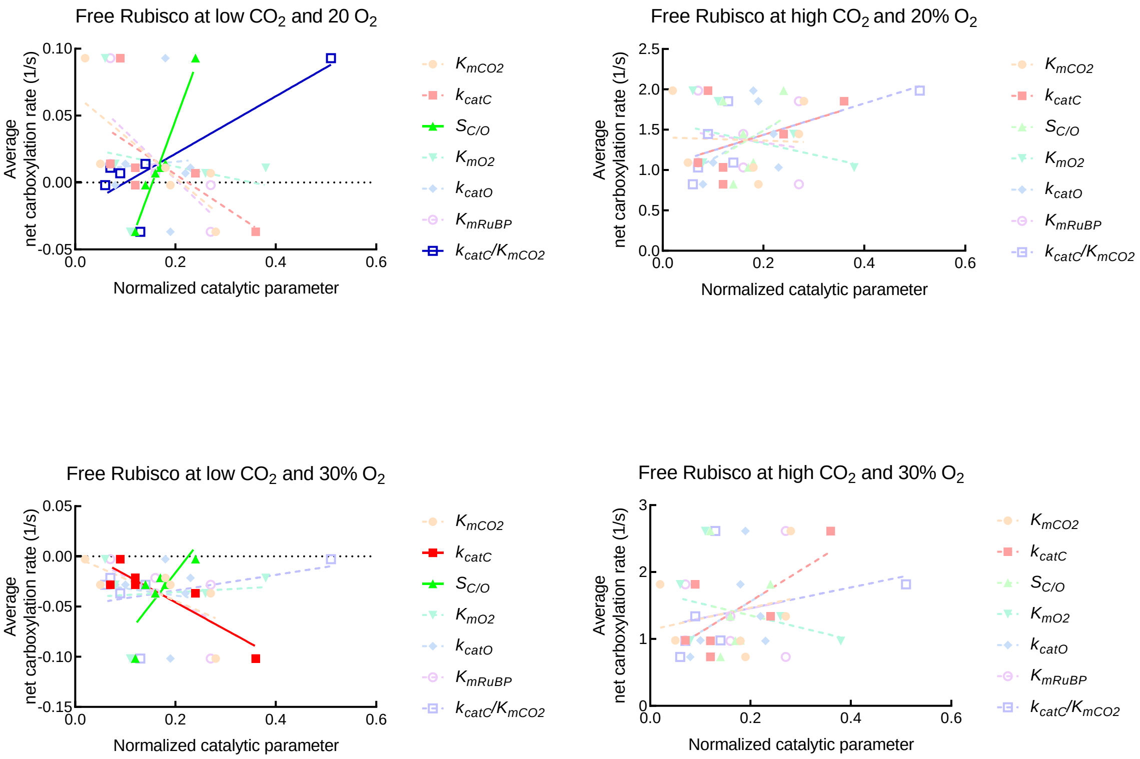


**Fig. S16. Correlation plots – free Rubisco under simulated varied atmospheres**

Correlation plots of average net carboxylation turnover rates (1/s) of different Rubisco enzymes and their catalytic parameters simulated as the free enzyme in the model. Here, the catalytic parameters of each enzyme have been normalized to the maximum value for that parameter across the enzymes analyzed (Table 2) for simplicity. Pearson correlations were calculated between average net carboxylation rates over the ranges of $CO_2$ and $O_2$ indicated (Fig. S14), and the value of each Rubisco catalytic parameter (*SI Appendix, SI* dataset S1). Parameters showing correlations at $p$ < 0.05 are plotted in *bold colors* with *solid lines*. Parameters showing correlations at $p$ > 0.05 are shown in *pale colors* with *dashed lines*.

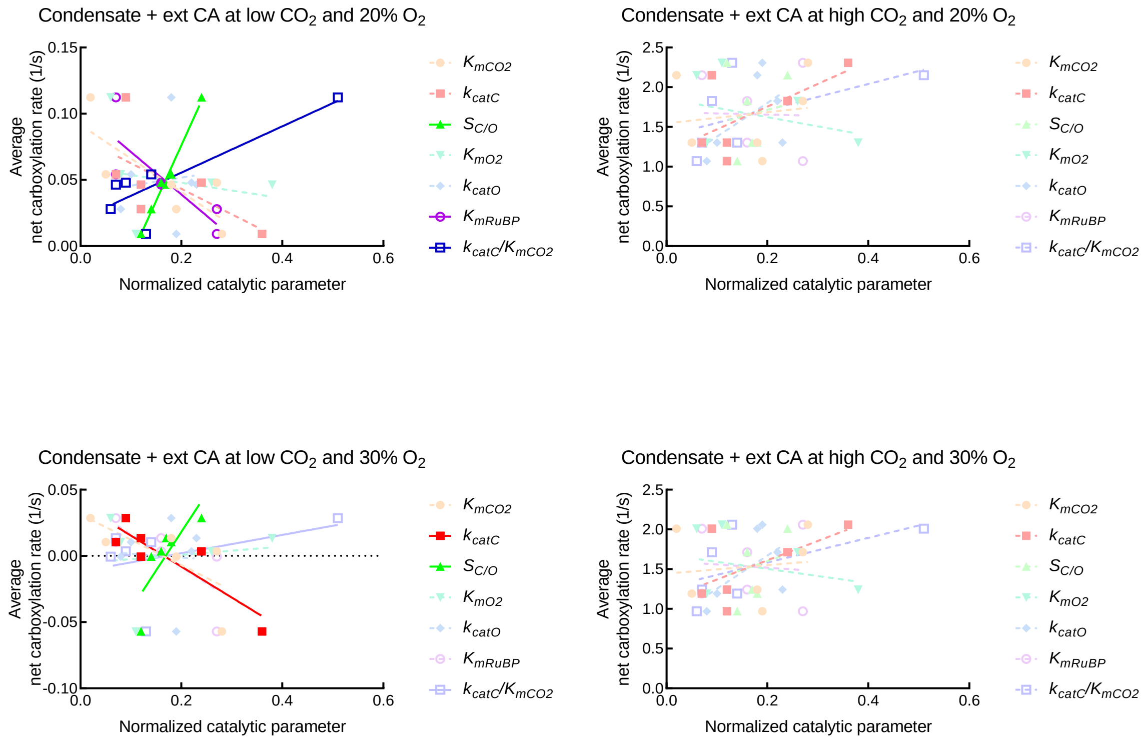


**Fig. S17. Correlation plots – Rubisco condensate with an external CA under simulated varied atmospheres**

Correlation plots of average net carboxylation turnover rates (1/s) of different Rubisco enzymes and their catalytic parameters simulated as a Rubisco condensate with an external CA in the model. Here, the catalytic parameters of each enzyme have been normalized to the maximum value for that parameter across the enzymes analyzed (Table 2) for simplicity. Pearson correlations were calculated between average net carboxylation rates over the ranges of $CO_2$ and $O_2$ indicated (Fig. S14), and the value of each Rubisco catalytic parameter (*SI Appendix, SI* dataset S1). Parameters showing correlations at $p < 0.05$ are plotted in *bold colors* with *solid lines*. Parameters showing correlations at $p > 0.05$ are shown in *pale colors* with *dashed lines*.

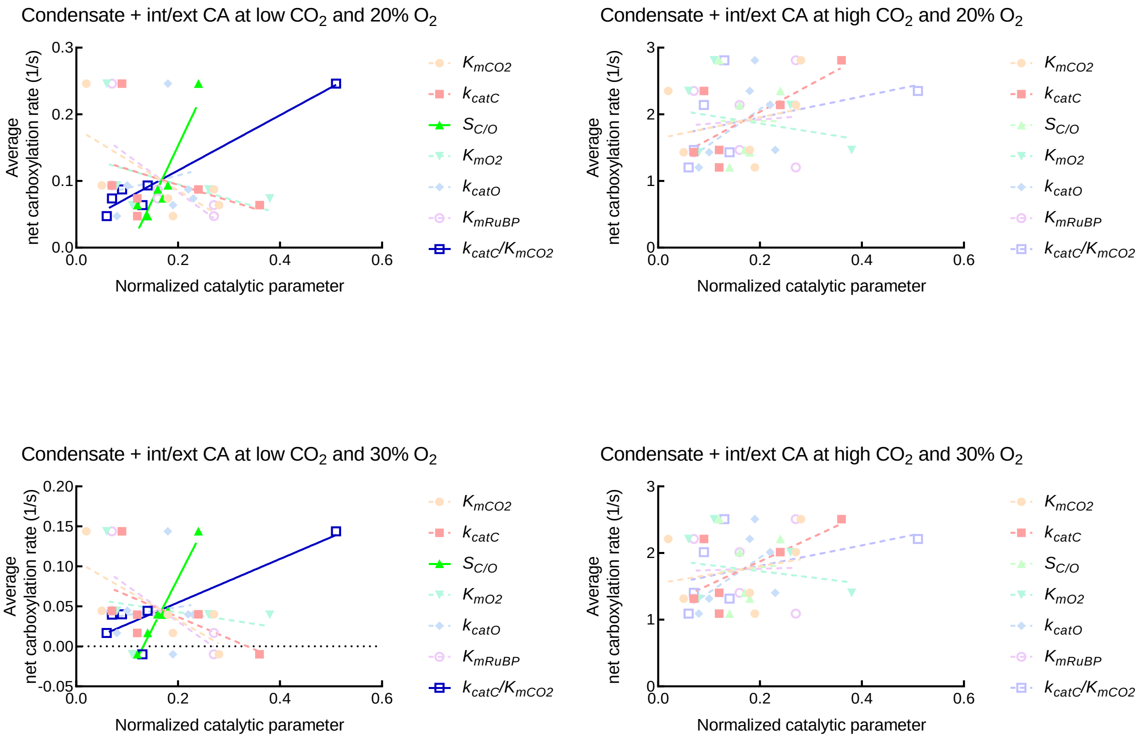


**Fig. S18. Correlation plots – Rubisco condensate with internal and external CA under simulated varied atmospheres**

Correlation plots of average net carboxylation turnover rates (1/s) of different Rubisco enzymes and their catalytic parameters simulated as a Rubisco condensate with both internal and external CA in the model. Here, the catalytic parameters of each enzyme have been normalized to the maximum value for that parameter across the enzymes analyzed (Table 2) for simplicity. Pearson correlations were calculated between average net carboxylation rates over the ranges of $CO_2$ and $O_2$ indicated (Fig. S14), and the value of each Rubisco catalytic parameter (*SI Appendix, SI* dataset S1). Parameters showing correlations at $p < 0.05$ are plotted in *bold colors* with *solid lines*. Parameters showing correlations at $p > 0.05$ are shown in *pale colors* with *dashed lines*.

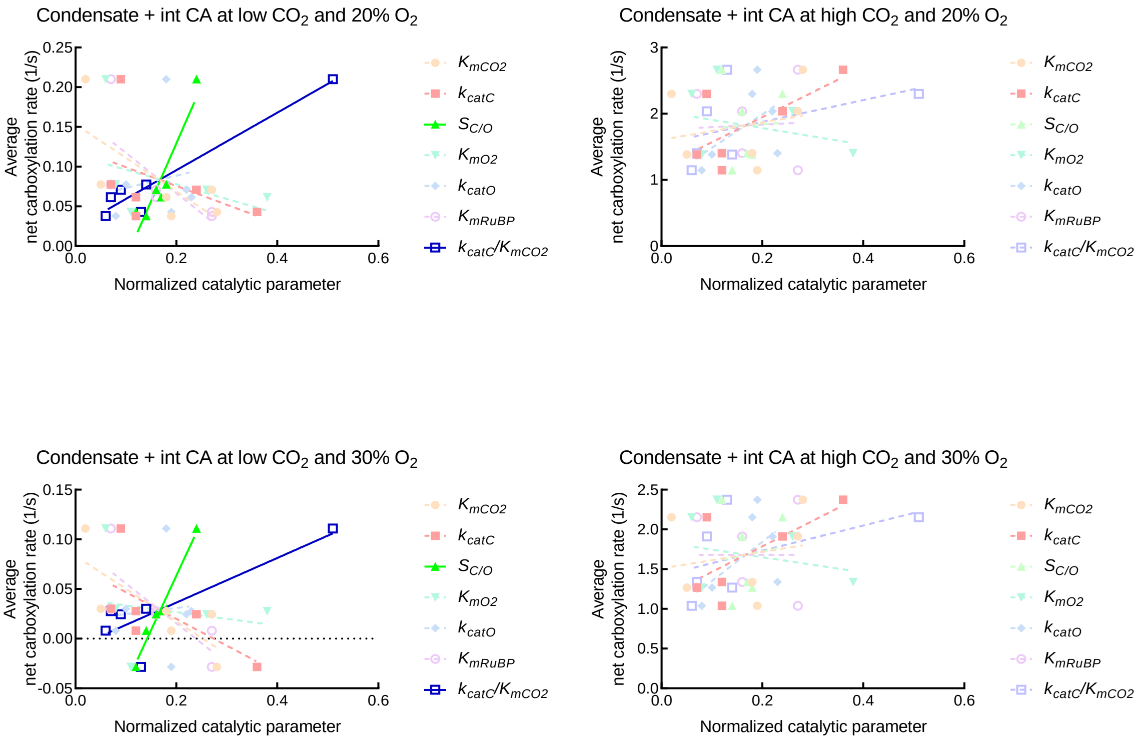


**Fig. S19. Correlation plots – Rubisco condensate with internal CA under simulated varied atmospheres**

Correlation plots of average net carboxylation turnover rates (1/s) of different Rubisco enzymes and their catalytic parameters simulated as a Rubisco condensate with internal CA in the model. Here, the catalytic parameters of each enzyme have been normalized to the maximum value for that parameter across the enzymes analyzed (Table 2) for simplicity. Pearson correlations were calculated between average net carboxylation rates over the ranges of $CO_2$ and $O_2$ indicated (Fig. S14), and the value of each Rubisco catalytic parameter (*SI Appendix, SI* dataset S1). Parameters showing correlations at $p < 0.05$ are plotted in *bold colors* with *solid lines*. Parameters showing correlations at $p > 0.05$ are shown in *pale colors* with *dashed lines*.

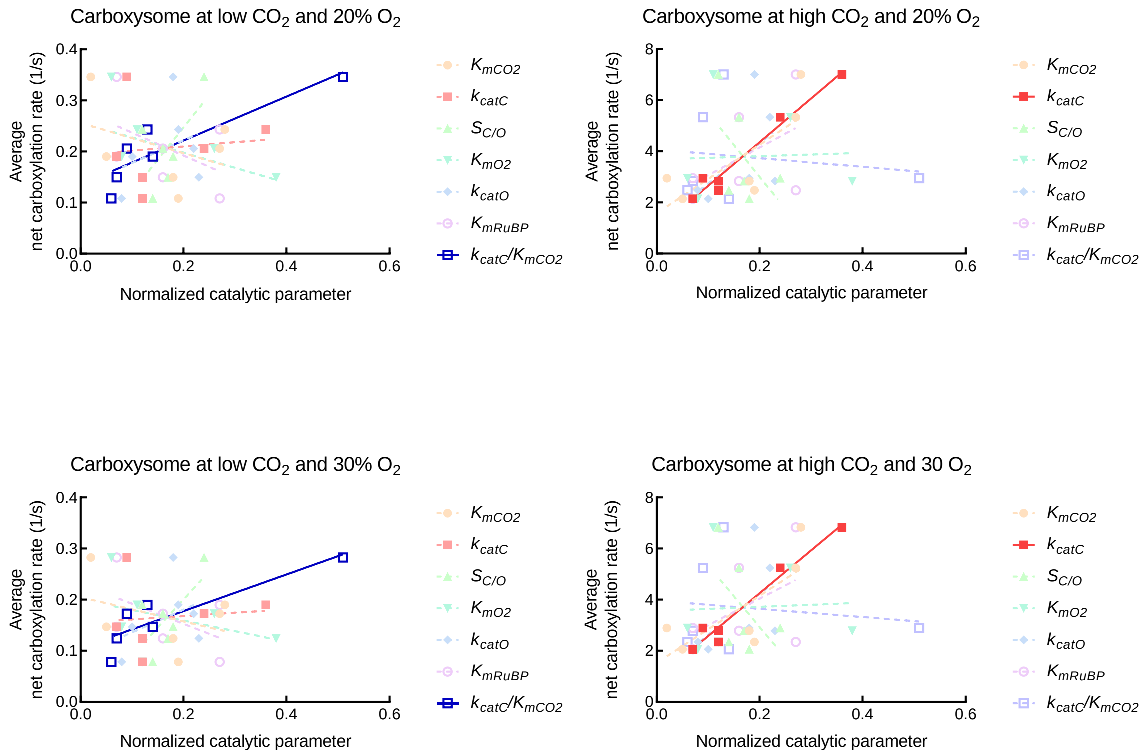


**Fig. S20. Correlation plots – Large contemporary carboxysome under simulated varied atmospheres**

Correlation plots of average net carboxylation turnover rates (1/s) of different Rubisco enzymes and their catalytic parameters simulated as a contemporary carboxysome (encapsulated Rubisco with an internal CA, , radius 1 × $10^{-6}$ m) in the model. Here, the catalytic parameters of each enzyme have been normalized to the maximum value for that parameter across the enzymes analyzed (Table 2) for simplicity. Pearson correlations were calculated between average net carboxylation rates over the ranges of $CO_2$ and $O_2$ indicated (Fig. S14), and the value of each Rubisco catalytic parameter (*SI Appendix, SI* dataset S1). Parameters showing correlations at $p < 0.05$ are plotted in *bold colors* with *solid lines*. Parameters showing correlations at $p > 0.05$ are shown in *pale colors* with *dashed lines*.

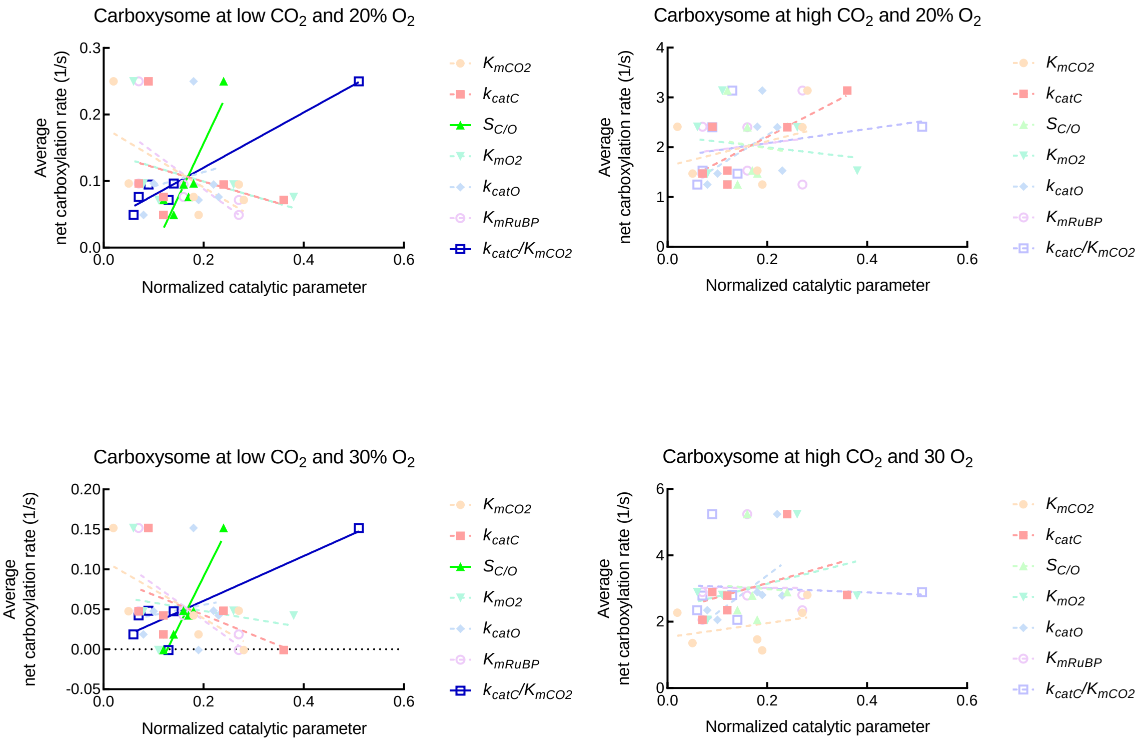


**Fig. S21 Correlation plots – Small contemporary carboxysome under simulated varied atmospheres**

Correlation plots of average net carboxylation turnover rates (1/s) of different Rubisco enzymes and their catalytic parameters simulated as a small contemporary carboxysome (encapsulated Rubisco with an internal CA, radius $5 \times 10^{-8}$ m) in the model. Here, the catalytic parameters of each enzyme have been normalized to the maximum value for that parameter across the enzymes analyzed (Table 2) for simplicity. Pearson correlations were calculated between average net carboxylation rates over the ranges of $CO_2$ and $O_2$ indicated (Fig. S14), and the value of each Rubisco catalytic parameter (*SI Appendix, SI* dataset S2). Parameters showing correlations at $p < 0.05$ are plotted in *bold colors* with *solid lines*. Parameters showing correlations at $p > 0.05$ are shown in *pale colors* with *dashed lines*.

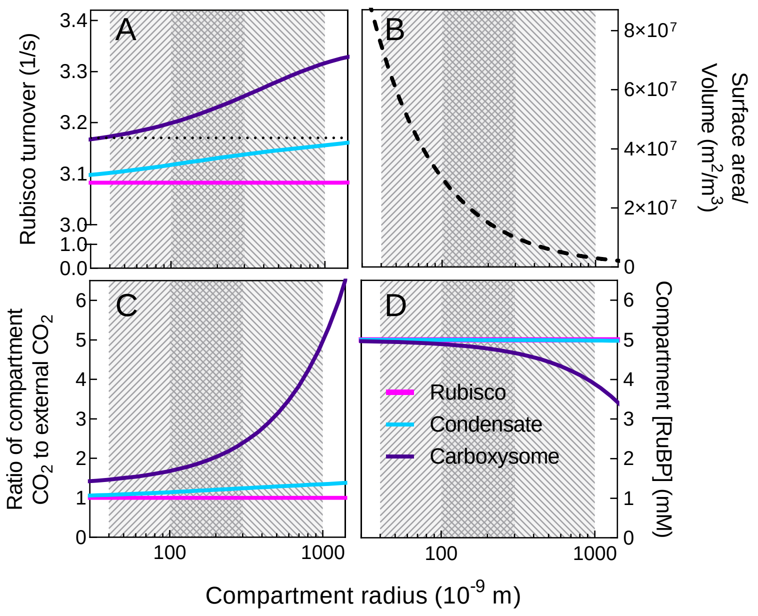


**Fig. S22 Model responses to condensate and carboxysome size**

Rubisco carboxylation turnover rate (A) increases by ≈4% for a carboxysome (*purple lines*) and ≈2% for a condensate (*blue lines*) over a broad range of compartment sizes ranging from very small carboxysomes (~5 x $10^{-8}$ m; left-most hatched region of plots) to large pyrenoids (~1 x $10^{-6}$ m; right-most hatched region of plots). Carboxysomes and pyrenoids exist in a wide size range with some overlap (cross-hatched region of plots). Assuming each compartment type as a sphere, the surface area/volume (*dashed black line*) decreases significantly with increasing size (B), leading to changes in relative diffusional flux of substrates to Rubisco. Under the conditions modelled here, the ratio of Rubisco compartment $CO_2$ to external $CO_2$ is roughly three times greater in large carboxysomes compared with small carboxysomes, while the differences are not as pronounced for variable condensate sizes (C). The high $CO_2$ in large carboxysomes leads to a decrease in compartment [RuBP] due to greater consumption rates by Rubisco (D). Modelling conditions are at saturating $HCO_3^-$ (20 mM) and RuBP (5 mM), using the tobacco Rubisco (Table 2). The free enzyme (*pink lines*), which is unaffected by size, is shown as a reference. Responses at each size, for each compartment type, is also dependent upon substrate concentration (Fig. S4).

**SI Equations**

**ODEs used to construct the modelling of condensates and carboxysomes. Parameters used in the equations are listed in SI Table S1.**

$$
\begin{aligned}
V_{\text{unstirred}} &= \text{Values[VolU].InitialValue} \\
V_{\text{condensate}} &= \text{Values[VolC].InitialValue} \\
[\text{Ce}] &= \frac{[\text{Be}]}{100} \\
\frac{\mathrm{d}([\text{R4e}] \cdot V_{\text{External}})}{\mathrm{d}t} &= -\left(\frac{\text{PermuR} \cdot \text{SurfU} \cdot ([\text{R4e}] - [\text{R4u}])}{\text{vol}_{(\text{DiffR4eu})}}\right) \\
&\quad - V_{\text{External}} \cdot (\text{kr1prot} \cdot [\text{R4e}] \cdot [\text{He}]) \\
&\quad + V_{\text{External}} \cdot (\text{kr2unprot} \cdot [\text{R3e}]) \\
\frac{\mathrm{d}([\text{R3e}] \cdot V_{\text{External}})}{\mathrm{d}t} &= -\left(\frac{\text{PermuR} \cdot \text{SurfU} \cdot ([\text{R3e}] - [\text{R3u}])}{\text{vol}_{(\text{DiffR3eu})}}\right) \\
&\quad + V_{\text{External}} \cdot (\text{kr1prot} \cdot [\text{R4e}] \cdot [\text{He}]) \\
&\quad - V_{\text{External}} \cdot (\text{kr2unprot} \cdot [\text{R3e}]) \\
\frac{\mathrm{d}([\text{P3e}] \cdot V_{\text{External}})}{\mathrm{d}t} &= -\left(\frac{\text{PermuP} \cdot \text{SurfU} \cdot ([\text{P3e}] - [\text{P3u}])}{\text{vol}_{(\text{DiffP3eu})}}\right) \\
&\quad - V_{\text{External}} \cdot (\text{kp1prot} \cdot [\text{P3e}] \cdot [\text{He}]) \\
&\quad + V_{\text{External}} \cdot (\text{kp2unprot} \cdot [\text{P2e}]) \\
\frac{\mathrm{d}([\text{P2e}] \cdot V_{\text{External}})}{\mathrm{d}t} &= -\left(\frac{\text{PermuP} \cdot \text{SurfU} \cdot ([\text{P2e}] - [\text{P2u}])}{\text{vol}_{(\text{DiffP2eu})}}\right) \\
&\quad + V_{\text{External}} \cdot (\text{kp1prot} \cdot [\text{P3e}] \cdot [\text{He}]) \\
&\quad - V_{\text{External}} \cdot (\text{kp2unprot} \cdot [\text{P2e}]) \\
\frac{\mathrm{d}([\text{Bu}] \cdot V_{\text{unstirred}})}{\mathrm{d}t} &= +\left(\frac{\text{PermuB} \cdot \text{SurfU} \cdot ([\text{Be}] - [\text{Bu}])}{\text{vol}_{(\text{DiffBeu})}}\right) \\
&\quad - \left(\frac{\text{PermcB} \cdot \text{SurfC} \cdot ([\text{Bu}] - [\text{Bc}])}{\text{vol}_{(\text{DiffBuc})}}\right) \\
&\quad + V_{\text{unstirred}} \cdot (\text{CAu} \cdot \text{CAk1u} \cdot [\text{Cu}]) \\
&\quad - V_{\text{unstirred}} \cdot (\text{CAu} \cdot \text{CAk2u} \cdot [\text{Bu}] \cdot [\text{Hu}]) \\
\frac{\mathrm{d}([\text{Cu}] \cdot V_{\text{unstirred}})}{\mathrm{d}t} &= - V_{\text{unstirred}} \cdot (\text{CAu} \cdot \text{CAk1u} \cdot [\text{Cu}]) \\
&\quad + V_{\text{unstirred}} \cdot (\text{CAu} \cdot \text{CAk2u} \cdot [\text{Bu}] \cdot [\text{Hu}]) \\
&\quad + \left(\frac{\text{PermuC} \cdot \text{SurfU} \cdot ([\text{Ce}] - [\text{Cu}])}{\text{vol}_{(\text{DiffCeu})}}\right) \\
&\quad - \left(\frac{\text{PermcC} \cdot \text{SurfC} \cdot ([\text{Cu}] - [\text{Cc}])}{\text{vol}_{(\text{DiffCuc})}}\right) \\
\frac{\mathrm{d}([\text{Ou}] \cdot V_{\text{unstirred}})}{\mathrm{d}t} &= +\left(\frac{\text{PermuO} \cdot \text{SurfU} \cdot ([\text{Oe}] - [\text{Ou}])}{\text{vol}_{(\text{DiffOeu})}}\right) \\
&\quad - \left(\frac{\text{PermcO} \cdot \text{SurfC} \cdot ([\text{Ou}] - [\text{Oc}])}{\text{vol}_{(\text{DiffOuc})}}\right) \\
\frac{\mathrm{d}([\text{Hu}] \cdot V_{\text{unstirred}})}{\mathrm{d}t} &= +\left(\frac{\text{PermuH} \cdot \text{SurfU} \cdot ([\text{He}] - [\text{Hu}])}{\text{vol}_{(\text{DiffHeu})}}\right)
\end{aligned}
$$

$$
\begin{aligned}
& - \left(\frac{\mathrm{PermcH}\cdot\mathrm{SurfC}\cdot([\mathrm{Hu}]-[\mathrm{Hc}])}{\mathrm{vol}_{(\mathrm{DiffHuc})}}\right)\\
& - V_{\mathrm{unstirred}}\cdot(\mathrm{kr1prot}\cdot[\mathrm{R4u}]\cdot[\mathrm{Hu}])\\
& + V_{\mathrm{unstirred}}\cdot(\mathrm{kr2unprot}\cdot[\mathrm{R3u}])\\
& - V_{\mathrm{unstirred}}\cdot(\mathrm{kp1prot}\cdot[\mathrm{P3u}]\cdot[\mathrm{Hu}])\\
& + V_{\mathrm{unstirred}}\cdot(\mathrm{kp2unprot}\cdot[\mathrm{P2u}])\\
& + V_{\mathrm{unstirred}}\cdot(\mathrm{CAu}\cdot\mathrm{CAk1u}\cdot[\mathrm{Cu}])\\
& - V_{\mathrm{unstirred}}\cdot(\mathrm{CAu}\cdot\mathrm{CAk2u}\cdot[\mathrm{Bu}]\cdot[\mathrm{Hu}])\\
\frac{\mathrm{d}([\mathrm{R4u}]\cdot V_{\mathrm{unstirred}})}{\mathrm{d}t} = & + \left(\frac{\mathrm{PermuR}\cdot\mathrm{SurfU}\cdot([\mathrm{R4e}]-[\mathrm{R4u}])}{\mathrm{vol}_{(\mathrm{DiffR4eu})}}\right)\\
& - \left(\frac{\mathrm{PermcR}\cdot\mathrm{SurfC}\cdot([\mathrm{R4u}]-[\mathrm{R4c}])}{\mathrm{vol}_{(\mathrm{DiffR4uc})}}\right)\\
& - V_{\mathrm{unstirred}}\cdot(\mathrm{kr1prot}\cdot[\mathrm{R4u}]\cdot[\mathrm{Hu}])\\
& + V_{\mathrm{unstirred}}\cdot(\mathrm{kr2unprot}\cdot[\mathrm{R3u}])\\
\frac{\mathrm{d}([\mathrm{R3u}]\cdot V_{\mathrm{unstirred}})}{\mathrm{d}t} = & + \left(\frac{\mathrm{PermuR}\cdot\mathrm{SurfU}\cdot([\mathrm{R3e}]-[\mathrm{R3u}])}{\mathrm{vol}_{(\mathrm{DiffR3eu})}}\right)\\
& - \left(\frac{\mathrm{PermcR}\cdot\mathrm{SurfC}\cdot([\mathrm{R3u}]-[\mathrm{R3c}])}{\mathrm{vol}_{(\mathrm{DiffR3uc})}}\right)\\
& + V_{\mathrm{unstirred}}\cdot(\mathrm{kr1prot}\cdot[\mathrm{R4u}]\cdot[\mathrm{Hu}])\\
& - V_{\mathrm{unstirred}}\cdot(\mathrm{kr2unprot}\cdot[\mathrm{R3u}])\\
\frac{\mathrm{d}([\mathrm{P3u}]\cdot V_{\mathrm{unstirred}})}{\mathrm{d}t} = & + \left(\frac{\mathrm{PermuP}\cdot\mathrm{SurfU}\cdot([\mathrm{P3e}]-[\mathrm{P3u}])}{\mathrm{vol}_{(\mathrm{DiffP3eu})}}\right)\\
& - \left(\frac{\mathrm{PermcP}\cdot\mathrm{SurfU}\cdot([\mathrm{P3u}]-[\mathrm{P3c}])}{\mathrm{vol}_{(\mathrm{DiffP3uc})}}\right)\\
& - V_{\mathrm{unstirred}}\cdot(\mathrm{kp1prot}\cdot[\mathrm{P3u}]\cdot[\mathrm{Hu}])\\
& + V_{\mathrm{unstirred}}\cdot(\mathrm{kp2unprot}\cdot[\mathrm{P2u}])\\
\frac{\mathrm{d}([\mathrm{P2u}]\cdot V_{\mathrm{unstirred}})}{\mathrm{d}t} = & + \left(\frac{\mathrm{PermuP}\cdot\mathrm{SurfU}\cdot([\mathrm{P2e}]-[\mathrm{P2u}])}{\mathrm{vol}_{(\mathrm{DiffP2eu})}}\right)\\
& - \left(\frac{\mathrm{PermcP}\cdot\mathrm{SurfC}\cdot([\mathrm{P2u}]-[\mathrm{P2c}])}{\mathrm{vol}_{(\mathrm{DiffP2uc})}}\right)\\
& + V_{\mathrm{unstirred}}\cdot(\mathrm{kp1prot}\cdot[\mathrm{P3u}]\cdot[\mathrm{Hu}])\\
& - V_{\mathrm{unstirred}}\cdot(\mathrm{kp2unprot}\cdot[\mathrm{P2u}])\\
\frac{\mathrm{d}([\mathrm{Bc}]\cdot V_{\mathrm{condensate}})}{\mathrm{d}t} = & + \left(\frac{\mathrm{PermcB}\cdot\mathrm{SurfC}\cdot([\mathrm{Bu}]-[\mathrm{Bc}])}{\mathrm{vol}_{(\mathrm{DiffBuc})}}\right)\\
& + V_{\mathrm{condensate}}\cdot(\mathrm{CAc}\cdot\mathrm{CAk1c}\cdot[\mathrm{Cc}])\\
& - V_{\mathrm{condensate}}\cdot(\mathrm{CAc}\cdot\mathrm{CAk2c}\cdot[\mathrm{Bc}]\cdot[\mathrm{Hc}])\\
\frac{\mathrm{d}([\mathrm{Cc}]\cdot V_{\mathrm{condensate}})}{\mathrm{d}t} = & - V_{\mathrm{condensate}}\cdot\left(\frac{\frac{[\mathrm{R4c}]}{[\mathrm{R4c}]+\mathrm{Kr}}\cdot\mathrm{Vcarb}\cdot[\mathrm{Cc}]}{\mathrm{Kc}+[\mathrm{Cc}]+\frac{\mathrm{Kc}\cdot[\mathrm{Oc}]}{\mathrm{Ko}}}\right)\\
& - V_{\mathrm{condensate}}\cdot(\mathrm{CAc}\cdot\mathrm{CAk1c}\cdot[\mathrm{Cc}])
\end{aligned}
$$

$$+V_{\text{condensate}} \cdot (\text{CAc} \cdot \text{CAk2c} \cdot [\text{Bc}] \cdot [\text{Hc}])$$

$$+\left(\frac{\text{PermcC} \cdot \text{SurfC} \cdot ([\text{Cu}] - [\text{Cc}])}{\text{vol}_{(\text{DiffCuc})}}\right)$$

$$\frac{\mathrm{d}([\text{Oc}] \cdot V_{\text{condensate}})}{\mathrm{d}t} = -V_{\text{condensate}} \cdot \left(\frac{\frac{[\text{R4c}]}{[\text{R4c}]+\text{Kr}} \cdot \text{Vox} \cdot [\text{Oc}]}{\text{Ko} + [\text{Oc}] + \frac{\text{Ko} \cdot [\text{Cc}]}{\text{Kc}}}\right)$$

$$+\left(\frac{\text{PermcO} \cdot \text{SurfC} \cdot ([\text{Ou}] - [\text{Oc}])}{\text{vol}_{(\text{DiffOuc})}}\right)$$

$$\frac{\mathrm{d}([\text{Hc}] \cdot V_{\text{condensate}})}{\mathrm{d}t} = +2 \cdot V_{\text{condensate}} \cdot \left(\frac{\frac{[\text{R4c}]}{[\text{R4c}]+\text{Kr}} \cdot \text{Vcarb} \cdot [\text{Cc}]}{\text{Kc} + [\text{Cc}] + \frac{\text{Kc} \cdot [\text{Oc}]}{\text{Ko}}}\right)$$

$$+V_{\text{condensate}} \cdot \left(\frac{\frac{[\text{R4c}]}{[\text{R4c}]+\text{Kr}} \cdot \text{Vox} \cdot [\text{Oc}]}{\text{Ko} + [\text{Oc}] + \frac{\text{Ko} \cdot [\text{Cc}]}{\text{Kc}}}\right)$$

$$+\left(\frac{\text{PermcH} \cdot \text{SurfC} \cdot ([\text{Hu}] - [\text{Hc}])}{\text{vol}_{(\text{DiffHuc})}}\right)$$

$$-V_{\text{condensate}} \cdot (\text{kr1prot} \cdot [\text{R4c}] \cdot [\text{Hc}])$$

$$+V_{\text{condensate}} \cdot (\text{kr2unprot} \cdot [\text{R3c}])$$

$$-V_{\text{condensate}} \cdot (\text{kp1prot} \cdot [\text{P3c}] \cdot [\text{Hc}])$$

$$+V_{\text{condensate}} \cdot (\text{kp2unprot} \cdot [\text{P2c}])$$

$$+V_{\text{condensate}} \cdot (\text{CAc} \cdot \text{CAk1c} \cdot [\text{Cc}])$$

$$-V_{\text{condensate}} \cdot (\text{CAc} \cdot \text{CAk2c} \cdot [\text{Bc}] \cdot [\text{Hc}])$$

$$\frac{\mathrm{d}([\text{R4c}] \cdot V_{\text{condensate}})}{\mathrm{d}t} = -V_{\text{condensate}} \cdot \left(\frac{\frac{[\text{R4c}]}{[\text{R4c}]+\text{Kr}} \cdot \text{Vcarb} \cdot [\text{Cc}]}{\text{Kc} + [\text{Cc}] + \frac{\text{Kc} \cdot [\text{Oc}]}{\text{Ko}}}\right)$$

$$-V_{\text{condensate}} \cdot \left(\frac{\frac{[\text{R4c}]}{[\text{R4c}]+\text{Kr}} \cdot \text{Vox} \cdot [\text{Oc}]}{\text{Ko} + [\text{Oc}] + \frac{\text{Ko} \cdot [\text{Cc}]}{\text{Kc}}}\right)$$

$$+\left(\frac{\text{PermcR} \cdot \text{SurfC} \cdot ([\text{R4u}] - [\text{R4c}])}{\text{vol}_{(\text{DiffR4uc})}}\right)$$

$$-V_{\text{condensate}} \cdot (\text{kr1prot} \cdot [\text{R4c}] \cdot [\text{Hc}])$$

$$+V_{\text{condensate}} \cdot (\text{kr2unprot} \cdot [\text{R3c}])$$

$$\frac{\mathrm{d}([\text{R3c}] \cdot V_{\text{condensate}})}{\mathrm{d}t} = +\left(\frac{\text{PermcR} \cdot \text{SurfC} \cdot ([\text{R3u}] - [\text{R3c}])}{\text{vol}_{(\text{DiffR3uc})}}\right)$$

$$+V_{\text{condensate}} \cdot (\text{kr1prot} \cdot [\text{R4c}] \cdot [\text{Hc}])$$

$$-V_{\text{condensate}} \cdot (\text{kr2unprot} \cdot [\text{R3c}])$$

$$\frac{\mathrm{d}([\text{P3c}] \cdot V_{\text{condensate}})}{\mathrm{d}t} = +2 \cdot V_{\text{condensate}} \cdot \left(\frac{\frac{[\text{R4c}]}{[\text{R4c}]+\text{Kr}} \cdot \text{Vcarb} \cdot [\text{Cc}]}{\text{Kc} + [\text{Cc}] + \frac{\text{Kc} \cdot [\text{Oc}]}{\text{Ko}}}\right)$$

$$+V_{\text{condensate}} \cdot \left(\frac{\frac{[\text{R4c}]}{[\text{R4c}]+\text{Kr}} \cdot \text{Vox} \cdot [\text{Oc}]}{\text{Ko} + [\text{Oc}] + \frac{\text{Ko} \cdot [\text{Cc}]}{\text{Kc}}}\right)$$

$$+\left(\frac{\text{PermcP} \cdot \text{SurfU} \cdot ([\text{P3u}] - [\text{P3c}])}{\text{vol}_{(\text{DiffP3uc})}}\right)$$

$$-V_{\text{condensate}} \cdot (\text{kp1prot} \cdot [\text{P3c}] \cdot [\text{Hc}])$$

$$
\begin{aligned}
\frac{\mathrm{d}([\mathrm{P2c}] \cdot V_{\mathrm{condensate}})}{\mathrm{d}t} = \quad & + V_{\mathrm{condensate}} \cdot (\mathrm{kp2unprot} \cdot [\mathrm{P2c}]) \\
& + \left( \frac{\mathrm{PermcP} \cdot \mathrm{SurfC} \cdot ([\mathrm{P2u}] - [\mathrm{P2c}])}{\mathrm{vol}_{(\mathrm{DiffP2uc})}} \right) \\
& + V_{\mathrm{condensate}} \cdot (\mathrm{kp1prot} \cdot [\mathrm{P3c}] \cdot [\mathrm{Hc}]) \\
& - V_{\mathrm{condensate}} \cdot (\mathrm{kp2unprot} \cdot [\mathrm{P2c}])
\end{aligned}
$$

$$
\begin{aligned}
\mathrm{VolC} &= \tfrac{4}{3} \cdot 3.1416 \cdot \mathrm{Values[RadiusC].InitialValue}^3 \\
\mathrm{VolU} &= \tfrac{4}{3} \cdot 3.1416 \cdot \mathrm{Values[RadiusU].InitialValue}^3 - \mathrm{Values[VolC].InitialValue} \\
\mathrm{SurfU} &= \mathrm{Values[Number].InitialValue} \cdot 4 \cdot 3.1416 \cdot \mathrm{Values[RadiusU].InitialValue}^2 \\
\mathrm{SurfC} &= \mathrm{Values[Number].InitialValue} \cdot 4 \cdot 3.1416 \cdot \mathrm{Values[RadiusC].InitialValue}^2 \\
\mathrm{RadiusU} &= 1.5 \cdot \mathrm{Values[RadiusC].InitialValue} \\
\mathrm{Vcarb} &= \mathrm{Values[Vc].InitialValue} \cdot \mathrm{Values[sites].InitialValue} \\
\mathrm{Vox} &= \mathrm{Values[Vo].InitialValue} \cdot \mathrm{Values[sites].InitialValue}
\end{aligned}
$$

### Legend for Dataset S1

This dataset contains: 1) net carboxylation turnover numbers for various Rubisco enzymes (Table 2) over the range 0.01 – 25 mM $HCO_3^-$ at 20% and 30% (*v/v*) $O_2$ in the external compartment (analogous to a fixed cell cytoplasm) in the COPASI model; 2) a set of evolution matrices for each of the Rubisco enzymes where each is modelled as potential intermediate evolutionary states from the free enzyme to contemporary carboxysomes at high and low $CO_2$ and $O_2$; 3) summary matrices in which the catalytic parameters of each Rubisco enzyme are compared to enable correlations between relative enzyme fitness at different potential intermediate evolutionary states from the free enzyme to large contemporary carboxysomes (radius $1 \times 10^{-6}$ m) at high and low $CO_2$ and $O_2$.

For the net carboxylation turnover numbers, these values are the sum of Rubisco carboxylation turnover numbers minus ½ Rubisco oxygenase turnover at each $HCO_3^-$ concentration in the range analyzed. Net rates are presented for 20% and 30% (*v/v*) $O_2$ atmospheres.

Evolution matrix data are calculated as % changes, between potential evolutionary states, in average net carboxylation rates over high or low $CO_2$ ranges at either 20% or 30% (*v/v*) $O_2$ (see Fig. 6). An additional set of evolution matrix data are supplied for the tobacco enzyme modelled at 50 µM RuBP in the model at 20% (*v/v*) $O_2$ as described in Fig. S15.

Summary matrix data for specific evolutionary intermediates are comparisons of net carboxylation rates, at high and low $CO_2$ and $O_2$, for each Rubisco enzyme used in the model. These tables also contain Pearson statistics for correlations between average net carboxylation (under each $CO_2$ range and $O_2$ concentration), and the catalytic parameters of each Rubisco analyzed (Table 2).

### Legend for Dataset S2

This dataset contains: 1) net carboxylation turnover numbers for various Rubisco enzymes (Table 2) over the range 0.01 – 25 mM $HCO_3^-$ at 20% and 30% (*v/v*) $O_2$ in the external compartment (analogous to a fixed cell cytoplasm) in the COPASI model; 2) a set of evolution matrices for each of the Rubisco enzymes where each is modelled as potential intermediate evolutionary states from the free enzyme to contemporary carboxysomes at high and low $CO_2$ and $O_2$; 3) summary matrices in which the catalytic parameters of each Rubisco enzyme are compared to enable correlations between relative enzyme fitness at different potential intermediate evolutionary states from the free enzyme to small contemporary carboxysomes (radius $5 \times 10^{-8}$ m) at high and low $CO_2$ and $O_2$.

For the net carboxylation turnover numbers, these values are the sum of Rubisco carboxylation turnover numbers minus ½ Rubisco oxygenase turnover at each $HCO_3^-$ concentration in the range analyzed. Net rates are presented for 20% and 30% (*v/v*) $O_2$ atmospheres.

Evolution matrix data are calculated as % changes, between potential evolutionary states, in average net carboxylation rates over high or low $CO_2$ ranges at either 20% or 30% (*v/v*) $O_2$ (see Fig. 6). An additional set of evolution matrix data are supplied for the tobacco enzyme modelled at 50 µM RuBP in the model at 20% (*v/v*) $O_2$ as described in Fig. S15.

Summary matrix data for specific evolutionary intermediates are comparisons of net carboxylation rates, at high and low $CO_2$ and $O_2$, for each Rubisco enzyme used in the model. These tables also contain Pearson statistics for correlations between average net carboxylation (under each $CO_2$ range and $O_2$ concentration), and the catalytic parameters of each Rubisco analyzed (Table 2).
